# Adaptive evolution of sesquiterpene deoxyphomenone in mycoparasitism by *Hansfordia pulvinata* associated with horizontal gene transfer from *Aspergillus* species

**DOI:** 10.1101/2024.03.22.586281

**Authors:** Kazuya Maeda, Takuya Sumita, Oumi Nishi, Hirotoshi Sushida, Yumiko Higashi, Hiroyuki Nakagawa, Tomoko Suzuki, Eishin Iwao, Much Zaenal Fanani, Yoshiaki Nishiya, Yuichiro Iida

## Abstract

Leaf mold caused by the ascomycete fungus *Cladosporium fulvum* is a devastating disease of tomato plants. The mycoparasitic fungus *Hansfordia pulvinata* is an effective biocontrol agent that parasitizes *C. fulvum* hyphae on leaves and secretes 13-deoxyphomenone, an eremophilane-type sesquiterpene, which was also identified as a sporulation-inducing factor in *Aspergillus oryzae*. Here, we identified deoxyphomenone biosynthesis (*DPH*) gene clusters conserved in both *H. pulvinata* and *Aspergillus* section *Flavi* including *A. oryzae* and *A. flavus*. Functional disruption of *DPH1* orthologous genes encoding sesquiterpene cyclase in *H. pulvinata*, *A. oryzae* and its close relative *A. flavus* revealed that deoxyphomenone in *H. pulvinata* had exogenic antifungal activity against the host fungus *C. fulvum* and controlled endogenic sporulation in *Aspergillus* species. Deoxyphomenone also inhibited mycelial growth of *C. fulvum* and the non-host tomato pathogen *Pseudocercospora fuligena*. Complete *DPH* clusters, highly similar to those in *H. pulvinata*, were exclusive to *Aspergillus* section *Flavi*, while species in other *Aspergillus* sections contained fragmented *DPH* clusters. A comparative genomics analysis revealed that these *DPH* gene clusters share a common origin and are horizontally transferred across large taxonomic distances from an ancestor of *Aspergillus* to *H. pulvinata*. Our results suggest that, after horizontal transfer, *H. pulvinata* maintained the *DPH* cluster as the inhibitory effect of deoxyphomenone on spore germination and mycelial growth contributed to its mycoparasitism on the host fungus *C. fulvum*.

## INTRODUCTION

The biotrophic filamentous fungus *Cladosporium fulvum* (Cooke) (syn. *Passalora fulva*, *Fulvia fulva*), the causal agent of leaf mold of tomato, is an economic problem in most countries that grow tomatoes (Videira et al., 2017; Mesarich et al., 2023). Following spore germination, hyphae enter the leaf through stomata to absorb nutrients from its host in the apoplast. After a few weeks, conidiophores emerge from the stomata and produce numerous spores on the leaves, which appear as brown lesions. Colonization of tomato leaves by *C. fulvum* is facilitated by numerous effector proteins that are secreted into the apoplast by the fungus (de Wit, 2016). To combat *C. fulvum*, breeders have introduced *Cf* resistance genes, which encode receptor-like proteins with extracytoplasmic leucine-rich repeats, into commercial tomato cultivars from wild relatives (de Wit, 2016; Mesarich, 2023). Tomato *Cf* gene encoded receptor-like proteins recognize corresponding effector proteins of *C. fulvum* as virulence factors in the apoplastic region and trigger a hypersensitive response that rapidly kills plant cells near the infection site, inhibiting the growth of the biotrophic pathogen and preventing its spread (Stergiopoulos and de Wit, 2009). However, the occasional loss or mutation of effector genes allows *C. fulvum* to escape the host immune system, resulting in the emergence of new races (de Wit, 2016). Although single resistance genes have been introduced into many tomato cultivars, overuse of resistant cultivars with only one or a few *Cf* genes has led to the development of *C. fulvum* races that have overcome these resistance genes. So far, 13 races have developed in Japan (Iida et al., 2015; Kubota et al., 2015; Yoshida et al., 2021), and *C. fulvum* has evolved to sequentially overcome multiple *Cf* genes (Rosa et al., 2023). The complex race structure of *C. fulvum* in Japan has enabled the fungus to overcome all *Cf* genes present in commercial cultivars (Iida et al., 2015). A further problem is that *C. fulvum* has developed resistance to several fungicides (Yan et al., 2007; Watanabe et al., 2017), and applications of fungicides have to be reduced due to environmental concerns. Thus, the use of antagonistic microorganisms instead of synthetic fungicides is desired for sustainable control of leaf mold.

We serendipitously discovered white mycelia growing over lesions of leaf mold-infected tomato leaves in a greenhouse and found that disease progression was inhibited on these leaves (**Figure 1A**) (Iida et al., 2018). In further studies, we showed that the fungus parasitizes hyphae of *C. fulvum* and inhibited all known races of this fungus (Iida et al., 2018). The mycoparasitic fungus was identified as *Hansfordia pulvinata* (Berk. & M.A. Curtis) S. Hughes [syn. *Dicyma pulvinata* (Berk. & M.A. Curtis) Arx] (Hughes, 1951; Deighton, 1972; von Arx, 1981, 1982) based on its morphological characteristics and a phylogenetic analysis (Iida et al., 2018). The fungus also has biocontrol activity against other plant parasitic fungi, including *Mycosphaerella berkeleyi*, causing late leaf spot of peanut, *Fusicladium heveae* causing South American leaf blight of rubber, and *Epichloë typhina* causing choke disease of grasses (Peresse and Lepicard, 1980; Mitchell and Taber, 1986; Mitchell et al., 1986; Mello et al., 2008; Alderman et al., 2010); it thus suppresses foliar fungal diseases of different host plants. We also demonstrated that isolate 414-3 of *C. fulvum* suppressed tomato leaf mold most effectively in the greenhouse (Iida et al., 2018). However, the mechanism of this mycoparasitism of *H. pulvinata* against *C. fulvum* is not yet clear.

**Figure 1.**
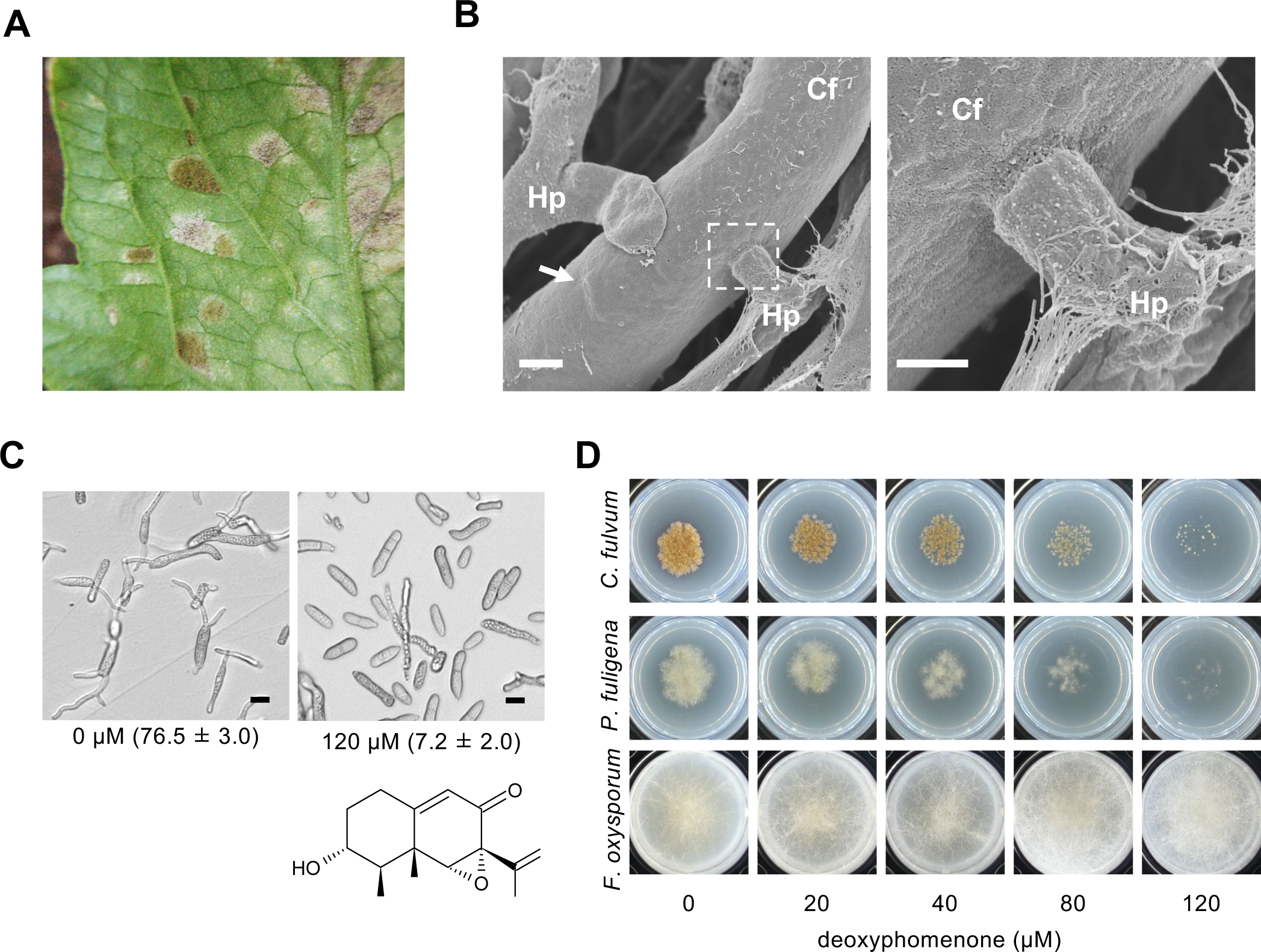
Deoxyphomenone produced by the mycoparasite *Hansfordia pulvinata* is an antifungal sesquiterpene active against the tomato leaf mold pathogen *Cladosporium fulvum*. (**A**) Leaf mold symptoms (brown lesions) caused by *C. fulvum* and lesions covered by white mycelia of *H. pulvinata* on a tomato leaf. (**B**) Scanning electron micrographs of hyphae of *H. pulvinata* parasitizing hyphae of *C. fulvum.* Right: detail of area in dotted box in left image. Arrow: bulge in hypha of *C. fulvum* caused by hyphae of *H. pulvinata*. (**C**) Germinated spores of *C. fulvum* 24 h after treatment with 120 µM deoxyphomenone or with 1% methanol (v/v) as a control (0 µM deoxyphomenone). Germination percentages are in parentheses. Chemical structure of deoxyphomenone is also shown. (**D**) Antifungal activities of various concentrations of deoxyphomenone in MM agar on mycelial growth of tomato pathogenic fungi (*Cladosporium fulvum*, *Pseudocercospora fuligena* and *Fusarium oxysporum* f. sp. *lycopersici*). Bars = 10 μm.

*H. pulvinata* produces 13-deoxyphomenone, an eremophilane-type sesquiterpene, as an antifungal compound during vegetative growth (Tirilly et al., 1983). Around the same time, this compound was also found and characterized (but named sporogen-AO 1) from *Aspergillus oryzae*, which is used to produce koji that is used for the production of many Japanese fermented foods such as Japanese sake and soy sauce (Tanaka et al., 1984a, 1984b). This sesquiterpene is toxic to *C. fulvum*, which is parasitized by *H. pulvinata* but it induces spore production in *A. oryzae*. In addition to its antifungal and sporogenic activities, deoxyphomenone is also phytotoxic (Tansakul et al., 2014; Del Valle et al., 2015), antimalarial (Daengrot et al., 2015), and cytotoxic to cancer cell lines (Tansakul et al., 2017; Liu et al., 2021). However, its biosynthetic pathway in these fungi, the molecular basis of its antifungal activity and spectrum, and the specificity its sporogenic activity in different *Aspergillus* species have not been elucidated.

Fungal secondary metabolites play important roles in development, pathogenicity, protection from biotic and abiotic stresses, antagonism, and communication with other organisms (Keller, 2019). Because deoxyphomenone is produced by most strains of *H. pulvinata* (Iida et al., 2018) and might be widely produced among *Aspergillus* species, it may have mycoparasitic and/or ecological functions in these fungi. In this study, we investigated the roles of deoxyphomenone in *H. pulvinata* and *Aspergillus* species. The antifungal spectrum of the compound was determined for *C. fulvum* and other tomato fungal pathogens that are not host fungi of *H. pulvinata*, such as *Pseudocercospora fuligena*, the causal agent of tomato black leaf mold, which is phylogenetically closely related to *C. fulvum* and the soilborne tomato pathogen *Fusarium oxysporum* f. sp. *lycopersici*. We identified deoxyphomenone biosynthetic gene clusters in *H. pulvinata* and *Aspergillus* genome sequences and functionally disrupted sesquiterpene cyclase genes to reveal the main function of deoxyphomenone. Furthermore, we used a comparative genomics approach to elucidate the origin of these gene clusters in *Aspergillus* and *H. pulvinata*.

## RESULTS

### Antifungal activity of deoxyphomenone against plant pathogens

*H. pulvinata* 414-3, the most effective biocontrol strain with a published draft genomic sequence (Sushida et al. 2019) and the highest deoxyphomenone production among several strains we tested, parasitizes *C. fulvum* hyphae in liquid culture without a carbon source (Iida et al., 2018). In our present tests, *H. pulvinata* did not parasitize *C. fulvum* when grown on nylon membranes on nutrient-rich potato dextrose agar (PDA). However, when the membranes were transferred to nutrient-poor water agar, scanning electron micrographs showed that hyphae of strain 414-3 penetrated the cell wall of *C. fulvum* (**Figure 1B**). Although we previously reported that hyphae of *H. pulvinata* coil around the host hyphae, it does not always do so, and it does not form appressorium-like structures, both common attributes of the mycoparasitic *Trichoderma* fungi.

Since strain 414-3 produces about 100 µM deoxyphomenone in minimal medium (MM) broth (Iida et al., 2018), we next evaluated its antifungal activity against *C. fulvum* using 120 µM commercial deoxyphomenone. At 120 µM, the compound inhibited colony formation and hyphal elongation by more than 90% on MM agar (**Supplementary figure 1A, 1B**). Deoxyphomenone also strongly inhibited spore germination at 120 µM, which was only 7.2% after 24 h compared with 76.5% for the control spores treated with 1% (v/v) methanol (**Figure 1C and Supplementary figure 1C**). When *C. fulvum* spores were treated with the fungicide captan at 100 µM and after 24 h captan was replaced by distilled water, almost all failed to germinate, but when deoxyphomenone (120 µM) was replaced by distilled water 24 h after treatment, about half of the treated spores did germinate (**Supplementary figure S1C**). Spores treated continuously with deoxyphomenone or captan for 48 h failed to germinate. Thus, the activity of deoxyphomenone is fungistatic; it inhibits the germination of *C. fulvum* spores but does not kill them. Deoxyphomenone treatment did not affect the structure of intracellular organelles or the cell membrane of the spores or induce necrosis in tomato leaves even at concentrations as high as 80 µM, contrary to a previous report (Tirilly et al., 1983) (**Supplementary figure S1D, E**). When plated on MM agar, external application of deoxyphomenone also inhibited colony formation of plant pathogenic fungi that are not hosts of *H. pulvinata*: the growth of *P. fuligena* phylogenetically closely related to *C. fulvum*, was inhibited at the same concentrations as used for *C. fulvum*, but growth of *F. oxysporum* f. sp. *lycopersici* was not affected (**Figure 1D**).

### Deoxyphomenone biosynthetic gene clusters are conserved between *H. pulvinata* and *Aspergillus*

To identify the genes involved in deoxyphomenone biosynthesis in the genome sequences of *H. pulvinata* and *A. oryzae*, we searched for homologous genes using the amino acid sequence of aristolocene synthase Ari1 from *Penicillium roqueforti* (Proctor and Hohn, 1993), since the structure of deoxyphomenone has an aristolochene-like backbone (**Figure 1C**), and found two candidates (gene IDs, DIP_001954 and DIP_006271) in *H. pulvinata* 414-3 (Sushida et al. 2019) and one (GenBank accession XP_023093357) in *A. oryzae* RIB40. Phylogenetic comparisons with other fungal sesquiterpene cyclases placed DIP_001954 and XP_023093357 in the same clade as enzymes that catalyze the 1,10-cyclization of farnesyl diphosphate, a common sesquiterpene precursor (**Supplementary figure S2**). The deduced amino acid sequences of DIP_001954 and XP_023093357 had 61.6% similarity (49.1% identity) (**Supplementary figure S3**). We named DIP_001954 “*HpDPH1*” (<u>d</u>eoxy<u>ph</u>omenone biosynthesis gene 1) and the homologous gene XP_023093357 “*AoDPH1*”.

In the nucleotide sequences around *HpDPH1* we found genes encoding short-chain dehydrogenase (*HpDPH2*), three cytochrome P450s (*HpDPH3, HpDPH4, HpDPH5*), and major facilitator superfamily transporter (*HpDPH6*), altogether forming the deoxyphomenone biosynthesis (*DPH*) gene cluster. We also found homologs of these genes in *A. oryzae* (**Figure 2A**). Also genes encoding a lipase/thioesterase family protein and an integral membrane protein were conserved in both fungi, but their function in deoxyphomenone biosynthesis is unknown. The position and orientation of all genes were highly conserved, but the loci of *AoDPH4* and *AoDPH5* were switched (**Supplementary figure S4**). Comparison of these genes revealed that the number of exons in the pairs *HpDPH1/AoDPH1* and *HpDPH5/AoDPH5* was identical and that the sequence similarity in each exon was conserved among all pairs (**Supplementary figure S5**). We presumed that the deoxyphomenone biosynthetic pathway involved cyclization of farnesyl diphosphate by HpDph1, followed by oxygenation and epoxidation by cytochrome P450s HpDph3, HpDph4 and HpDph5, and subsequently hydroxylated by HpDph2 (**Figure 2B**). The putative substrate binding pocket that is conserved in the major facilitator superfamily transporter (Quistgaard et al., 2016) was also conserved in HpDph6/AoDph6 (**Supplementary figure S6**).

**Figure 2.**
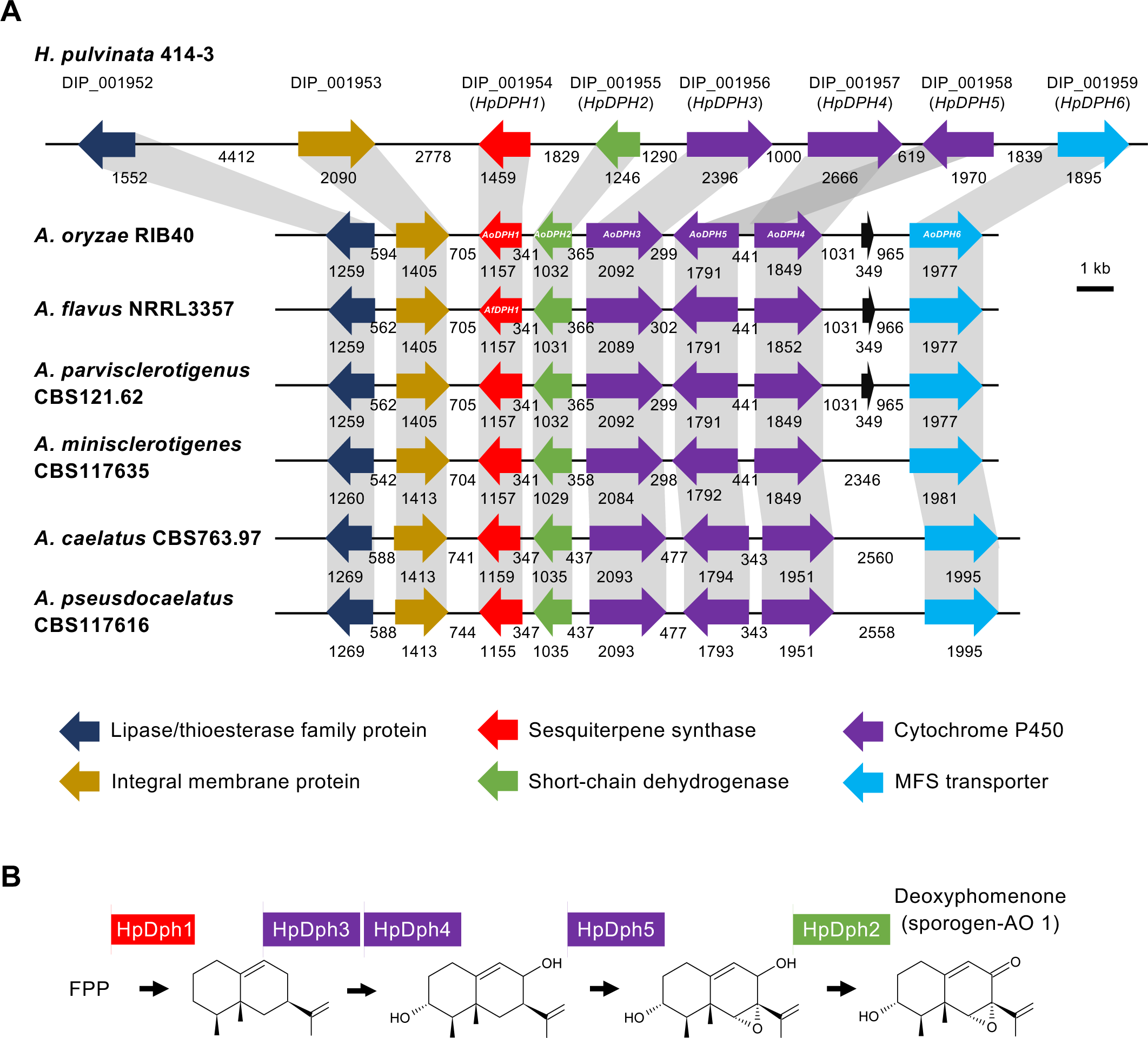
Synteny of the deoxyphomenone biosynthetic (*DPH*) gene clusters conserved in the mycoparasite *Hansfordia pulvinata* and *Aspergillus* section *Flavi*. (**A**) Gene clusters conserved in *H. pulvinata* and *Aspergillus*. The original gene IDs (DIP) and names (*HpDPH1* to *HpDHP6*) of *H. pulvinata* are above the genes (detailed in Supplementary Table S1). Protein annotations are indicated by color. The names of *HpDPH* homologous genes in *Aspergillus* mentioned in the text are indicated in the arrows. Numbers indicate the length of the genes and UTRs in base pairs. ORFs (black arrows) were detected in the upstream region of *HpDPH6* in three *Aspergillus* species, and no homologous gene sequence was found in the NCBI non-redundant database. (**B**) Predicted biosynthetic pathway of deoxyphomenone. Intermediates were logically inferred. FPP, farnesyl pyrophosphate.

To obtain a comprehensive view of the distribution of the *DPH* cluster in other fungi, we extended the search to *HpDPH1* homologues in the MycoCosm database (mycocosm.jgi.doe.gov; Grigoriev, 2014). Genes partially homologous to *HpDPH1* were found in 139 fungal genomes of the *Eurotiomycetes*, *Sordariomycetes* and *Dothideomycetes* (**Supplementary table S1**). Besides the *DPH* gene cluster present in *H. pulvinata*, other *DPH* genes were detected as a gene cluster in *Aspergillus* species only (*Eurotiomycetes*), but not in the *Xylariales* (*Sordariomycetes*), to which *H. pulvinata* belongs, which suggests that the latter fungus has obtained this cluster by horizontal transfer. Notably, the cluster was completely conserved in *Aspergillus* species section *Flavi* to which *A. oryzae* belongs (**Supplementary figure S7 and Figure 2A**).

We then analyzed expression of the putative deoxyphomenone biosynthetic genes during mycoparasitism of *H. pulvinata* 414-3 *in vitro* using quantitative real-time PCR. *H. pulvinata* never parasitize *C. fulvum* on the nutrient-rich PDA, but it did so on the nutrient-poor water agar (**Figure 2A**). The expression of the *HpDPH* genes in *H. pulvinata* did not differ when grown on PDA medium or on water agar (**Figure 3**). In contrast, the expression of *HpDPH* genes was significantly upregulated when *H. pulvinata* parasitized *C. fulvum* on water agar.

**Figure 3.**
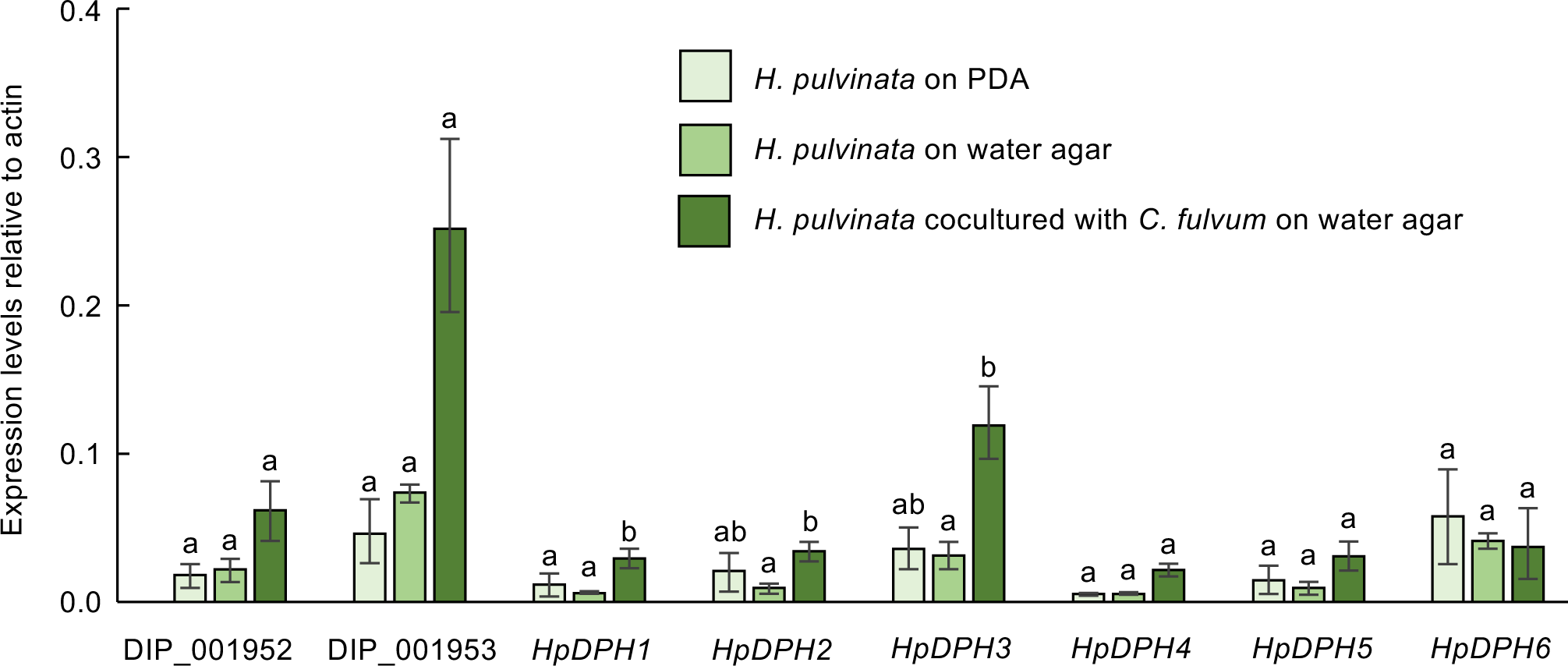
Relative gene expression levels of the deoxyphomenone biosyntheticsis (*DPH*) genes in the mycoparasite *Hansfordia pulvinata*. A spore suspension of *H. pulvinata* 414-3 was sprayed on *Cladosporium fulvum* on water agar (WA). *H. pulvinata* and *C. fulvum* were grown on PDA or WA as controls. Expression was quantified by qPCR and normalized using those of *H. pulvinata* actin gene. Values are the means of three biological replicates (± SD). Significant differences (*P* < 0.05) in expression for a gene among the three treatments are indicated by different letters.

Expression of *HpDPH1*, *HpDPH2* and *HpDPH3* genes, which encode sesquiterpene cyclase, short-chain dehydrogenase, and cytochrome P450, respectively, were significantly induced during mycoparasitism (**Figure 3**). Expression of DIP_001952 (lipase/thioesterase family protein) and DIP_001953 (integral membrane protein) genes also increased, suggesting that these proteins are associated with deoxyphomenone biosynthesis or mycoparasitism.

### Deoxyphomenone is the main antifungal compound of *H. pulvinata* effective against *C. fulvum*

Since the *DPH* gene cluster was also found in the genus *Aspergillus*, we assayed for the presence of deoxyphomenone in the MM culture filtrate from 22 strains of *A. oryzae* and 12 strains of *A. flavus*. In contrast to *H. pulvinata* 414-3, which produced deoxyphomenone as high as about 80 µM, *A. oryzae* and *A. flavus* strains produced only low levels of deoxyphomenone: the highest concentrations in *A. oryzae* and *A. flavus* strains are about 0.8 µM and about 3.5 µM, respectively (**Supplementary figure S8**). We then measured deoxyphomenone by the most productive strains of *A. oryzae* (RIB40) and *A. flavus* (NBRC114564) in different culturing conditions and found that both strains produced the compound when grown as still cultures in MM broth (**Supplementary figure S9**). Interestingly, neither strain produced deoxyphomenone at 35 °C, the optimum temperature for vegetative growth and sporulation, but showed maximal production at 25 °C, at which vegetative growth and sporulation are retarded. Similarly, *H. pulvinata* showed the highest production of deoxyphomenone at 28 °C, a temperature at which vegetative growth is also retarded (**Supplementary figure S9**).

To determine the function of *HpDPH1* in deoxyphomenone biosynthesis in *H. pulvinata* 414-3, we generated Δ*HpDPH1* knockout mutant strains of 414-3 by homologous recombination through *Agrobacterium tumefaciens*-mediated transformation (**Supplementary figure S10**). Quantitative analysis by LC-MS/MS revealed the presence of deoxyphomenone in the culture filtrate of wild-type 414-3 strain, but none in that of the *ΔHpDPH1* mutant strain KO10 (**Figure 4A**). Strain 414-3 started to produce deoxyphomenone at 3 d and increased until 15 d, whereas strain KO10 did not produce any (**Figure 4B**). The culture filtrate of 14-day-old cultures of 414-3 containing deoxyphomenone strongly inhibited the germination of *C. fulvum* spores, whereas those of the Δ*HpDPH1* mutant strains KO10 and KO37 showed no inhibitory activity, suggesting that deoxyphomenone is the main compound contributing to the antifungal activity of *H. pulvinata* (**Figure 4C**). The antifungal activity was partially restored in the loss-of-function Δ*HpDPH1* mutant strain (CO10A) after introduction of a functional *AoDPH1* gene (**Figure 4C**). This is most likely explained by restoration of deoxyphomenone production that accumulated in the culture filtrate of CO10A. The Δ*HpDPH1* mutant KO10 parasitized *C. fulvum* similar to the wild-type 414-3 strain, with no differences in spore number or vegetative growth, as we expected previously (Iida et al., 2018) (**Supplementary figure S11**).

**Figure 4.**
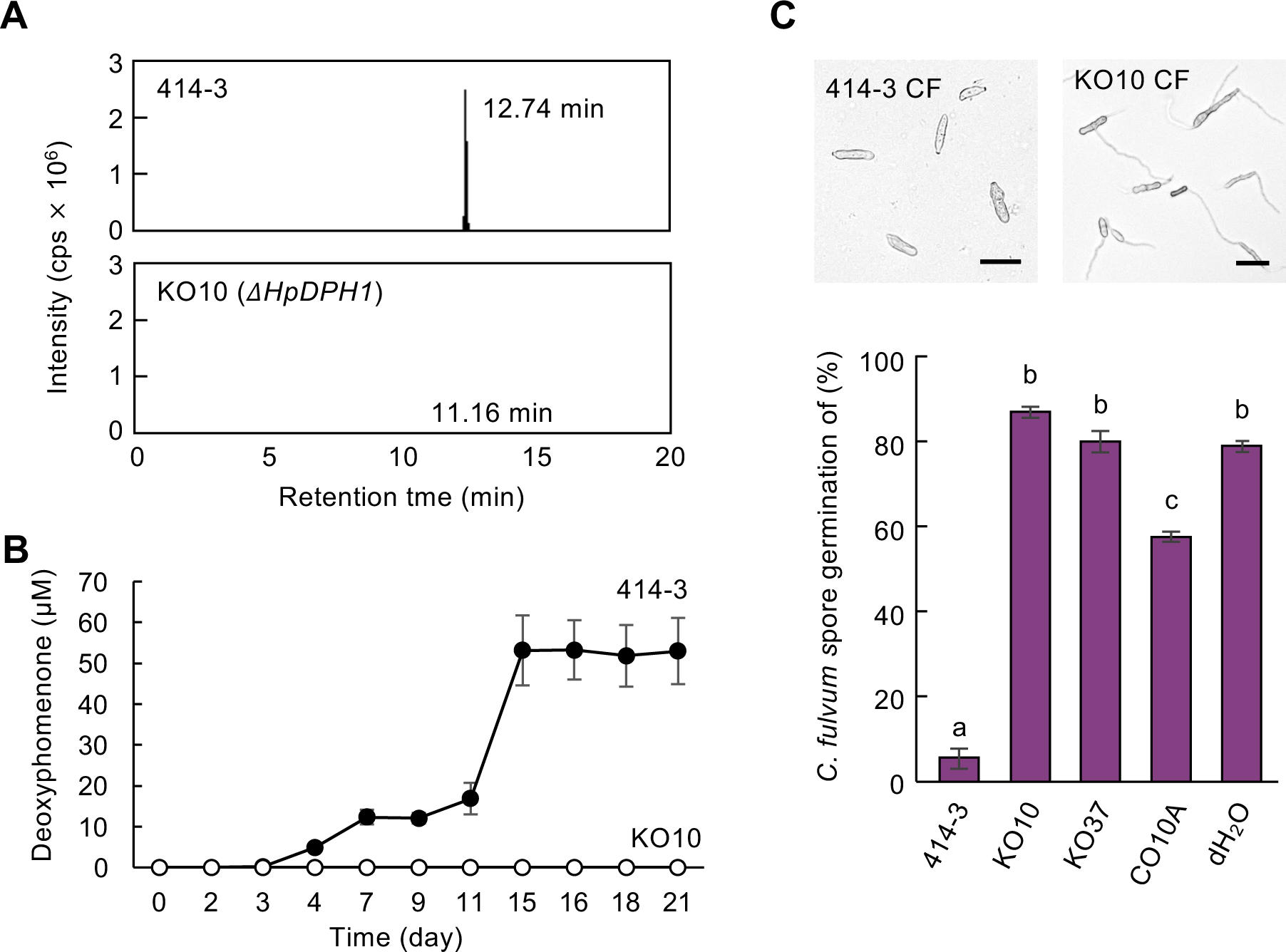
Deoxyphomenone production by wild-type and knock-out mutants of *Hansfordia pulvinata* and their antifungal activity. Wild-type 414-3 of *H. pulvinata*, *ΔHpDPH1* mutants (KO10, KO37) and *AoDHP1-* complemented KO37 (CO10A) were cultured in MM broth for 2 weeks, and the filtered culture filtrate collected to (**A**) quantify deoxyphomenone production in 414-3 and mutant KO10 using LC-MS/MS. The retention time of the highest peak is given for each. (**B**) Time course of deoxyphomenone production in 414-3 and mutant KO10. (**C**) Germination of *Cladosporium fulvum* spores after 24 h of exposure to deoxyphomenone in culture filtrate (CF) of 414-3 or KO10 (bars = 20 μm). Different letters in graph indicate significant differences (*P* < 0.05) in germination among treatments. Values are means of three replicates (± SD).

### The function of deoxyphomenone differs between *H. pulvinata* and the *Aspergillus* species

We also generated knockout strains of the homologous gene *AfDPH1* in *A. flavus* NBRC114564, which produced more deoxyhomenone than *A. oryze* RIB40 (**Supplementary figure S8**). LC-MS/MS analysis showed that wild-type strain and ectopic transformants produced deoxyphomenone, while the Δ*AfDPH1* mutant strains produced none (**Figure 5A**). As mentioned earlier, deoxyphomenone (sporogen-AO 1) was initially identified as a sporogenic factor in *A. oryzae* (Tanaka et al., 1984b, 1984a). Three Δ*AfDPH1* mutant strains grown on MM agar produced significantly less spores than the wild-type NBRC114564 (**Figure 5B, C**). Two ectopic transformants produced similar quantities of spores as the wild-type. Conversely, a Δ*AoDPH1* mutant of *A. oryzae* RIB40 completely lost the ability to produce deoxyphomenone, but its spore production and colony morphology were similar to those of the wild-type and ectopic transformants (**Supplementary figure S12**).

**Figure 5.**
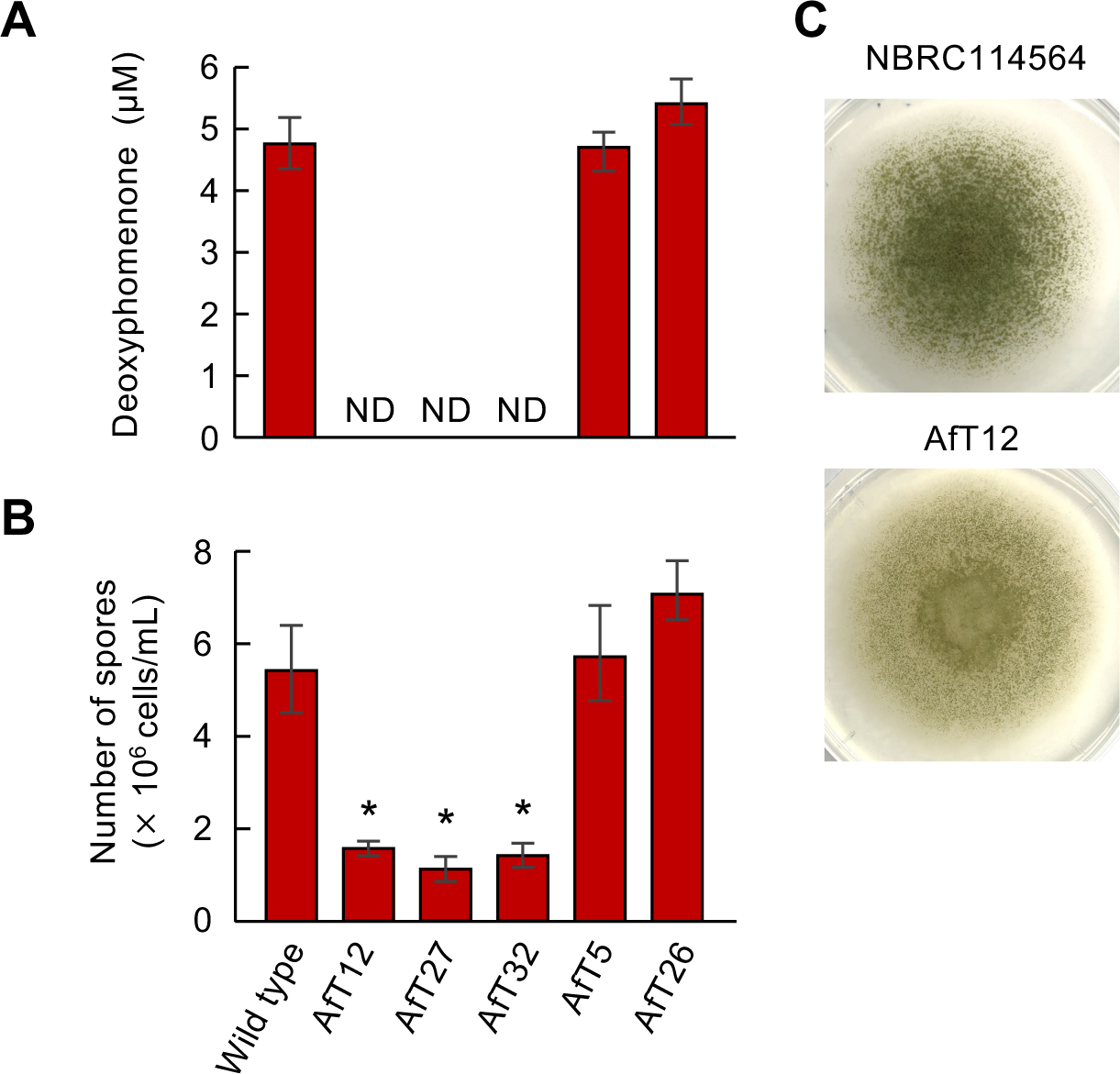
Deoxyphomenone production and sporulation by *A. flavus* wild-type NBRC114564 and Δ*AfDPH1* mutant strains and ectopic mutants. The wild-type, three Δ*AfDPH1* mutants, and two ectopic mutant strains were cultured in MM broth or agar. Values are means of three replicates (± SD). (**A**) LC-MS/MS analysis of deoxyphomenone in culture filtrates of the various strains. ND: not detected. (**B**) Number of spores formed on agar. Asterisk indicates significant difference (*P* < 0.05) compared to the wild-type in Williams’ test. (**C**) Colony morphology of the wild-type, Δ*AfDPH1* mutant and ectopic AfT32 mutant.

We then studied the phenotype of the fungal strains after exposure to deoxyphomenone on MM. Since *H. pulvinata* 414-3 yielded about 30–100 μM deoxyphomenone during vegetative growth, we confirmed that deoxyphomenone was fungistatic against *C. fulvum* over a wide range of concentrations; inhibition by deoxyphomenone was concentration-dependent; inhibition of spore germination of *C. fulvum* was around 25% at 10 μM, reaching about 90% at 80 μM (**Figure 6A**). Thus, the concentration of deoxyphomenone secreted by strain 414-3 seems sufficient to inhibit *C. fulvum* spore germination. As Tanaka et al. (1984b) treated *A. oryzae* strains with higher concentrations of deoxyphomenone (sporogen-AO 1) to test sporogenic activity, we also tested concentrations up to 120 μM against *A. oryzae* RIB40 and *A. flavus* NBRC114564. In *Aspergillus*, deoxyphomenone treatment also affected sporulation in a concentration-dependent manner, but the effect varied among different species: compared to no treatment, treatment with 120 µM deoxyphomenone reduced spore production by RIB40, but increased it in NBRC114564 by ∼90% (**Figure 6B**). While deoxyphomenone did not affect colony growth or sporulation of *H. pulvinata* strain 414-3, colony morphology and formation of aerial hyphae were altered in *Aspergillus* strains, and colony growth was also slightly inhibited in RIB40, indicating that RIB40 is less tolerant to this sesquiterpene than 414-3 (**Figure 6B and Supplementary figure S13**). These results show that deoxyphomenone has quite different activities in different fungi; it has exogenic antifungal activity against the host fungus and affects endogenic sporogenesis in *Aspergillus* species.

**Figure 6.**
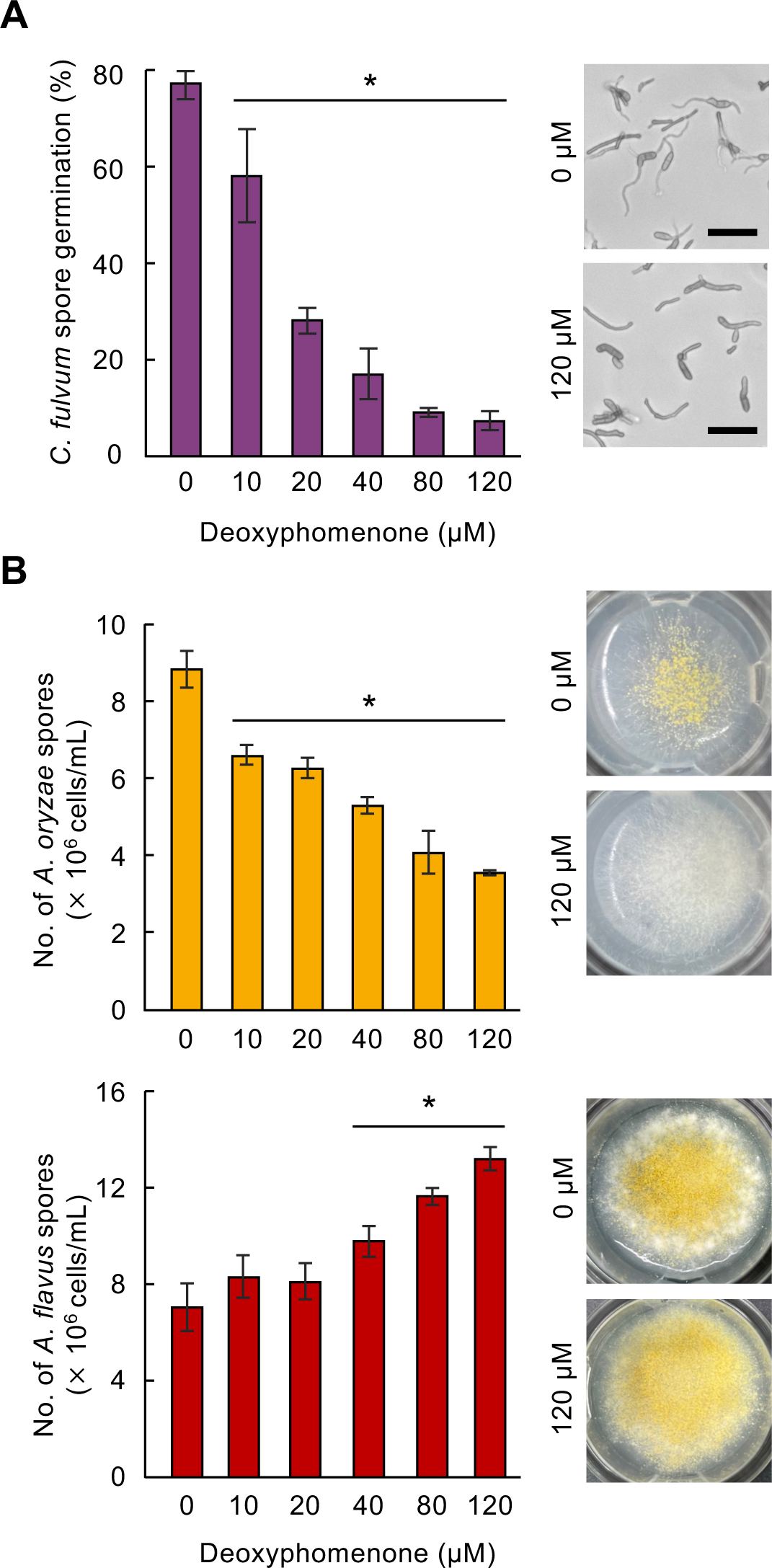
Different effects of deoxyphomenone on spore germination of *Cladosporium fulvum* and spore production of two *Aspergillus* species. Wild-types *C. fulvum* CF301, *A. oryzae* RIB40 and *A. flavus* NBRC114564 were grown on MM agar supplemented with deoxyphomenone. (**A**) Inhibition of spore germination of *Cladosporium fulvum*. (**B**) Effects on sporulation of RIB40 and NBRC114564. Values are the mean of three biological replicates. Error bars indicate standard deviations. Asterisks indicate significant difference (*p* < 0.025) compared to control treatment (0 μM) by Williams’ test.

### Deoxyphomenone biosynthetic gene cluster was horizontally transferred from an *Aspergillus* ancestor to *H. pulvinata*

Next, we analyzed the commonality and conservation of the *DPH* clusters in genomic sequences of *Aspergillus* species. Complete *DPH* clusters were detected only in *Aspergillus* species from the section *Flavi* (Group I, **Figure 2A**), and the protein sequences of HpDph2 to HpDph6 in these species were highly conserved, with ∼70% pairwise similarity to those of *H. pulvinata* (**Figure 7**). Meanwhile, partial *DPH* clusters were present in *Aspergillus* species of the section *Circumdati* (Group II: *HpDPH1*, *HpDPH2*, *HpDPH3*) and the sections *Nidulantes* and *Clavati* (Group III: *HpDPH1*, *HpDPH2*, *HpDPH4*) (**Figure 7**). Group II species also contained a lipase/thioesterase family protein- and an integral membrane protein-encoding genes downstream of *HpDPH1*, the same as in Group I (**Figure 7 and Supplementary figure S7**). Group III further lacked the commonality and conservation of the *DPH* clusters: the similarities with *HpDPH1* and *HpDPH2* genes were much lower than in the other two groups, the position and the order of *DPH2* and *DPH4* genes in the genome sequence were quite different, and unrelated genes were interspersed in the cluster (**Figure 7** and **Supplementary figure S7**). These results suggested that Group II and III species lost parts of the *DPH* cluster and no longer produced deoxyphomenone.

**Figure 7.**
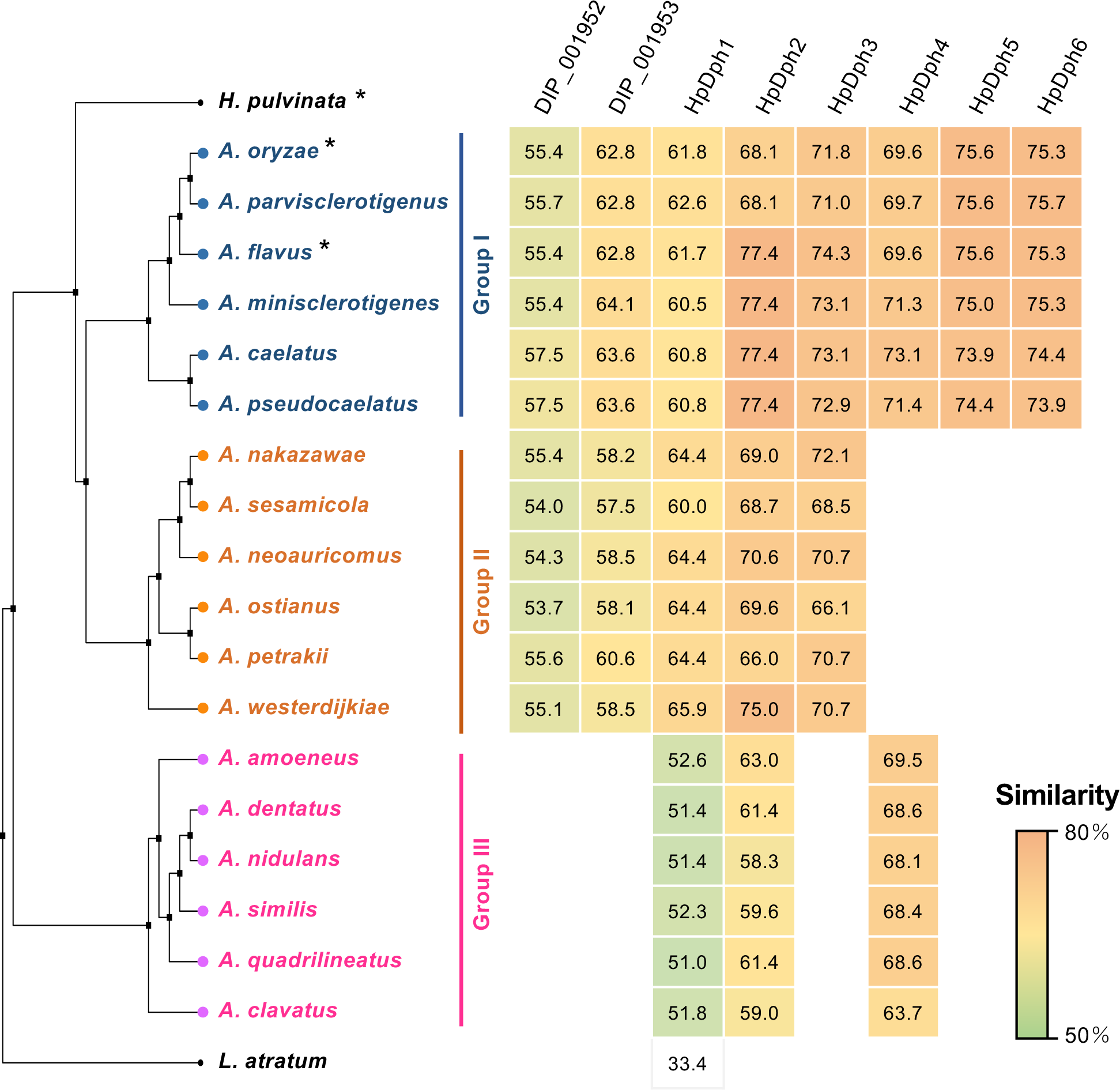
Phylogeny and sequence similarity of deoxyphomenone biosynthetic (*DPH*) genes in *Hansfordia pulvinata* and *Aspergillus* species. The phylogenetic species tree based on the fungal genomic sequences (approximately 15 kb) listed in Supplementary Table S1 was constructed using maximum likelihood. Asterisks represent species that have been verified to produce deoxyphomenone. Sequence of *Lyophyllum atratum* belonging to *Agaricomycetes* was used as an outgroup. *Aspergillus* species were categorized into three groups based on the presence of *DPH* genes. The heatmap represents the pairwise similarity (%) of the deduced amino acid sequences of the *DPH* homologous genes.

To infer the evolutionary origin of the *DPH* clusters in *H. pulvinata* and *Aspergillus*, we analyzed the evolutionary divergence of the clusters among 60 fungal genomes using genomic and phylogenetic approaches with the NOTUNG algorithm, which reconciles differences between species trees and gene trees by inferring gene duplications, transfers, and losses (Stolzer et al., 2012). Reconciliation of the species tree with the HpDph1 protein tree detected a transfer event (T1 in **Figure 8**) early in *Aspergillus* section *Circumdati*, indicating a transfer of *HpDPH1* of *H. pulvinata* from Group II (*Circumdati*) (**Supplementary figure S14**). Likewise, the HpDph2 and HpDph4 protein trees indicated transfers from Group I (section *Flavi*) to *H. pulvinata* (T2 in **Figure 8**). Since the *DPH* clusters are highly conserved in *H. pulvinata* and *Aspergillus*, these results suggest an evolutionary scenario in which the *DPH* cluster was transferred from the common ancestor of Groups I and II (sections *Flavi* and *Circumdati*) to *H. pulvinata* at the same time and that genes were lost in time in *Aspergillus* species (**Figure 8**). Since all duplication events detected in *DPH* genes occurred after the horizontal transfer events (**Supplementary figure S14**), it is unlikely that the duplication is associated with the transfer of the *DPH* cluster. Altogether these results suggest that the *DPH* cluster arose before speciation within the *Aspergillus* genus. Finally, because we inferred the horizontal transfer of the *DPH* cluster from *Aspergillus* to *H. pulvinata*, we investigated the association of deoxyphomenone with the mycoparasitism. Contrary to our expectations, *H. pulvinata* 414-3 did not directly penetrate the hyphae of *A. oryzae* RIB40, as observed with the host fungus *C. fulvum*, but physical contact such as adhesion to or coiling around the hyphae of the RIB40 was detected (**Supplementary figure S15**).

**Figure 8.**
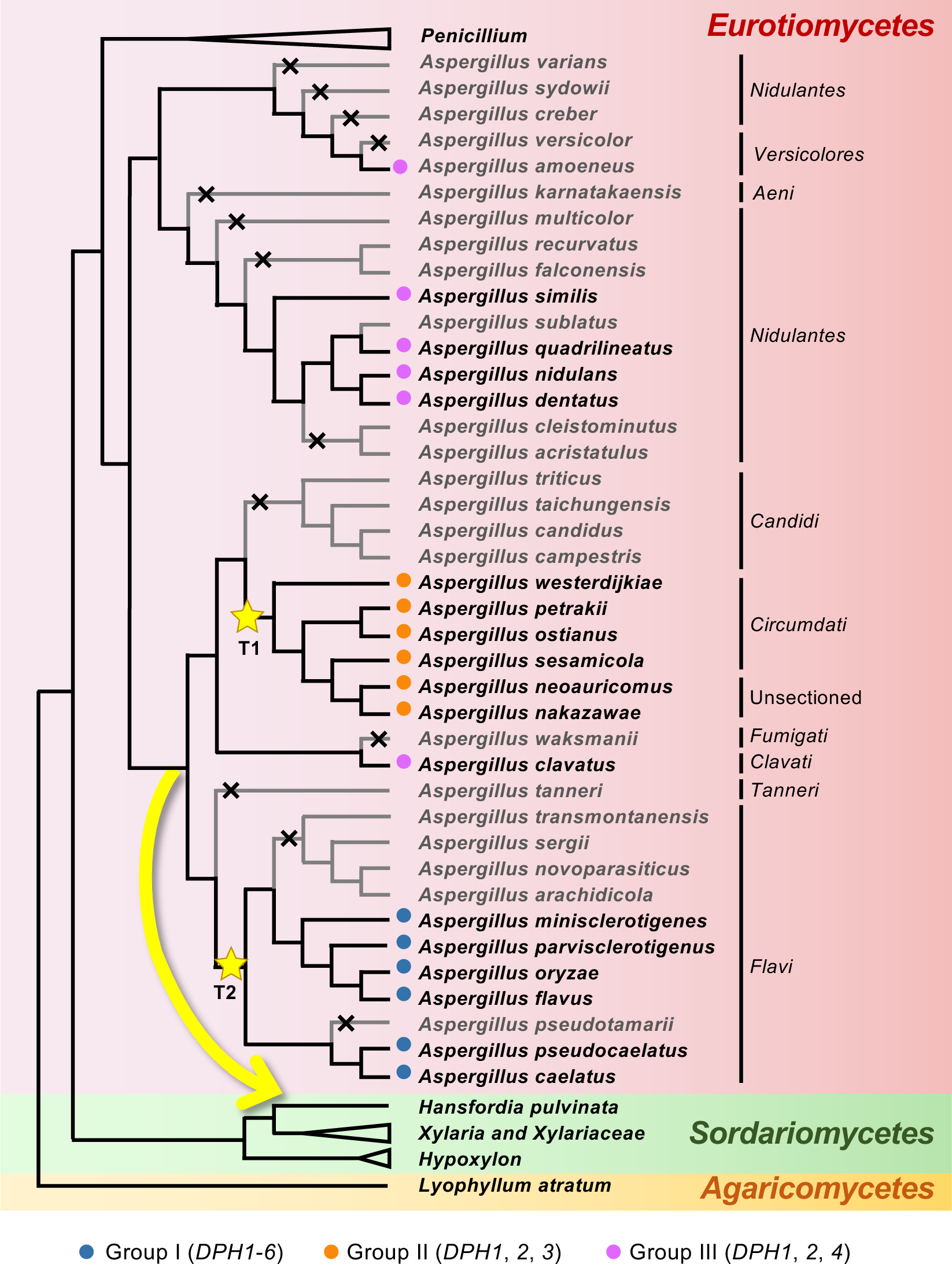
Proposed horizontal transfer of the deoxyphomenone biosynthetic (*DPH*) gene cluster from ancestral *Aspergillus* species to the mycoparasite *Hansfordia pulvinata*. The phylogenetic species tree was based on 500 monocore genes (a single homolog in each of the species) from 60 fungal genomes and constructed using maximum likelihood, and horizontal gene transfer, duplication and loss events inferred in Supplementary figure S14 were described. Horizontal gene transfer events for the *HpDPH1* (T1) and *DPH2/DPH4* (T2) genes are indicated by yellow stars. Proposed horizontal transfer of the *DPH* cluster from ancestral *Aspergillus* to *H. pulvinata* is highlighted by a yellow arrow. Gene loss is marked by an x on the branch. The three *Aspergillus* groups classified by the presence of *DPH* genes shown are the same as in Figure 7. Fungal species with either a complete or partial *DPH* cluster are in black, and those without the cluster are in grey. *Aspergillus* sections are given on the right. *Lyophyllum atratum* was used as an outgroup.

## DISCUSSION

Strict regulation of expression through the physical linkage of genes involved in the same biosynthetic pathway leads to physiological economization; thus, biosynthetic genes for secondary metabolites in fungi are often maintained in clusters (Hagee et al., 2020), which allows for the rapid conversion of potentially toxic or chemically unstable intermediate products (McGary et al., 2013). In the present study, the *DPH* clusters were highly conserved between *H. pulvinata* and *Aspergillus* section *Flavi*, suggesting that the final product, deoxyphomenone, plays critical roles in distinct stages of the life cycle of these fungi. Indeed, it acts as an antifungal agent of the mycoparasite *H. pulvinata* against the host fungus *C. fulvum* and controls sporulation at least in *A. flavus*. The correlation of these fungal clusters can be explained through the most parsimonious evolutionary scenario, which is constructed by reconciling the species tree with the *DPH* protein tree in consideration of the costs of evolutionary events (i.e., gene duplications, transfers, and losses) (Stolzer et al., 2012). Interestingly, we inferred that the *DPH* cluster was horizontally transferred across a large taxonomic distance from the genus *Aspergillus* (*Eurotiomycetes*) to *H. pulvinata* (*Sordariomycetes*), suggesting that *H. pulvinata* maintained the cluster as deoxyphomenone production seemed to be more conductive to mycoparasitism. Horizontal gene transfer is not rare in fungi (Mehrabi et al. 2011; Rokas et al., 2018); gene clusters for the biosynthesis of products such as fumonisin (Proctor et al., 2013, Khaldi and Wolfe, 2011), host-specific toxins in *Alternaria* species (Tsuge et al., 2016), and cercosporin (de Jonge et al., 2018) have been horizontally transferred across classes, but the final products function in the recipient fungi basically in a similar way.

We deduced the deoxyphomenone biosynthetic pathway from the component genes of the *DPH* cluster in *H. pulvinata* and *Aspergillus*. The bicyclic eremophilane-type sesquiterpene deoxyphomenone is probably formed by the cyclization of farnesyl pyrophosphate catalyzed by sesquiterpene cyclase (Proctor and Hohn, 1993), followed by oxygenation and epoxidation of the double bond in *H. pulvinata* and *Aspergillus*. Although deoxyphomenone has also been detected in *Penicillium* species, which are taxonomically close to *Aspergillus*, the *DPH* cluster was not detected in the published genome sequences of *Penicillium* species; thus, the pathway of deoxyphomenone biosynthesis is likely different between the two genera. On the basis of a relaxed molecular clock analysis, *Aspergillus* and *Penicillium* likely diverged 94.0 million years ago (mya) with *Aspergillus* sections *Flavi* and *Circumdati* originating 51.1 mya (Barelli et al., 2015; Steenwyk et al., 2019). Because the *DPH* clusters are fully or partially conserved in *Aspergillus* sections *Flavi* and *Circumdati,* we presume that their horizontal transfer to *H. pulvinata* occurred at least before the common ancestor of these sections.

It is still unclear how *H. pulvinata* acquired the *DPH* clusters from the ancestral *Aspergillus*. Since horizontal transfer of foreign DNA first requires contact between fungal cells, the involvement of anastomosis between conidia, germ tubes or hyphae, and subsequent heterokaryon formation, have been proposed (Kubicek et al., 2019; Fleißner et al., 2022; Bucknell and McDonald, 2023). *H. pulvinata* has a wide host range, including plant pathogens (Peresse and Lepicard, 1980; Mitchell and Taber, 1986; Mitchell et al., 1986; Mitchell et al., 1987; Mello et al., 2008; Alderman et al., 2010), and plant pathogens in *Aspergillus* sections *Flavi* and *Circumdati* may provide opportunities for contact with *H. pulvinata* on plants and subsequent gene transfer. Given the conservation of *DPH* clusters revealed in this study and the nature of the recipient fungus, we propose to add mycoparasitism as one of the mechanisms that facilitated horizontal gene transfer between fungi. Mycoparasitic *Trichoderma* species have been suggested to have acquired by horizontal transfer nearly half its genes encoding plant cell wall-degrading carbohydrate-active enzymes and auxiliary proteins enabling them to be parasitism on a broad range of *Ascomycota* (Druzhinina et al., 2018). *H. pulvinata* 414-3 did not establish a direct parasitic relationship with *A. oryzae* RIB40 as it did with *C. fulvum*, and no distinct anastomosis was observed, but it firmly adhered to and coiled around RIB40 hyphae, indicating basic compatibility facilitating horizontal between the two fungi. Since *H. pulvinata* completely kills the host *C. fulvum*, *C. fulvum* might not be a stable source of foreign DNA, but rather *H. pulvinata* might have acquired the gene cluster in direct contact with nonhost fungi. In the genomic sequence of *H. pulvinata*, a gene cluster derived from *Aspergillus* was found in this study, but no horizontal gene transfer between host fungi, including *C. fulvum*, has been found so far. Although transduction by mycoviruses or propagation of retrotransposons could also have facilitated horizontal gene transfer (Richards et al., 2006), no transposon-like sequences were detected around the *DPH* cluster.

Several environmental cues such as temperature, light, nutrients, pH, and competing or synergistic organisms influence fungal activity, including transcriptional regulation of secondary metabolism–associated gene clusters (Keller, 2019). Aflatoxin production in *A. flavus* is generally upregulated at a suboptimal 30 °C (Ching’anda et al., 2021). Similarly, we found that deoxyphomenone production was also highest at temperatures unsuitable for growth of *H. pulvinata* and *Aspergillus* species (**Supplementary figure S8**). In *A. flavus*, Δ*AfDPH1* mutants that no longer produced deoxyphomenone formed significantly less spores. These results suggest that in at least *A. flavus*, deoxyphomenone stimulates sporulation in poor nutritional environments. In addition, *A. flavus* strains tended to produce more deoxyphomenone than *A. oryzae* strains, but the level was not sufficiently high for antifungal activity. Gibbons et al. (2012) reported that the expression levels of many secondary metabolite gene clusters of *A. oryzae*, including those for *DPH* and aflatoxin, were repressed compared with those of *A. flavus*. *A. oryzae*, used to ferment rice to produce Japanese sake, is probably derived from *A. flavus*, a pathogen toxic to plants and animals (Gibbons et al., 2012). *A. oryzae* is cultured with the budding yeast *Saccharomyces cerevisiae* during sake brewing and might have lost the ability to produce aflatoxin during its domestication from *A. flavus* (Watarai et al., 2019) because aflatoxin is genotoxic to yeast (Keller-Seitz et al., 2004). Since *A. oryzae* is usually grown under optimal fermentation conditions, the gene encoding deoxyphomenone required for sporulation by *A. flavus* in unfavorable conditions may also have been silenced or lost in *A. oryzae*. In fact, external application of deoxyphomenone inhibited sporulation and colony formation only in *A. oryzae*, indicating that *A. oryzae*, which produces only low levels of the compound, is less tolerant to this sesquiterpene (**Supplementary figure S13**). The mycoparasite *H. pulvinata* also produced deoxyphomenone at temperatures unfavorable for its growth, but in the presence of its host fungus *C. fulvum*, expression of the *HpDPH* genes was strongly induced, suggesting that deoxyphomenone might be crucial for mycoparasitic activity of *H. pulvinata*. However, the *ΔHpDPH1* mutant strains of *H. pulvinata* parasitized *C. fulvum*, comparable to the wild-type strain, supporting that deoxyphomenone is not required directly for its mycoparasitism (Iida et al., 2018). Spore germination of *C. fulvum* treated with deoxyhomenone for 24 h was halved, also suggesting an epidemiological effect on the mycoparasitism of *H. pulvinata*.

Deoxyphomenone did not affect the organelles in *C. fulvum* spores, and the specific target is still unknown. Although it is not directly involved in the mycoparasitism of *H. pulvinata* and is not toxic to plants, it had significant fungistatic activity against plant pathogens—the host *C. fulvum* and the nonhost *P. fuligena* of *H. pulvinata*—which might indicate a potential effect on biotrophic fungal pathogens of tomato plants. *H. pulvinata* remains an efficient biocontrol agent that can be used as a biofungicide to suppress races and fungicide-tolerant strains of *C. fulvum* that cause problems in tomato crops world-wide (Iida et al., 2015; Mesarich et al., 2023). Among *Aspergillus* species commonly used in fermentation, the use of deoxyphomenone to control sporulation should promote high efficiency for industrial fermentation and enzyme production.

## MATERIALS AND METHODS

### Growth conditions for fungal strains and plants

Strains of *H. pulvinata* 414-3 and *C. fulvum* CF301 were grown on PDA (half-strength, BD Difco, Franklin Lakes, NJ, USA), MM agar or broth (15 g sucrose, 5 g ammonium tartrate, 1 g NH_4_NO_3_, 1 g KH_2_PO_4_, 0.5 g MgSO_4_×7H_2_O, 0.1 g NaCl, 0.1 g CaCl_2_×H_2_O, 25 μL 0.2 mg/mL biotin, 15 g agar and 1 mL trace elements per liter) at 25 °C in the dark for 1 and 2 weeks, respectively, then spores were collected in sterile distilled water. *A. oryzae* and *A. flavus* strains (**Supplementary figure S7**) from NITE-NBRC (www.nite.go.jp/nbrc/catalogue/) were cultured on PDA, MM agar and broth, and Czapek-Dox agar and broth (3 g NaNO_3_, 2 g KCl, 1 g KH_2_PO_4_, 0.5 g MgSO_4_×7H_2_O 20 g glucose and 15 g agar per liter adjusted to pH 6.5) at 30 °C in the dark for a week, and spores were collected in sterile distilled water containing 1% (v/v) Tween 20. The concentration of spores was determined using a hemocytometer. Strains were maintained on PDA at 25 °C for a few months. Spore suspensions containing 10% v/v glycerol of all fungi except *H. pulvinata* were preserved at –80 °C for long-term storage; *H. pulvinata* was stored in 10% v/v glycerol with 5% w/v trehalose.

For examining mycoparasitic interactions on tomato leaves, tomato cultivar Moneymaker, which lacks any apparent *Cf* resistance gene, was grown in plastic pots for 3 weeks in a climate chamber at 25 °C with 16 h light/8 h dark. Seedlings were transplanted to soil in the greenhouse and grown for 3 weeks. The lower sides of tomato leaves were sprayed with a spore suspension of *C. fulvum* CF301 (1 × 10^5^ spores/mL). After 2 weeks, a spore suspension of *H. pulvinata* 414-3 (1 × 10^5^ spores/mL) was sprayed on the brown lesions that had formed on the abaxial surfaces of the leaves. White mycelial patches of *H. pulvinata* were observed a week after inoculation.

### Electron microscopy

*C. fulvum* CF301 or *A. oryzae* RIB40 were cultured on a nylon membrane (0.45 μm; GVS Japan, Tokyo, Japan) placed on PDA in a petri dish and incubated as described above. The membranes were then transferred to water agar and incubated for a week in the same conditions. A spore suspension of *H. pulvinata* (1 × 10^5^ spores/mL) was sprayed on the test colony on the membrane, then white colonies were observed 1 week later.

Small pieces (1 × 1 cm^2^) were cut from the membrane and fixed in 2.5% (v/v) glutaraldehyde in 100 mM cacodylate buffer (pH 7.4) overnight at 4 °C, then washed three times in the buffer, fixed in 1% (w/v) osmium tetroxide in the buffer for 1 h at room temperature and washed three times in distilled water. Specimens were then dehydrated through a graded ethanol series, critical-point dried in absolute ethanol using Leica EM CPD300 critical point dryer (Leica Microsystems, Wetzlar, Germany), then mounted on stubs, coated with platinum-palladium, and observed with a Hitachi SU-8220 scanning electron microscope (Tokyo, Japan).

For observing the ultrastructure of *C. fulvum* spores with transmission electron microscopy (TEM), spores of strain CF301 were treated with deoxyphomenone (120 µM) at 25 °C for 24 h in the dark. Spores were collected by centrifugation, washed three times with sterile water, then fixed in phosphate-buffered 2% glutaraldehyde at 4 °C overnight. The samples were again collected by centrifugation, then treated with 2% potassium permanganate in phosphate buffer for 1 h at room temperature and post-fixed in 2% w/v osmium tetroxide in phosphate buffer for 2 h at 4 °C. The samples were dehydrated in a graded ethanol series and embedded in epoxy resin (Fujifilm Wako, Osaka, Japan) at 60 °C for 48 hours. Ultrathin sections were cut using an ultramicrotome Leica UCT Ultracut (Leica Microsystems), then stained with uranyl acetate for 15 min and lead solution for 5 min and observed with a Hitachi TEM H-7600.

### Assays of antifungal activity and plant toxicity of deoxyphomenone

*C. fulvum* spores were treated with deoxyphomenone [(+)-13-deoxyphomenone, sporogen-AO1; Apollo Scientific, Bredbury, UK] in 1% methanol (v/v) at 25 °C in the dark for 24 h. Then samples were observed for germination and hyphal elongation using a Nikon E600 light microscope (Tokyo, Japan).

The antifungal activity of deoxyphomenone was compared with that of an *N*-halo-alkylthioimide fungicide, captan (orthocide80; Arysta Life Science, Tokyo, Japan). *C. fulvum* spores (1 × 10^3^ cells/mL) were treated with distilled water, deoxyphomenone (120 µM), or captan (100 µM) on a hydrophobic TF0808 glass slide (Matsunami Glass, Osaka, Japan) for 24 h at 100% humidity, then each solution was replaced with fresh solution, and samples incubated another 24 h. Germinated spores were then counted using the Nikon light microscope. For the controls, dH_2_O or 1% methanol (v/v) were used. Each treatment was done in triplicate, and at least 200 spores and hypha were assessed for each treatment.

Spore suspensions of the strains *C. fulvum* CF301, *P. fuligena* Pf17923, kindly provided by K. Nakajima and T. Kawakami, and *F. oxysporum* f. sp. *lycopersici* CK3-1 were adjusted to 1 × 10^6^ spores/mL and 10 μL were dropped on MM agar supplemented with different concentrations of deoxyphomenone (20 to 120 μM). The strains were cultured at 25 °C in the dark for 2 weeks or 3 days.

To evaluate the toxicity of deoxyphomenone on tomato plants, we injected the abaxial surface of three leaves on each of two 1-month-old tomato plants with about 20 µL of deoxyphomenone (20 to 80 µM) or with 1% methanol (v/v) as a control. The plants were then incubated in a climate chamber for 1 week, then leaves were examined for necrosis.

### Phylogenetic analyses of deoxyphomenone biosynthetic gene clusters

Deoxyphomenone biosynthetic genes and *DPH* clusters were initially identified in the genomic sequences of *H. pulvinata* 414-3 (Sushida et al., 2019) and *A. oryzae* RIB40 (Machida et al., 2005) using the BLAST program (blast.ncbi.nlm.nih.gov). Sequences shown in **Supplementary table S1 and S2** were aligned using MAFFT online version 7 (Katoh and Standley, 2013) with default parameters and trimmed using TrimAl version 1.4 (Capella-Gutiérrez et al., 2009). Maximum likelihood phylogenetic trees were constructed using RAxML version 8.2.12 with 1000 bootstraps; the PROTCATAUTO and GTRCAT models were selected for amino acid and nucleotide sequences, respectively (Stamatakis, 2014). Dot plot alignments were generated from the MAFFT analysis (**Supplementary figure S4**). Pairwise sequence alignments of *DPH6* homologous genes were performed using EMBOSS Needle (Madeira et al. 2022).

The phylogenetic species tree in **Figure 8** was constructed using maximum likelihood based on the 500 monocore genes (a single homolog in each of the species) using CVtree version 3.0 (*k* = 16) (Qi et al., 2004) by uploading the genomic information for the 60 published fungi shown in **Supplementary table S1**.

Reconciliation analysis between species and gene trees was performed using NOTUNG v.2.9 (Stolzer et al., 2012) according to the instructions (version 2.8 beta) to infer the evolutionary trajectories of the *DPH* clusters. Event scores were calculated as the total costs of duplications, transfers and losses. Costs/weights were set as duplications (D), 1.5; transfers (T), 8.0; losses (L), 1.0 (ratio D:T:L is 1:5.3:0.67) for amino acid sequences, and D, 1.5; T, 6.0; L, 1.0 (ratio D:T:L is 1:4:0.67) for nucleotide sequences for the gene clusters.

### Quantitative detection of deoxyphomenone

*H. pulvinata* (1 × 10^6^ spores) was cultured in 50 mL of MM broth with shaking at 25 °C in a light/dark cycle of 16 h/8 h for 2 weeks. *A. oryzae* and *A. flavus* (1 × 10^6^ spores) were incubated in MM broth without shaking at 25 °C in the dark for 2 weeks. The culture supernatants were collected by centrifugation and filtered through a 0.45 μm filter (Merk, Darmstadt, Germany). Deoxyphomenone was quantified using a 4000 QTRAP LC-MS/MS system (Sciex, MA, US) equipped with a 1290 series HPLC system (Agilent Technologies, CA, US) as previously reported (Iida et al., 2018). Commercial deoxyphomenone (Apollo Scientific) was used as a standard.

### Construction of transformation vectors

Genomic DNA was extracted from *H. pulvinata* and *A. oryzae* mycelia grown in MM broth for a week using Nucleo-Mag Plant (TakaraBio, Shiga, Japan) according to the manufacturer’s instructions. The DNA was then amplified by PCR using PrimeSTAR GXL Premix (TakaraBio) or KOD ONE (TOYOBO, Osaka, Japan) according to the instructions.

For generating the gene knockout vector, the upstream (1.2 kb) and downstream (1.0 kb) regions of *HpDPH1* were amplified from *H. pulvinata* genomic DNA using primer sets Dpprx2LF1/Dpprx2LR1 and Dpprx2UF1/Dpprx2UR1, respectively (**Supplementary table S3**). Geneticin-resistance gene (*gen*) cassettes were amplified from pRM254 (Mehrabi et al., 2015) using primers pRM254 attL1_F/pRM254 attL1_R. These primers contain a 5′ overhang sequence to overlap the sequences at the ends of the linearized plasmid or the *gen* cassette (**Supplementary table S3**). Plasmid pPM43GW was amplified using the primers pPM43GW_RB_F and pPM43GW_RB_R to linearize and remove the *ccdB* gene, which encodes a product that is lethal to bacterial cells. Four fragments were combined to generate the gene replacement vector pGW43_dpprx2ko using the In-Fusion EcoDry Cloning Kit (TakaraBio).

To introduce the *AoDPH1* gene into Δ*HpDPH1* strain KO10, *AoDPH1* (without introns) and hygromycin resistance gene (*hph*) cassettes were synthesized by VectorBuilder, Inc. (Chicago, IL, USA). The *AoDPH1* and *hph* genes were driven under the promoters of the *A. nidulans trpC* gene and *Cochliobolus heterostrophus gpd1* (NCBI accession X02390.1 and X63516.1), respectively (**Supplementary figure S9A**). Synthesized DNA was inserted into the blunt end of the PvuII-digested plasmid pPZP-PvuII kindly provided by Prof. Chihiro Tanaka.

To construct gene replacement vector for *AoDPH1* and *AfDPH1* gene, a plasmid pSH75 (Kimura and Tsuge 1993) was digested with *Bgl* II and *Hin*d III, and the resulting 2.4 kb fragment was used as a backbone. Marker gene cassettes (4.7 kb) containing a pyrithiamine resistant gene (*ptrA*) and an *EGFP* gene cassette were amplified from a plasmid pPTREXeGFP using the primers pPTREXeGFP_F and pPTREXeGFP_R. The upstream (1.5 kb) and downstream (1.5 kb) regions of *AoDPH1* were amplified from genomic DNA of *A. oryzae* RIB40 using primer sets AoDPH1_LB_F/AoDPH1_LB_R and AoDPH1_RB_F/AoDPH1_RB_R, respectively (**Supplemental table S3**). These primers contain a 5′ overhang sequence to overlap the sequences at the ends of the pSH75 fragment or the marker cassette (**Supplementary table S3**). These four fragments were combined using the In-Fusion EcoDry Cloning Kit (TakaraBio), then the resulting plasmid was named pSH75PTRdAoDPH1. The plasmid was linearized by digestion with *Swa* I and used for fungal transformation.

Plasmid vectors were introduced into *Escherichia coli* DH5α (NIPPON GENE, Tokyo, Japan) and extracted using MagExtractor-Plasmid-(TOYOBO) according to the instructions. The correct orientation of the fragments in the final constructs was confirmed by PCR.

### Fungal transformation

All transformations were done as previously described (Okmen et al., 2013) with some modifications for *H. pulvinata*. Briefly, *A. tumefaciens* (*Rhizobium radiobacter*) strain AGL-1 was transformed with the plasmid vectors used for the *H. pulvinata* transformation using the Gene Pulser electroporator (Bio-Rad Laboratories, Hercules, CA, USA) according to the instructions, then grown on LBman agar (10 g tryptone, 10 g NaCl, 5 g yeast extract, 10 g mannitol, 20 g agar per liter) supplemented with appropriate antibiotic at 28 °C for 3 days. Colonies were then collected and resuspended in IM broth (1 g glucose, 2.05 g K_2_HPO_4_, 1.45 g KH_2_PO_4_, 0.15 g NaCl, 0.50 g MgSO_4_·7H_2_O, 0.07 g CaCl_2_·2H_2_O, 0.5 g (NH_4_)_2_SO_4_, 0.5% (w/v) glycerol, 8.53 g 2-(*N*-morpholino)ethanesulfonic acid [pH 5.3] per liter) supplemented with 200 µM acetosyringone at an optical density at 600 nm (OD_600_) of 0.2. The bacterial suspension (200 µL) was mixed with 200 µL of a spore suspension of *H. pulvinata* (1.0 × 10^5^ cells/mL), then the suspension was spread on a nylon membrane (GVS Japan) that had been placed on IM agar (IM; 20 g per liter) in a plate. After 2 days of incubation in the dark at 25 °C, the membranes were transferred to PDA supplemented with either 200 µg/mL geneticin (G418 sulfate; Fujifilm Wako) or 100 µg/mL hygromycin (Fujifilm Wako), and 50 µg/mL meropenem trihydrate (Fujifilm Wako). Transformed fungal colonies appeared after a week were transferred to new PDA plates supplemented with geneticin (Fujifilm Wako) or hygromycin (Fujifilm Wako) and meropenem trihydrate (Fujifilm Wako) at the same concentration as described above and incubated in the dark at 25 °C for a week. Gene replacement was confirmed by PCR amplification using the primers listed in **Supplementary table S3**.

Protoplasts of *A. oryzae* RIB40 and *A. flavus* NBRC114564 were transformed using polyethylene glycol. First, 2.4 × 10^8^ spores harvested from a 10-d-old culture on Czapek-Dox agar were resuspended in 200 mL of Czapek-Dox broth and incubated at 30 °C at 160 rpm for 18 h. The mycelium was then collected using miracloth (Merck, NJ, USA) and washed once with distilled water, then incubated in 20 mL of a protoplast solution consisted of 10 mM phosphate buffer (pH6.0), 0.8 M NaCl, 10 mg/mL Lysing enzyme (Sigma-Aldrich, MO, USA), 5 mg/mL Cellulase Onozuka R-10 (Yakult Pharmaceutical, Tokyo, Japan), and 2.5 mg/mL Yatalase (Takara, Shiga, Japan) at 30 °C with shaking at 83 rpm for 3 h. The protoplasts were filtered through a mesh and centrifuged at 2000 × *g* and 4 °C for 5 min. The pellet was washed once in 0.8 M NaCl and centrifuged at 2000 × *g* and 20 °C for 5 min, then resuspended in 1.2 mL of solution 1 (Sol1: 9.35 g NaCl, 2 mL 1 M CaCl_2_, 2 mL 1 M Tris-HCl per 200 mL). Two handled fourty microliters of solution 2 (Sol2: 40% (w/v) PEG4000, 10 mL 1 M CaCl_2_, 10 mL 1 M Tris-HCl per 200 mL) was added to the protoplast suspension and the resulting mixture was dispensed 300 μL each into 14-mL round tubes A linearized vector DNA was added to the mixture and held on ice for 30 min, then 1 mL of Sol2 was added and held at 25 °C for 20 min. The protoplasts were washed once in 10 mL of Sol1, and centrifuged at 2000 × *g* at 20 °C for 5 min. After the supernatant was discarded, the protoplasts were resuspended in 300 μL of Sol1, then placed on a regeneration selection medium consisted of Czapek-Dox agar supplemented with 5% (w/v) NaCl, 0.1 μg/mL pyrithiamine hydrobromide (Sigma-Aldrich, MO, USA). The resulting plates were overlayed with 5 mL of soft agar medium consisted of 5% (w/v) NaCl and 0.5% (w/v) agar, and incubated at 30 °C for 4 d. The transformants grown on the regeneration selection medium were isolated and transferred to Czapek-Dox agar containing 0.2 μg/mL pyrithiamine hydrobromide and incubated at 30 °C for another selection. Gene replacement of the transformants was confirmed by PCR amplification using the primers listed in **Supplementary table S3**.

### Quantitative real-time PCR

*C. fulvum* CF301 and *H. pulvinata* 414-3 were cultured on nylon membranes (GVS Japan) on PDA at 25 °C in the dark for 1 and 2 weeks. The membranes were transferred to water agar and then incubated in the same conditions for a week. *C. fulvum* colonies were sprayed with a spore suspension of *H. pulvinata* (1 × 10^5^ spores/mL) and checked for white colony growth of *H. pulvinata* 1 week later. Total RNA was extracted from the mycelia using the RNeasy Plant Mini Kit (Qiagen, Hilden, Germany). cDNA libraries were generated using SuperScript IV VILO Master Mix with ezDNase Enzyme (ThermoFisher Scientific, Waltham, MA, USA) according to the manufacturer’s instructions. Primers for quantitative real-time PCR of 90– 150-bp fragments of cDNA were designed (**Supplementary table S3**), and the PCR was run on the LightCycler 480 System (Roche, Basel, Switzerland) using the KAPA SYBR Fast qPCR Kit (Nippon Genetics, Tokyo, Japan) according to the manufacturer’s instructions. Relative expression levels were calculated using the comparative CT (2^−ΔΔCT^) method (Livak and Schmittgen, 2001). The data were normalized to the transcript level of the *H. pulvinata* actin gene (**Supplementary table S3**). Transcript levels of target genes in each RNA sample were measured for three independent experiments, each with two replicates.

### In-well assay and observation of mycoparasitism

Green fluorescence protein (GFP)-expressing *C. fulvum* strain CF301gfp (Iida et al., 2018) and *H. pulvinata* strains were cultured on PDA as mentioned above to prepare spore suspensions of CF301gfp (5 × 10^4^ spores/mL) and *H. pulvinata* (1 × 10^6^ spores/mL). Then 50 µL of a spore suspension of CF301gfp and of each *H. pulvinata* strain were added to 100 µL of MM broth without a carbon source (MM without sucrose) in each well of a 48-well plate. At least 8 wells were used per treatment, and CF301gfp was used as the control. The plates were incubated at 25 °C in the dark for 3 d. Hyphae were observed using a fluorescence microscope BZ-X800 (KEYENCE, Osaka, Japan); lack of GFP fluorescence from *C. fulvum* was correlated with the mycoparasitic activity of *H. pulvinata* using the hybrid cell count system of the microscope. We named this simple method for detecting mycoparasitism the IWAO method, derived from “in-well assay and observation” and in honor of the method’s developer, E. Iwao.

### Statistical analyses

Means and standard deviations of number of spores, spore germination, and hyphal length were calculated. Significant differences (*P* < 0.05) in the number of spores produced by the different fungal strains, percentage spore germination and hyphal lengths of *C. fulvum*, and colony diameter of *Aspergillus* strains were evaluated using either Tukey’s test or Williams’ multiple comparison. Significant differences (*P* < 0.05) in gene expression levels were determined using Welch’s *t*-test followed by Bonferroni–Holm correction for multiple testing. All data were analyzed in the program R version 4.0.3 (www.r-project.org).

## Data availability

The genomic sequence of *H. pulvinata* 414-3 is available in the DDBJ repository under an accession number BJKQ00000000 (BioProject: PRJDB8178) (Sushida et al., 2019). Other fungal sequences shown in **Supplementary table S1** are available on the fungal genome portal MycoCosm (mycocosm.jgi.doe.gov) of the Joint Genome Institute (Lawrence Berkeley National Institute, Berkeley, CA, USA; Grigoriev, 2014) and in the DDBJ/EMBL/GenBank repository (www.ncbi.nlm.nih.gov). Auxiliary data are available through Figshare (doi.org/10.6084/m9.figshare.24559120.v1).

## ACKNOWLEDGEMENTS

We are grateful to Kentaro Ikeda and Hiroshi Sakai for providing *H. pulvinata* strains, Daisuke Hagiwara for *A. oryzae* RIB40 and plasmid vector pPTREXeGFP, and Kaori Nakajima and Taku Kawakami for the *P. fuligena* strain, Yasuyuki Kubo and Sayo Kodama for valuable suggestions, Chieko Tanaka for laboratory management, and Pierre J.G.M. de Wit for critical reading of the manuscript. This work was supported by a Grant-in-Aid for Scientific Research from JSPS (17H05022 and 20H02993) (Y. Iida).

## Conflict of interest

The authors declare that they have no commercial or financial relationships that could be construed as a potential conflict of interest.

## Author contributions

YI designed the study and managed the research funds. KM, HS and YH performed qRT-PCR and analyzed the data. KM, TaS and ON transformed fungal strains. ToS did the electron microscopy and EI did the light microscopy. KM, TaS, YH and HN did the mass spectrometry analyses. MZF and YN predicted the biosynthetic pathways and identified the genes for enzymes. KM and HS did the phylogenetic analyses. KM and YI wrote the paper. All authors contributed to the article and approved the submitted version.

## Supplementary figures

Figure S1. Effect of deoxyphomenone on *Cladosporium fulvum* and tomato leaves.

Figure S2. Phylogenetic tree of fungal sesquiterpene cyclases.

Figure S3. Alignment and homology of the amino acid sequences of the predicted sesquiterpene cyclases identified in *Hansfordia pulvinata* and *Aspergillus oryzae* genome sequences.

Figure S4. Dot plot analysis of the deoxyphomenone biosynthesis (*DPH*) gene clusters from *Hansfordia pulvinata* 414-3 and *Aspergillus oryzae* RIB40.

Figure S5. Exon-intron structures of deoxyphomenone biosynthetic (*DPH*) genes.

Figure S6. Alignment of deduced amino acid sequences of *HpDPH6* homologs from *Hansfordia pulvinata* and six *Aspergillus* species.

Figure S7. Phylogenetic analysis of genomic sequences containing *HpDPH1* homologous genes and the upstream region using the maximum likelihood method.

Figure S8. Deoxyphomenone produced by *Hansfordia pulvinata* 414-3 and strains of *Aspergillus oryzae* and *A. flavus*.

Figure S9. Deoxyphomenone production by *Hansfordia pulvinata* 414-3, *Aspergillus oryzae* RIB40 and *A. flavus* NBRC114564 on MM agar or in broth at different temperatures.

Figure S10. PCR detection of the inserted cassettes in transformants and wild-type strains.

Figure S11. *In vitro* assay of *Hansfordia pulvinata* mycoparasitic activity against GFP-expressing *Cladosporium fulvum*.

Figure S12. Deoxyphomenone production and sporulation of *A. oryzae* RIB40 and transformants.

Figure S13. Effect of deoxyphomenone on mycelial growth of *Hansfordia pulvinata, Aspergillus oryzae* and *A. flavus*.

Figure S14. Reconciliation of gene trees with species tree based on *DPH* nucleotide and amino acid sequences.

Figure S15. Hyphal contact between the mycoparasite *Hansfordia pulvinata* 414-3 and *Aspergillus oryzae* RIB40.

Supplemental table S1. Fungal genome sequences used in this study.

Supplemental table S2. Amino acid sequences of *DPH* biosynthesis genes from *Hansfordia pulvinata* and *Aspergillus* species.

Supplemental table S3. Primer sequences used in this study.

## Notes

https://doi.org/10.6084/m9.figshare.24559120.v1

### Competing Interest Statement

The authors have declared no competing interest.

### Summary of Updates

Added ORCiD numbers for some authors.

