## Supplemental figures and tables for "Adaptive evolution of sesquiterpene deoxyphomenone in mycoparasitism by *Hansfordia pulvinata* associated with horizontal gene transfer from *Aspergillus* species"

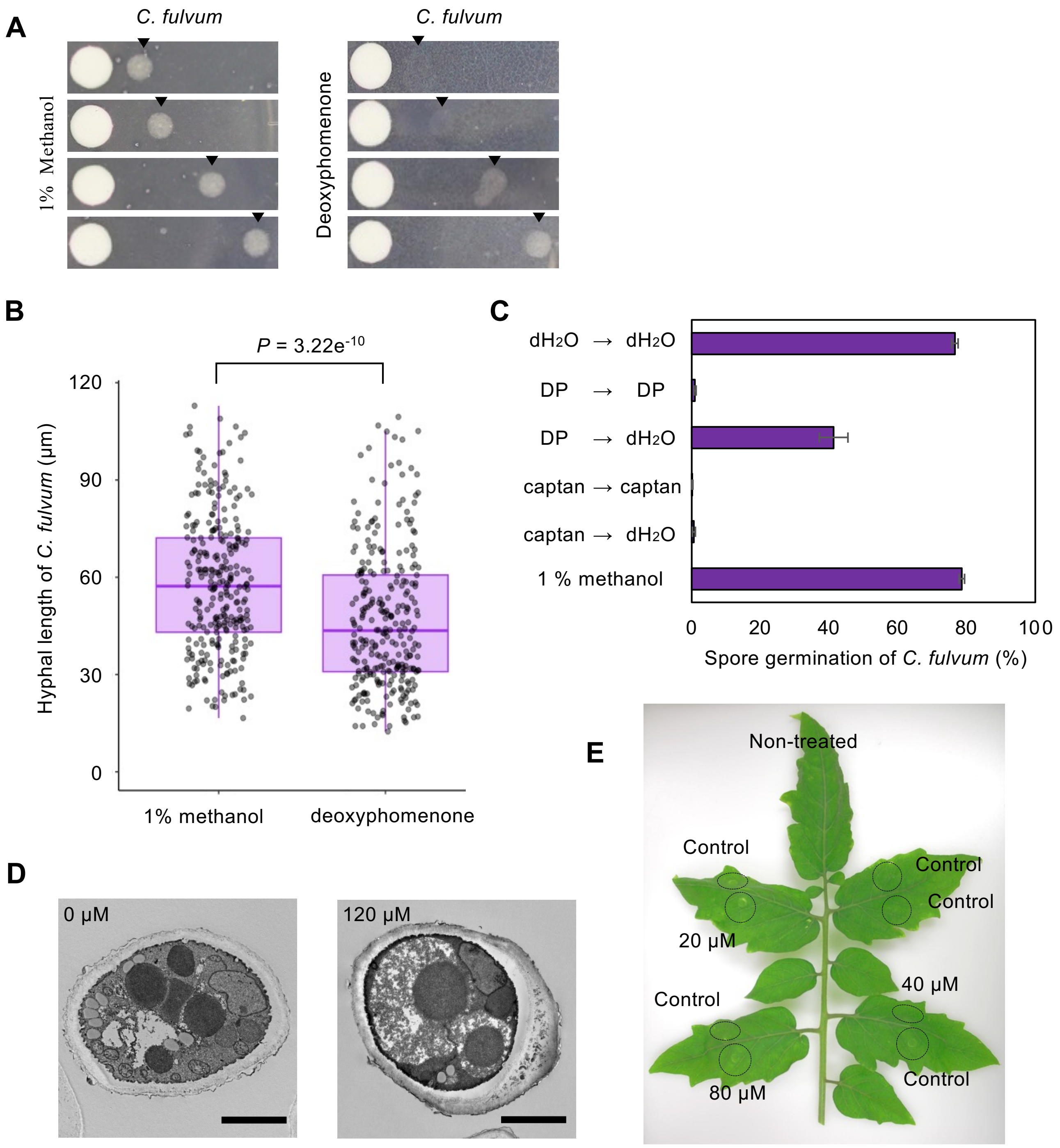


**Figure S1. Effect of deoxyphomenone on *Cladosporium fulvum* and tomato leaves.**
Deoxyphomenone (120 µM) and 1% methanol (v/v) as a control were used. (**A**) Effect on spore germination. A filter paper containing deoxyphomenone was placed on the left on PDA, then a spore suspension of *C. fulvum* was dropped (arrowheads) at different distances from the paper and incubated at 25 ºC for 2 weeks. (**B**) Effect on hyphal elongation. Spores were germinated in a sterile distilled water at 25 ºC for 24 h. Germinated spores were treated with deoxyphomenone, then hyphal length was measured after 24 h using a light microscope. The line in the center of the box indicates the median; box margins represent the 25th and 75th percentiles. The length of the box is the interquartile range; whiskers indicate the minimum and maximum. Significant differences among treatments were determined using Tukey's test. (**C**) Fungistatic activity of deoxyphomenone. Spores of *C. fulvum* were suspended in distilled water (dH2O), deoxyphomenone (DP) or 100 µM fungicide captan for 24 h, then solutions were replaced with fresh solutions as shown, and treated for a further 24 h. Germinated spores were counted using a light microscope. Continuous treatment with dH2O or 1% methanol (v/v) was used as a positive control. Values are the means of three replicates (± SD). (**D**) Transmission electron micrographs of interior of *C. fulvum* spores 24 h after treatment with deoxyphomenone. Bars = 2 µm. (**E**) Evaluation of 20, 40 and 80 µM deoxyphomenone or 1% methanol (v/v) as a control for toxicity on tomato leaves 2 weeks after 1-month-old leaves were injected with the compound.

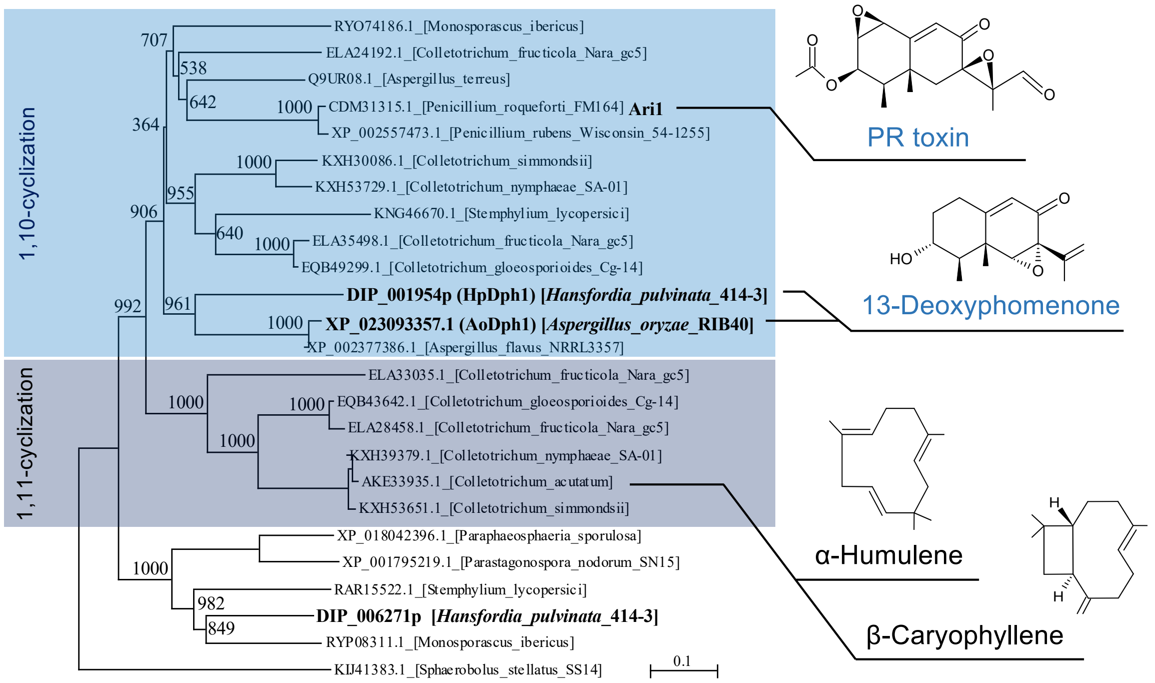


**Figure S2. Phylogenetic tree of fungal sesquiterpene cyclases.**
The phylogenetic tree was generated from amino acid sequences encoding sesquiterpene synthases by maximum likelihood phylogenetic analysis. Predicted aristolochene synthase-like proteins identified in *H. pulvinata* 414-3 and *A. oryzae* RIB40 genome sequences, and the aristolochene synthase Ari1 are shown in bold. GenBank/EMBL/DDBJ accession numbers or unique numbers for *H. pulvinata*are are indicated. *Sphaerobolus stellatus* (*Basidiomycota*) was used as an outgroup. Numbers beside the branches represent bootstrap values based on 1000 replicates. The scale bar corresponds to 0.1 estimated amino acid substitutions per site. Representative metabolites and proteins associated with their cyclization are connected by lines.

```
DIP-001954p(HpDPH1)     1 MLSTLRSFELHKLLGLTGSSSNDSTTTTTTITTTAASAAPQPKAQQQPQP    AYPSESEDMARDRRRFPSLAATTDTASPSPSAFAPEIHPAADRVSREVAE    100
                                                 .......::|:|.....||.:...|:.    ..      :.:|||..|.||    ..:|.||||:..|||.:|.||.||..
XP_023093357(AoDPH1)    1 -----------------------MLQRLWALSTSAIKLPFQPFSFGAPRD    LL------IEKDRRSMPCLA----KEAPPPSAFSATIHPLSDSVSTEVDN     67

DIP-001954p(HpDPH1)   101 AFLARWPFRGEAERRRFRAADFPRVTCLYFPHARAERIGFACRLLTLLFL    VDDLLEEMGLEEGSAYNERLISISKGDEEPEPDKPAEVLTRELWEDMRAC    200
                          .||..||||.:.||.||.||.|.||||||||.|..:|||||||:||:|||    :|||||||.|:|||.|||:|||||:||..|:...||:.:..:||||||||
XP_023093357(AoDPH1)   68 YFLQNWPFRTDNERARFHAAGFSRVTCLYFPMAMDDRIGFACRMLTILFL    IDDLLEEMSLDEGSTYNEKLISISRGDVAPDRTIPAQWIMYDLWEDMRAC    167

DIP-001954p(HpDPH1)   201 DARLAGEIKEPVFTFMRAQTDGSRLTLRELGAYFEYREKDVGQA------    ----LLSALMRFSMALHLTPAQLRLAQPLERNCARHISVVNDIYSWRKEV    290
                          |..||.|:.||||||||||||.:|||:.:.|.|.:|||||||||              |||.|.|::|.|:||...||:|.|.|||||:||:::|||||||||:
XP_023093357(AoDPH1)  168 DHVLADELLEPVFTFMRAQTDKTRLTIHQFGEYLDYREKDVGQAIKAQTK    CIRSLLSGLQRYTMKLYLTEEDLRMAAPAERNCAKHIAILNDIYSWRKEL    267

DIP-001954p(HpDPH1)   291 RASETLHEEGAALCSSVAVLAAEASVPASGARRVLWALCREWEAEHRRME    GDLLRRLVEEEEEEGDQGGQVLRVYVRGLESQMSGNERWSESTPRYHKVK*   391
                          .||:|||.||||:||||.||:...::..:..:||||.:|||||:.|:::               ..|....|.:..|..|:.|||.||||||||||||||||..
XP_023093357(AoDPH1)  268 LASKTLHHEGAAICSSVQVLSEVTALSHAATQRVLWTMCREWESVHKQL-    ----------VTEVAGTGSRDLLDYIHGLEFQMSGNERWSESTPRYHF*--   355
```

Identity: 197/401 (49.1%) / Similarity: 247/401 (61.6%) / Gaps: 56/401 (14.0%)

```
DIP-001954p(HpDPH1)     1 -----------MLSTLRSFELHKLLGLTGSSS-----NDSTTTTTTITTT    AASAAPQPKAQQQPQPAYPSESEDMARDRRRFPSLAATTDTASPSPSAFA     84
                                     .|||:.|  |.:|..:...||     |.|:..:.|..||    |.:..|.|:..|:|                           |...|:...
DIP-006271p             1 MASALLSLPTAALSTILS--LVRLSSVVSPSSTITTKNPSSNASATEETT    ATTPPPVPENSQRP---------------------------AGLRPTRLT     71

DIP-001954p(HpDPH1)    85 PEIHPAADRVSREVAEAFLARWPFRGEAERRRFRAADFPRVTCLYFPHAR    AERIGFACRLLTLLFLVDDLLEEMGLEEGSAYNERLISISKGDEEPEPDK    184
                          ...|...::...:|.:.||..|||:.:..||||....:....|:..|.:.    .|||.:.|||||:.||:|||::.|.:.||:|:|..::...:|.:.|:.|.
DIP-006271p            72 ARKHRLTEQTVEQVNDFFLRNWPFKTDKHRRRFVDEGYAFFVCVLVPESL    DERIHWGCRLLTVGFLIDDLVDNMNVAEGAAFNAAVVECCRGTQLPDRDV    171

DIP-001954p(HpDPH1)   185 PAEVLTRELWEDMRACDARLAGEIKEPVFTFMRAQTDGSRLTLRELGAYF    EYREKDVGQALLSALMRFSMALHLTPAQLRLAQPLERNCARHISVVNDIY    284
                          |::.:..:|:|.|||.|.|||.|:.:|...|:.||.||||.....|..||    |||:.|:|:.|:|.:|||...|.:|.|:|.:.:|::.|..:||:.|||:.
DIP-006271p           172 PSQWIMYDLFEAMRAVDRRLADELLQPTIDFLLAQVDGSRRRPMNLAEYF    EYRDADLGKGLISGIMRFCGGLSMTTAELDVVRPVDENVMKHITFVNDVC    271

DIP-001954p(HpDPH1)   285 SWRKEVRASETLHEEGAALCSSVAVLAAEASVPASGARRVLWALCREWEA    EHRRMEGDLLRRLVEEEEEEGDQG-GQVLRVYVRGLESQMSGNERWSEST    383
                          |:.||..|:|..:|.| .:||||.::||...|....|:||:|...|.||.    .|..|:.|:|         .|..| ...||.|:|.:|.|.||||.||..|
DIP-006271p           272 SYEKERLAAEAGYELG-EICSSVPIVAAWLGVGEDDAKRVMWQAARGWED    RHLAMKRDIL---------AGPLGASSALRTYLRWVEYQASGNELWSLLT    361

DIP-001954p(HpDPH1)   384 PRYHKVK*------------    391
                          |||::...
DIP-006271p           362 PRYNRFGALGFTEGRPEAQ*    381
```

Identity: 140/420 (33.3%) / Similarity: 207/420 (49.3%) / Gaps: 68/420 (16.2%)

```
XP_023093357(AoDPH1)    1 MLQRLWALSTSA-------IKLPFQPFSFGAPRDLLIEKDRRSMPCLAKE    ----APP-------------PSAFSATIHPLSDSVSTEVDNYFLQNWPFR     76
                          |...|.:|.|:|       ::|.    |..:|...:..|:..|.....:|        .||             |:..:|..|.|::....:|:::||:||||:
DIP-006271p             1 MASALLSLPTAALSTILSLVRLS----SVVSPSSTITTKNPSSNASATEE    TTATTPPPVPENSQRPAGLRPTRLTARKHRLTEQTVEQVNDFFLRNWPFK     96

XP_023093357(AoDPH1)   77 TDNERARFHAAGFSRVTCLYFPMAMDDRIGFACRMLTILFLIDDLLEEMS    LDEGSTYNEKLISISRGDVAPDRTIPAQWIMYDLWEDMRACDHVLADELL    176
                          ||..|.||...|::...|:..|.::|:||.:.||:||:.||||||::.|:    :.||:.:|..::...||...|||.:|:|||||||:|.|||.|..||||||
DIP-006271p            97 TDKHRRRFVDEGYAFFVCVLVPESLDERIHWGCRLLTVGFLIDDLVDNMN    VAEGAAFNAAVVECCRGTQLPDRDVPSQWIMYDLFEAMRAVDRRLADELL    196

XP_023093357(AoDPH1)  177 EPVFTFMRAQTDKTRLTIHQFGEYLDYREKDVGQAIKAQTKCIRSLLSGL    QRYTMKLYLTEEDLRMAAPAERNCAKHIAILNDIYSWRKELLASKTLHHE    276
                          :|...|:.||.|.:|.......||.:||:.|:|          :.|:||:    .|:...|.:|..:|.:..|.:.|..|||..:||:.|:.||.||::. .:|
DIP-006271p           197 QPTIDFLLAQVDGSRRRPMNLAEYFEYRDADLG----------KGLISGI    MRFCGGLSMTTAELDVVRPVDENVMKHITFVNDVCSYEKERLAAEA-GYE    285

XP_023093357(AoDPH1)  277 GAAICSSVQVLSEVTALSHAATQRVLWTMCREWESVHKQLVTEV-AGT--    GSRDLLDYIHGLEFQMSGNERWSESTPRYHF*--------------    355
                          ...|||||.:::....:.....:||:|...|.||..|..:..:: ||.    .|..|..|:..:|:|.||||.||..||||:..
DIP-006271p           286 LGEICSSVPIVAAWLGVGEDDAKRVMWQAARGWEDRHLAMKRDILAGPLG    ASSALRTYLRWVEYQASGNELWSLLTPRYNRFGALGFTEGRPEAQ*    381
```

Identity: 134/396 (33.8%) / Similarity: 205/396 (51.8%) / Gaps: 56/396 (14.1%)

**Figure S3. Alignment and homology of the amino acid sequences of the predicted sesquiterpene cyclases identified in *Hansfordia pulvinata* and *Aspergillus oryzae* genome sequences.**
Amino acid sequences of two candidates of *H. pulvinata* 414-3, DIP_001954 (HpDph1) and DIP_006271, and one of *A. oryzae* RIB40 XP_023093357 (AoDph1) were compared.

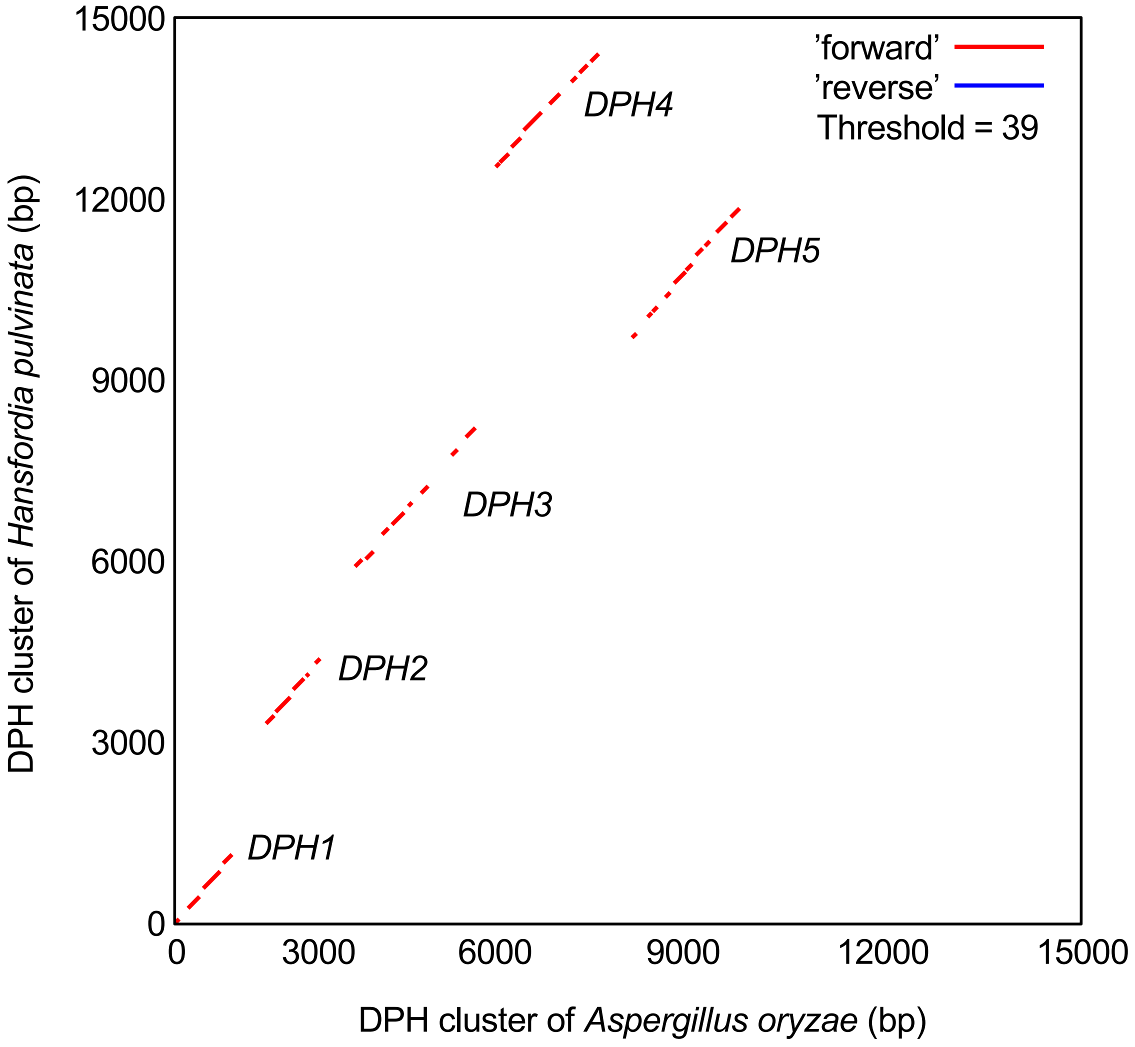


**Figure S4. Dot plot analysis of the deoxyphomenone biosynthesis (*DPH*) gene clusters from *Hansfordia pulvinata* 414-3 and *Aspergillus oryzae* RIB40.**

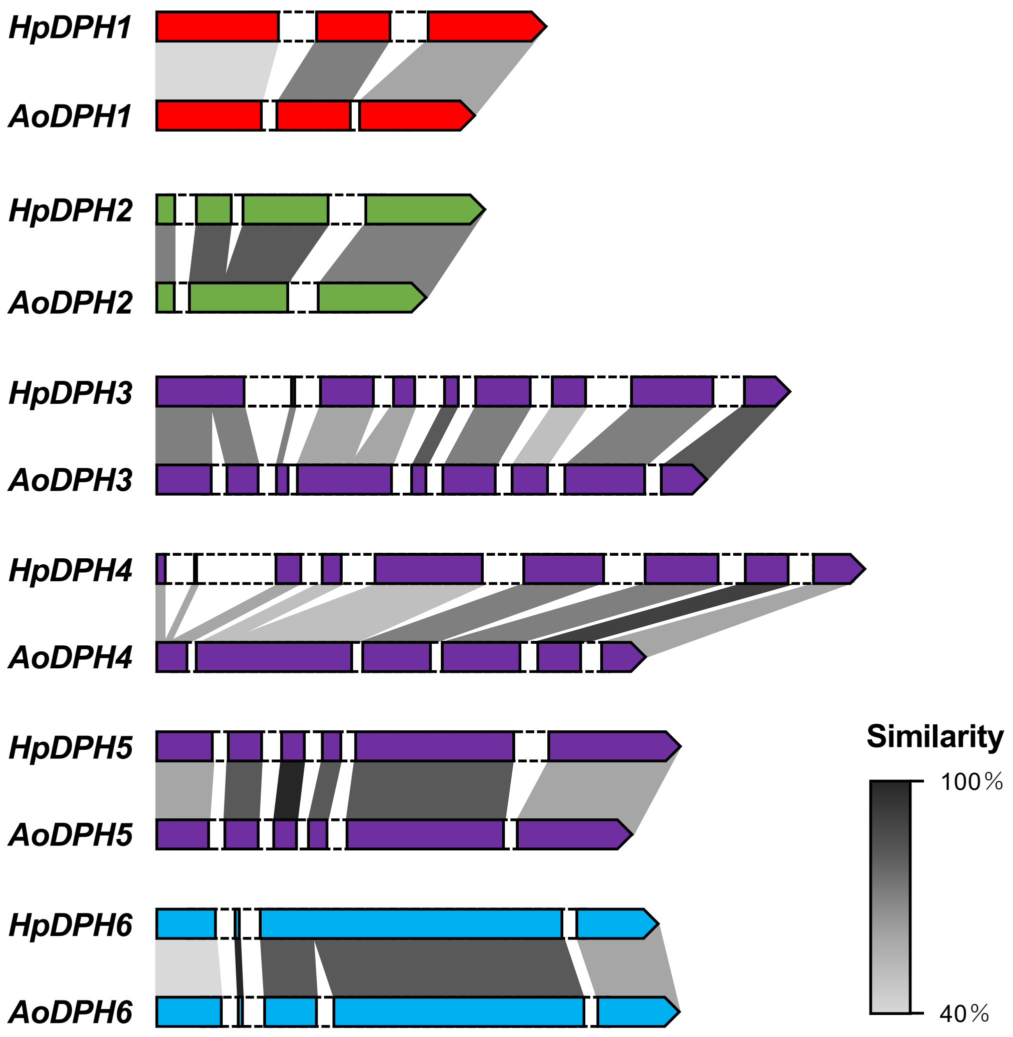


**Fig. S5. Exon-intron structures of deoxyphomenone biosynthetic (*DPH*) genes.**
Colored boxes: exon regions; boxes with dashed line: intron regions in *DPH* genes; gray shading: similarity of amino acid sequences.

```
H. pulvinata            MATEKHPA-----------SLPGSQDAPPKTEAPPQGVERDSDAYGSDGEHGGGSSITEN TDADQKGGGLVLVITVAALAMGMFLVSLDMTIVATAIPRITDEFGQIELVGWYGAAFFTT
A. oryzae               MSTKTRSSQTGETAVSSRISTPATLEKPNKDALPSH----DPQVMEVRNKEDSGSGVDDA PTEEPSTGGFTLLMTVTALAMSMFLVSLDMTIVATAIPKITDEFGGIEDEGWYGSAFFIT
A. parvisclerotigenus   MSTKTRSSQTGETAVSSRISTPATLEKPNKDALPSH----DPQVMEVRNKEDSGSGVDDA PTEEPSTGGFTLLMTVTALAMSMFLVSLDMTIVATAIPKITDEFGGIEDEGWYGSAFFIT
A. flavus               MSTKTRSSQTGETAVSSRISTPATLEKPNKDASPSH----DPQVMEVRNKEDSGSGVDDA PTEEPSTGGFTLLMTVTALAMSMFLVSLDMTIVATAIPKITDEFGGIEDEGWYGSAFFIT
A. minisclerotigenes    MSTKTRSSHTGETAVSSRISTPATPEKPNKDASPSH----DPQVMEVRNKEETGSGVDDA PTDEPSTGGFTLFMTVTALAMSMFLVSLDMTIVATAIPKITDEFGGIEDEGWYGSAFFIT
A. caelatus             MSTKTRSSQTGETAVSSGTSTPATLEKSNKDVSPSH----ETPVVGDREKEERGSGVDDA PTDKPSTGGFTFLMTVTALAMSMFLVSLDMTIVATAIPKITDEFGGIEDEAWYGSAFFIT
A. pseudocaelatus       MSTKTRSSQTGETAVSSGTSTHATLEKSNKDVSPSH----ETPVMGDREKEERGSGVDDA PTDKPSTGGFTFLMTVTALAMSMFLVSLDMTIVATAIPKITDEFGGIEDEAWYGSAFFIT
                        *:*:.:.:           *  .: : . *   *.:    :. .     :.  **.: :  .  . . **:.:.:**:****.****************:****** **   .***:*** *

H. pulvinata            LGGFQAAWGKAYKYFPLKITFLLSIFLFGVGSLICGVAPNSLTLIVGRAITGVGGAGVSC GCFTIIALSAPPKQQPAYIGILGASYGIAAAIGPLLGGAFTDKVTWRWCFYINLPIGAVT
A. oryzae               LGSFQAAWGKAYKYFPLKTSFLLSIFLFEVGSLICGVAPNSLALCIGRAITGVGGAGISS GAFTIIALSAPPKQRPAYIGILGASYGVAAAIGPLVGGAFTTNITWRWCFYINLPIGGLA
A. parvisclerotigenus   LGSFQAAWGKAYKYFPLKTSFLLSIFLFEVGSLICGVAPNSLALCIGRAITGVGGAGISS GAFTIIALSAPPKQRPAYIGILGASYGVAAAIGPLVGGAFTTNITWRWCFYINLPIGGLA
A. flavus               LGSFQAAWGKAYKYFPLKTSFLLSIFLFEVGSLICGVAPNSLALCIGRAITGVGGAGISS GAFTIIALSAPPKQRPAYIGILGASYGVAAAIGPLVGGAFTTNITWRWCFYINLPIGGLA
A. minisclerotigenes    LGSFQAAWGKAYKYFPLKTSFLLSIFLFEVGSLICGVAPNSLALCIGRAITGVGGAGISS GAFTIIALSAPPKQRPAYIGILGASYGVAAAIGPLVGGAFTTNITWRWCFYINLPIGGLA
A. caelatus             LGSFQAAWGKAYKYFPLKISFLLSILLFEIGSLICGVAPNSIALCIGRAITGVGGAGISS GAFTIIALSAPPKQRPAYIGILGASYGVAAAIGPLVGGAFTTNITWRWCFYINLPIGGLA
A. pseudocaelatus       LGSFQAAWGKAYKYFPLKISFLLSILLFEIGSLICGVAPNSIALCIGRAITGVGGAGISS GAFTIIALSAPPKQRPAYIGILGASYGVAAAIGPLVGGAFTTNITWRWCFYINLPIGGLA
                        **.************** :*****:** :***********::* :***********:*. *.************:************:*******:***** ::*************.::

H. pulvinata            VAVIAFFYKPPPPLPSLNEPLREKLLQLDIPGTLLLTAALVCFVLSLQWAGVSRPWSDST VIGTLVGFGVLLIAYAVVQWLQGPRAAMVGHLLQRNVVIAMVYIHLIAGSFFLLVYYLPI
A. oryzae               AGIILFFYRPPPPLPFLDTPLKEKLLQMDLVGTLILVSALVCYVLALQWGGISLSWGDSK VIGTLVGFVVLLAVYAIVQWVQKERAAMIGRLFRRNVVVMMIYIDLLAGTFFLLVYYLPI
A. parvisclerotigenus   AGIILFFYRPPPPLPFLDTPLKEKLLQMDLVGTLILVSALVCYVLALQWGGISLSWGDSK VIGTLVGFVVLLAVYAIVQWVQKERAAMIGRLFRRNVVVMMIYIDLLAGTFFLLVYYLPI
A. flavus               AGIILFFYRPPPPLPFLDTPLKEKLLQMDLVGTLILVSALVCYVLALQWGGISLSWGDSK VIGTLVGFVVLLAVYAIVQWVQKERAAMIGRLFRRNVVVMMIYIDLLAGTFFLLVYYLPI
A. minisclerotigenes    AGIILFFYRPPPPLPFLDTPLKEKLLQMDLVGTLILVSALVCYVLALQWGGISLSWGDSK VVGTLVGFVVLLAVYAIVQWVQKERAAMIGRLFRRNVLVMMIYIDLLAGTFFLLVYYLPI
A. caelatus             GGIILFFYRPPPPLPFLDTPLTEKLLQMDLPGTLILVSALVCYVLALQWGGVSLSWGDSK VVGTLVGFGVLLVVYAIVQRAQKERAAMIGRLFRRNVVIMMIYIDLLAGTFFLLVYYLPI
A. pseudocaelatus       GGIILFFYRPPPPLPFLDTPLTEKLLQMDLPGTLILVSALVCYVLALQWGGVSLSWGDSK IVGTLVGFGVLLVVYAIVQRAQKERAAMIGRLFRRNVVIMMIYIDLLAGTFFLLVYYLPI
                        .:* ***:****** *: ** *****:*: ***:*.:****:**:***.*:* .*.**. ::****** *** .**:**  *  ****:*:*::***:: *:**.*:**:*********

H. pulvinata            YFQVVNGVSAAQSGIRNLPLILGQSVFATISGVLLSKFERPQPFLIAGATLVTISSGLLY TLDVDTEPGKWIGYQLLAGVGVGWCFQVPVVTAQSTVQPEHVSSVTAMVLTVQTIGGSIF
A. oryzae               YFQVVSGVSAAQSGIRNLPLILAQSVSTVISGVTLSKFGYPQPFLLVGGVLTAIGSGLLY TLEVNTGSGKWIGYQLLAGIGIGWCFQIPVVTAQASVAPEDLPSVTAMVLTVQTLGGSVF
A. parvisclerotigenus   YFQVVSGVSAAQSGIRNLPLILAQSVSTVISGVTLSKFGYPQPFLLVGGVLTAIGSGLLY TLEVNTGSGKWIGYQLLAGIGIGWCFQIPVVTAQASVAPEDLPSVTAMVLTVQTLGGSVF
A. flavus               YFQVVSGVSAAQSGIRNLPLILAQSVSTVISGVTLSKFGYPQPFLLVGGVLTAIGSGLLY TLEVNTGSGKWIGYQLLAGIGIGWCFQVPVVTAQASVAPEDLPSVTAMVLTVQTLGGSVF
A. minisclerotigenes    YFQVVSGVSAAQSGIRNLPLILAQSVSTVISGVTLSKFGYPQPFLLVGGVLTAIGSGLLY TLEVNTGSGKWIGYQLLAGIGIGWCFQVPVVTAQASVAPEDLPSVTAMVLTVQTLGGSVF
A. caelatus             YFQVVSGVSAAQSGIRNLPLILAQSASTVISGVTLSKFGYPQPFLLVGGVLTAVGSGLLY TLEVNTGSGKWIGYQLLAGIGIGWCFQVPLVTAQASVAPEDLPSVTAMVLTVQTLGGSVF
A. pseudocaelatus       YFQVVSGVSAAQSGIRNLPLILAQSASTVISGVTLSKFGYPQPFLLVGGVLTAIGSGLLY TLEVNTGSGKWIGYQLLAGIGIGWCFQVPLVTAQASVAPEDLPSVTAMVLTVQTLGGSVF
                        *****.****************.**. :.**** ****   *****:.*..*.::.***** **:*:* .***********:*:*****:*:****::* **.:.***********:***:*

H. pulvinata            VSAGQSALVNILFKSLPAGFSDSDAAAVANTAGEKLRETVSPEHFPAVLRAYMDGIRGAL LIAVVASGLAFVASLLLRSTKLRTGNSAAAAV
A. oryzae               VSAGQSAMVNVLLKNLRSETSTVDAVKVAMTGATELRQKFSSAQMPFILTAYMNGLQAAF LVAVVASSVATVVCLATRWIKFSGQTTAVAAA
A. parvisclerotigenus   VSAGQSAMVNVLLKNLRSETSTVDAVKVAMTGATELRQKFSSAQMPFILTAYMNGLQAAF LVAVVASSVATVVCLATRWIKFSGQTTAVAAA
A. flavus               VSAGQSAMVNVLLKNLRSETSTVDAVKVAMTGATELRQKFSSAQMPFILTAYMNGLQAAF LVAVVASSVATVVCLATRWIKFSGQTTAVAAA
A. minisclerotigenes    VSAGQSAMVNVLLKNLRSETSTVDAVKVAMTGATELRQKFSSAQMPFILTAYMNGLQAAF LVAVVASSVATVVCLATRWIKFSGKTTAAAAA
A. caelatus             VSAGQSAMVNVLLKNLRSETSTIDAVKVAMTGATELRQKFSSTQMPFILTSYMEGLQATF LVAVIASSVATVVCLATRWVKFSGQTTAAAAA
A. pseudocaelatus       VSAGQSAMVNVLLKNLRSETSTIDAVKVAMTGATELRQKFSSTQMPFILTSYMEGLQATF LVAVIASSVATVVCLATRWVKFSGQTTAAAAA
                        *******:**:*:*.* :   *  **. ** *.. :**:..*. ::* :* :**:*::.:: *:**:**.:* *..*  *  *:   .:*.**.
```

**Figure S6. Alignment of deduced amino acid sequences of *HpDPH6* homologs from *Hansfordia pulvinata* and six *Aspergillus* species.** The substrate binding pocket conserved in the transporters of the major facilitator superfamily is indicated in bold. Accession numbers and sequences are listed in Supplementary Table S2.

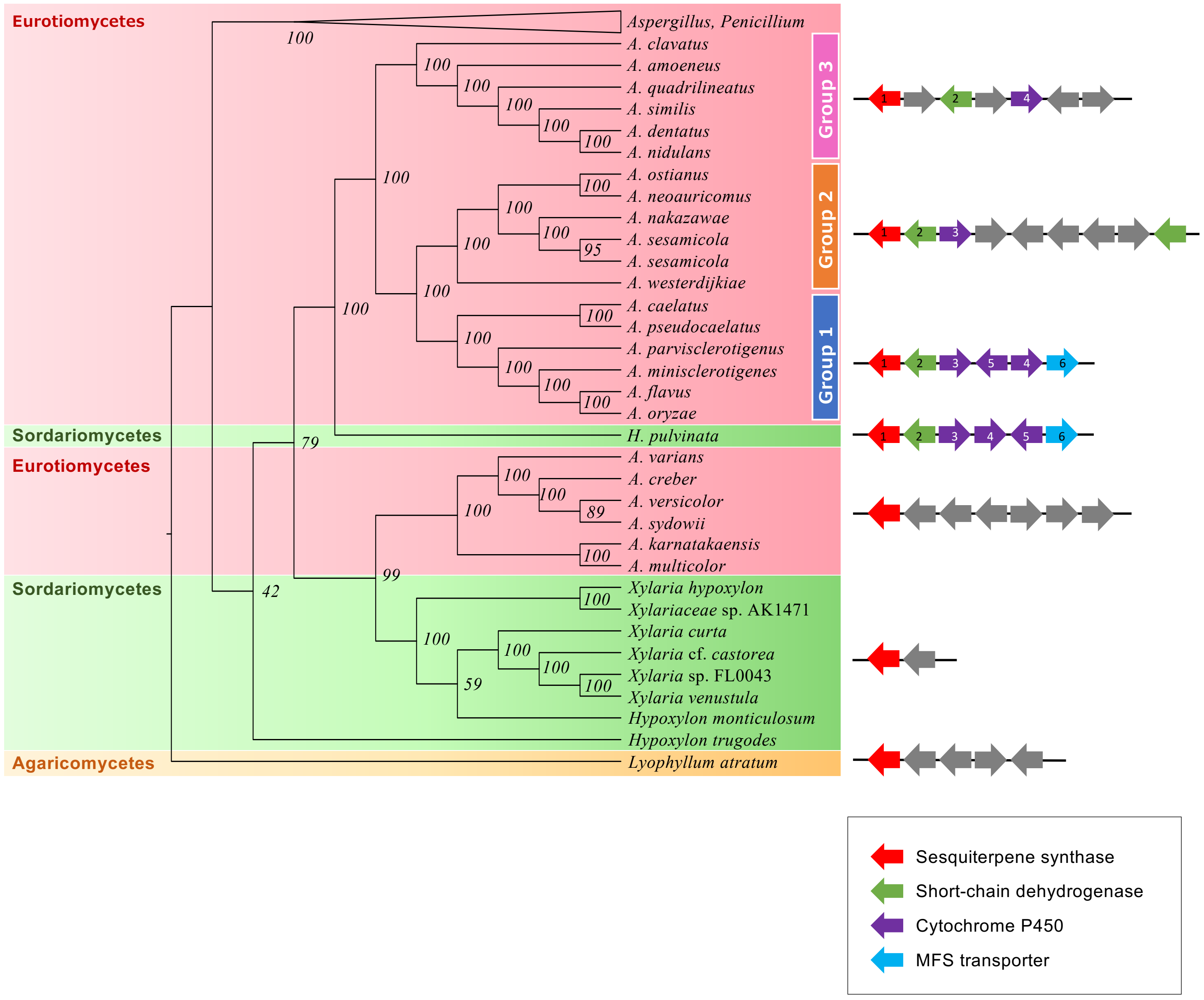


**Figure S7. Phylogenetic analysis of genomic sequences containing *HpDPH1* homologous genes and the upstream region using the maximum likelihood method.**

The 60 fungal genome sequences (approximately 15 kb) used in the phylogenetic analysis are listed in Supplementary Table S1. Numbers at branches represent bootstrap percentages (1000 replicates). The sequence of *Lyophyllum atratum* belonging to *Agaricomycetes* was used as an outgroup. Schematic diagrams of genomic structures on the right indicate gene orientation with annotation information in color; numbers in the arrows indicate genes that are more than 50% homologous to HpDph1 to HpDph6 proteins of *H. pulvinata*.

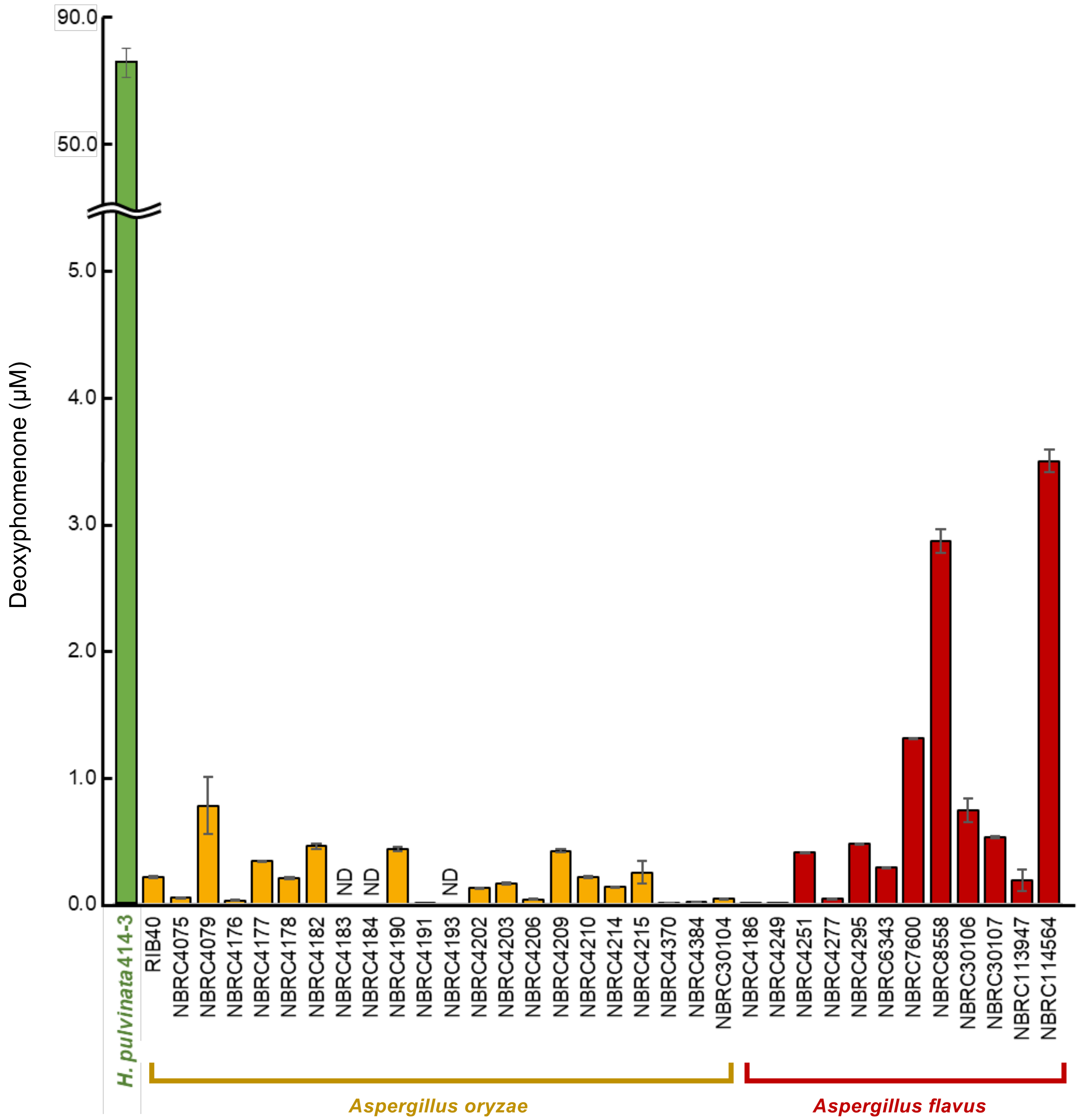


**Figure S8. Deoxyphomenone produced by *Hansfordia pulvinata* 414-3 and strains of *Aspergillus oryzae* and *A. flavus*.**

Strains were cultured in MM broth at 25 ºC. Deoxyphomenone in the culture filtrate was quantified using LC-MS/MS. Values are the means of three replicates (± SD). ND, not detected.

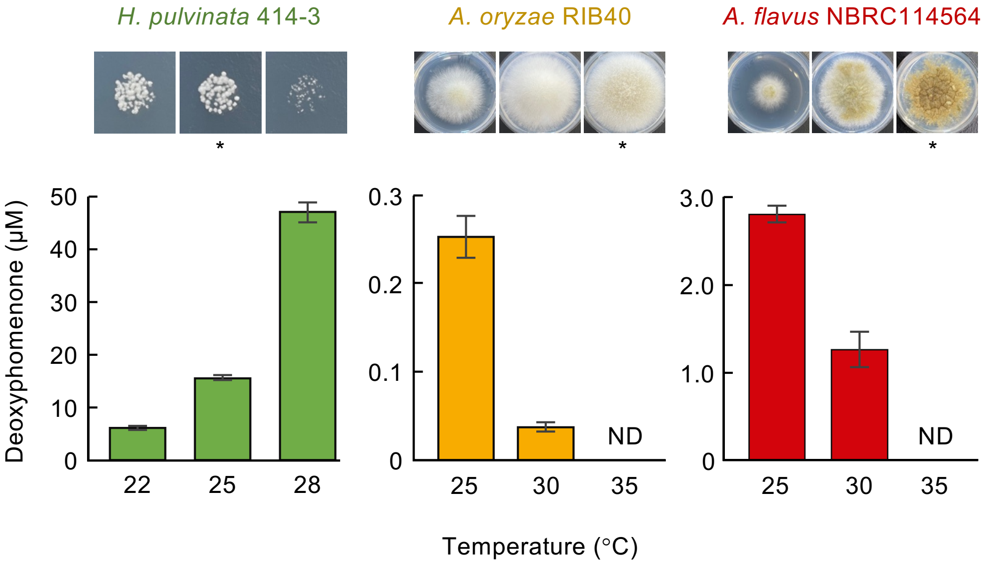


**Figure S9. Deoxyphomenone production by *Hansfordia pulvinata* 414-3, *Aspergillus oryzae* RIB40 and *A. flavus* NBRC114564 on MM agar or in broth at different temperatures.**
Deoxyphomenone in the culture filtrate of the broth was quantified by LC-MS/MS. The optimum temperature for growth on agar is indicated by the asterisk. Values are the means of three replicates (± SD). ND, not detected.

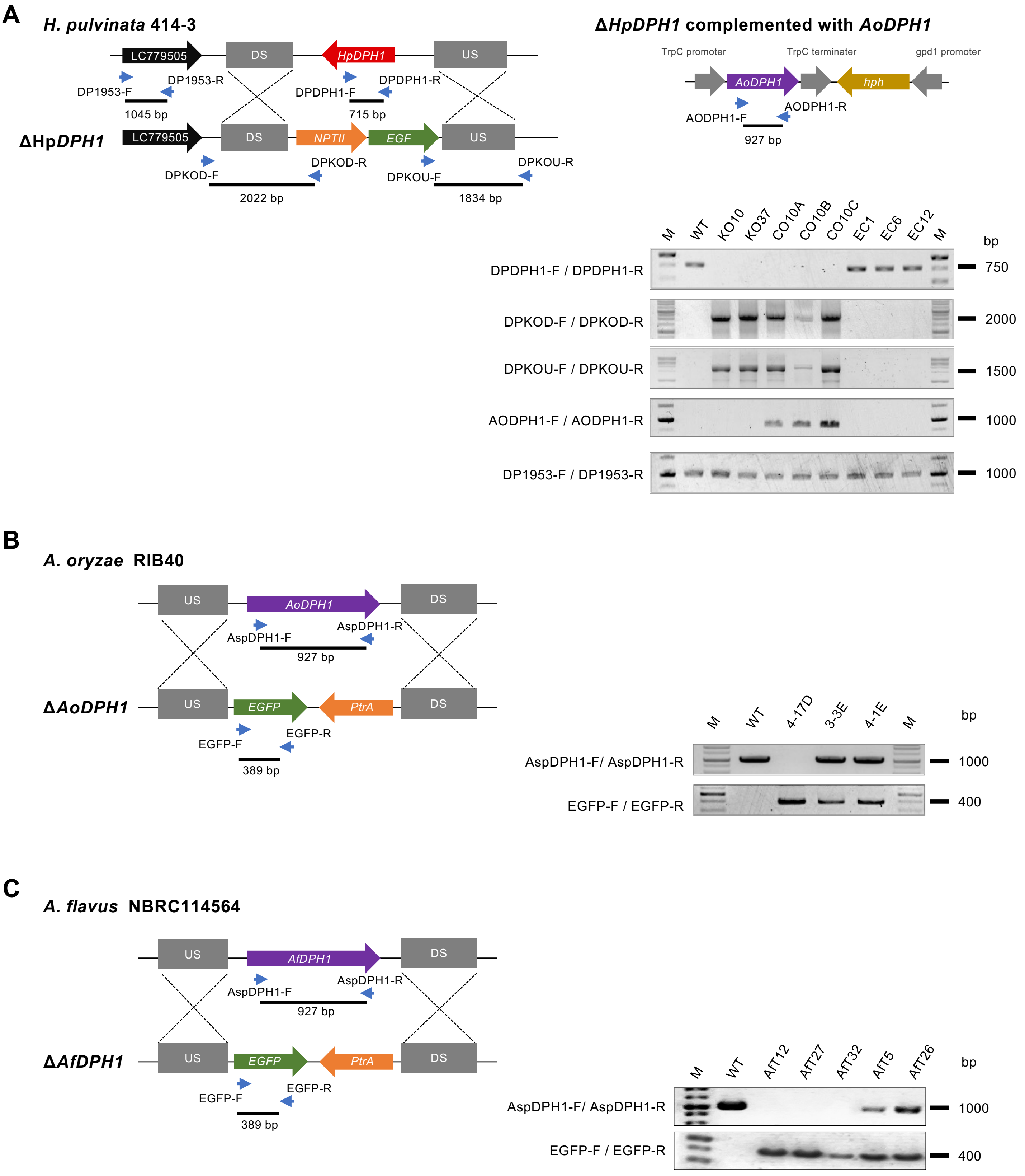


**Figure S10. PCR detection of the inserted cassettes in transformants and wild-type strains.**
Representation of the locus of *HpDPH1* homologous genes in wild-type strains of *Hansfordia pulvinata* 414-3 (A), *Aspergillus oryzae* RIB40 (B), *A. flavus* NBRC114564 (C) and transformants. Target genes were replaced by homologous recombination of the downstream (DS) and upstream (US) regions. *HpDPH1* knock-out mutant strain KO10 of *H. pulvinata* was complemented with functional *AoDPH1*. Large arrows: gene sequence; small blue arrows: primer sequence. The expected sizes of the amplicons are indicated under the primers. Each amplicon is shown on the right.

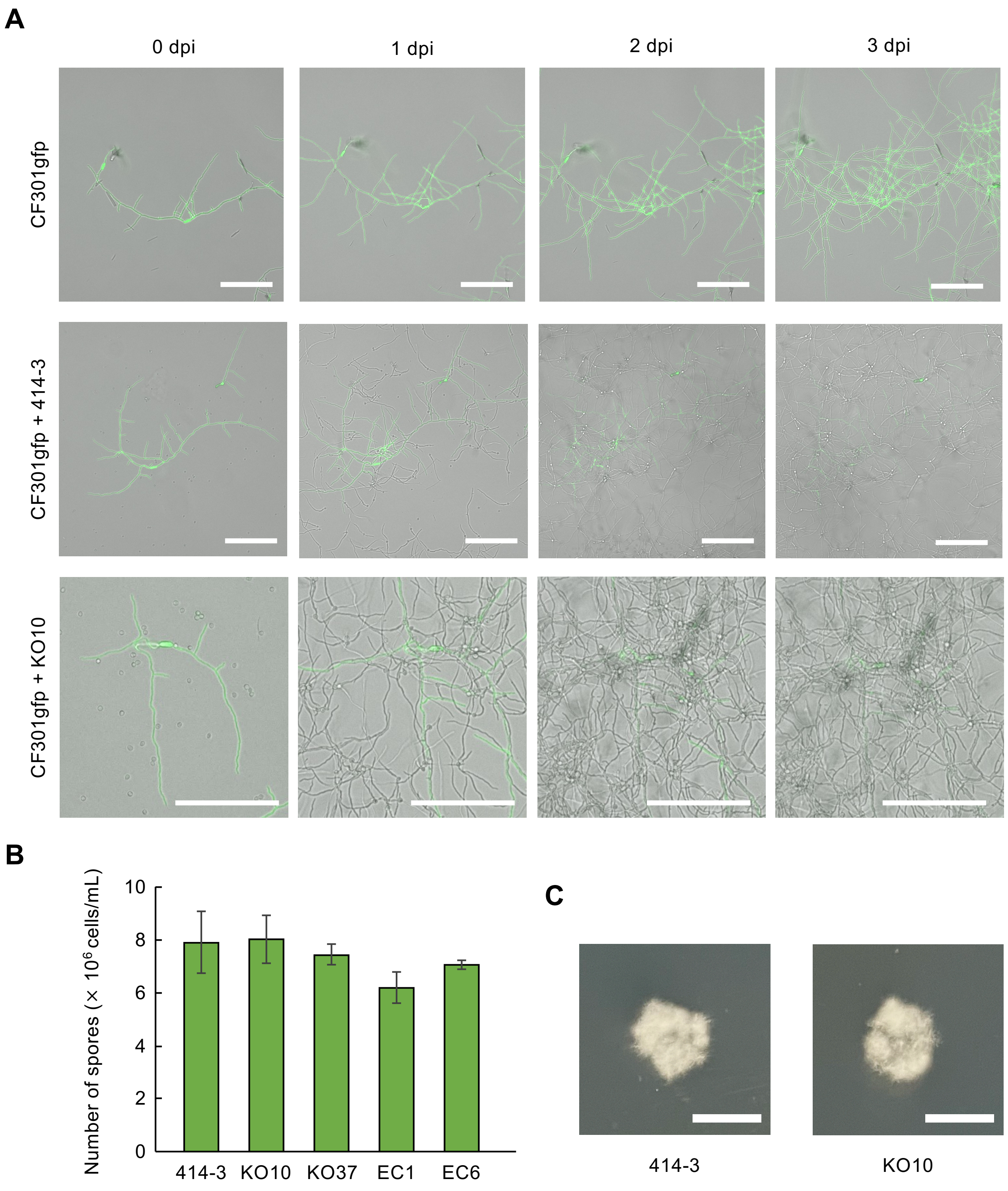


**Figure S11.** ***In vitro*** **assay of** ***Hansfordia pulvinata*** **mycoparasitic activity against GFP-expressing** ***Cladosporium fulvum.***
(A) *H. pulvinata* wild-type 414-3 or Δ*HpDPH1* mutant strain KO10 were cocultured with *C. fulvum* CF301gfp, which constitutively expresses GFP, in MM broth without a carbon source at 25 ºC. The parasitized *C. fulvum* cells lost GFP fluorescence. Bars indicate 100 µm. (B) Number of spores of 414-3, Δ*HpDPH1* mutants (KO10 and KO37) and ectopic strains (EC01 and EC06). Values are means of three biological replicates. Error bars indicate the standard deviation. (C) Mycelial growth of *H. pulvinata* 414-3 and KO10 strains on PDA.

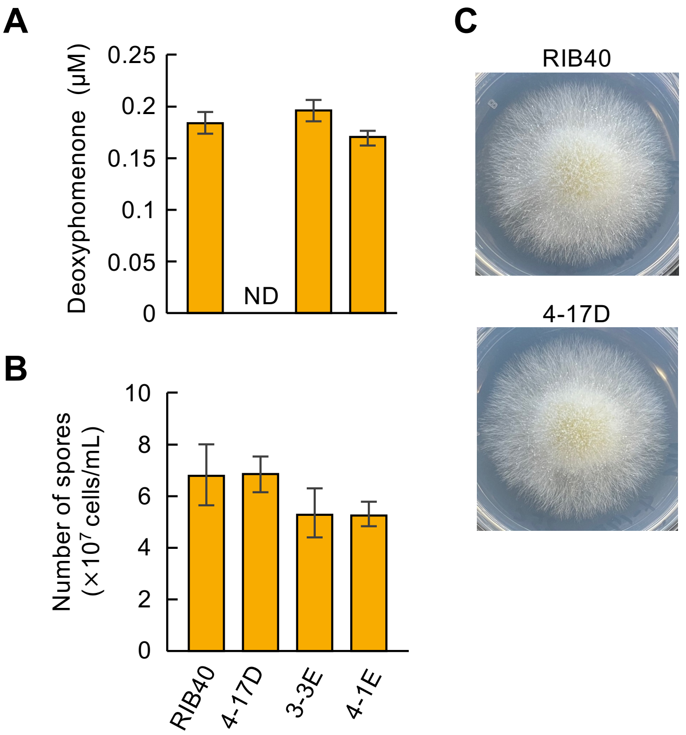


**Figure S12. Deoxyphomenone production and sporulation of *A. oryzae* RIB40 and transformants.** Wild-type RIB40, Δ*AoDPH1* mutant 4-17D, and ectopic strains 3-3E and 4-1E were cultured in MM broth or agar. Values in A and B are means of three replicates. Error bars indicate the standard deviation. (**A**) LC-MS/MS quantification of deoxyphomenone in culture filtrates. ND, not detected. (**B**) Number of spores formed on agar. (**C**) Colony morphology of wild-type RIB40 and Δ*AoDPH1* mutant 4-17D on MM agar 25°C.

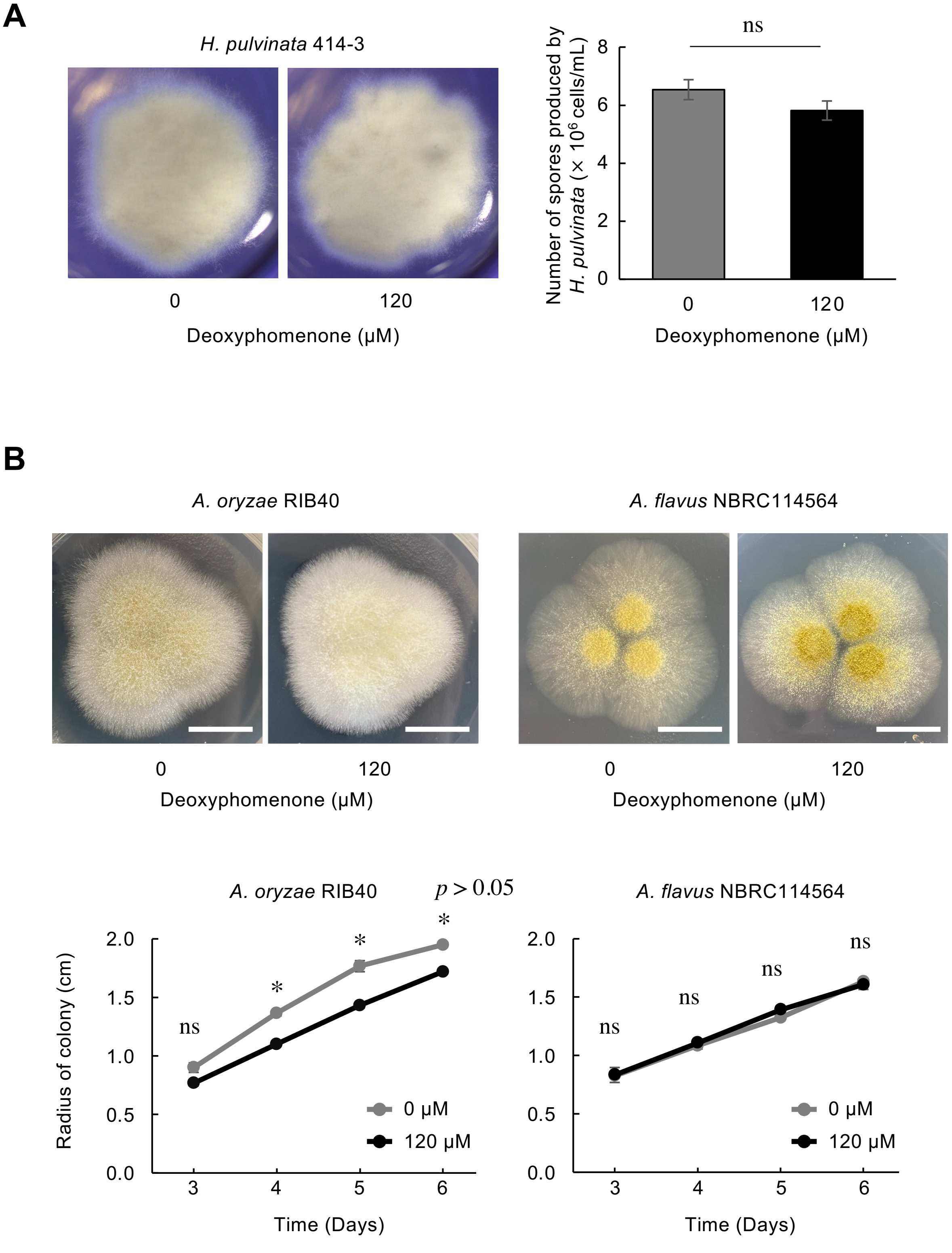


**Figure S13. Effect of deoxyphomenone on mycelial growth of *Hansfordia pulvinata*, *Aspergillus oryzae* and *A. flavus*.** Strains were grown on MM agar with 1% methanol (0 µM) or 120 µM deoxyphomenone. Values are means of three biological replicates (± SD). Means were compared for significant differences amount treatments using Tukey's test. ns, no significance. (**A**) Colony morphology and number of spores produced by *H. pulvinata* 414-3. (**B**) Colony of *A. oryzae* RIB40 and *A. flavus* NBRC114564. Bars = 1 cm. Fewer ocherous spores were produced by RIB40 in the presence of deoxyphomenone but more were produced by NBRC114564.

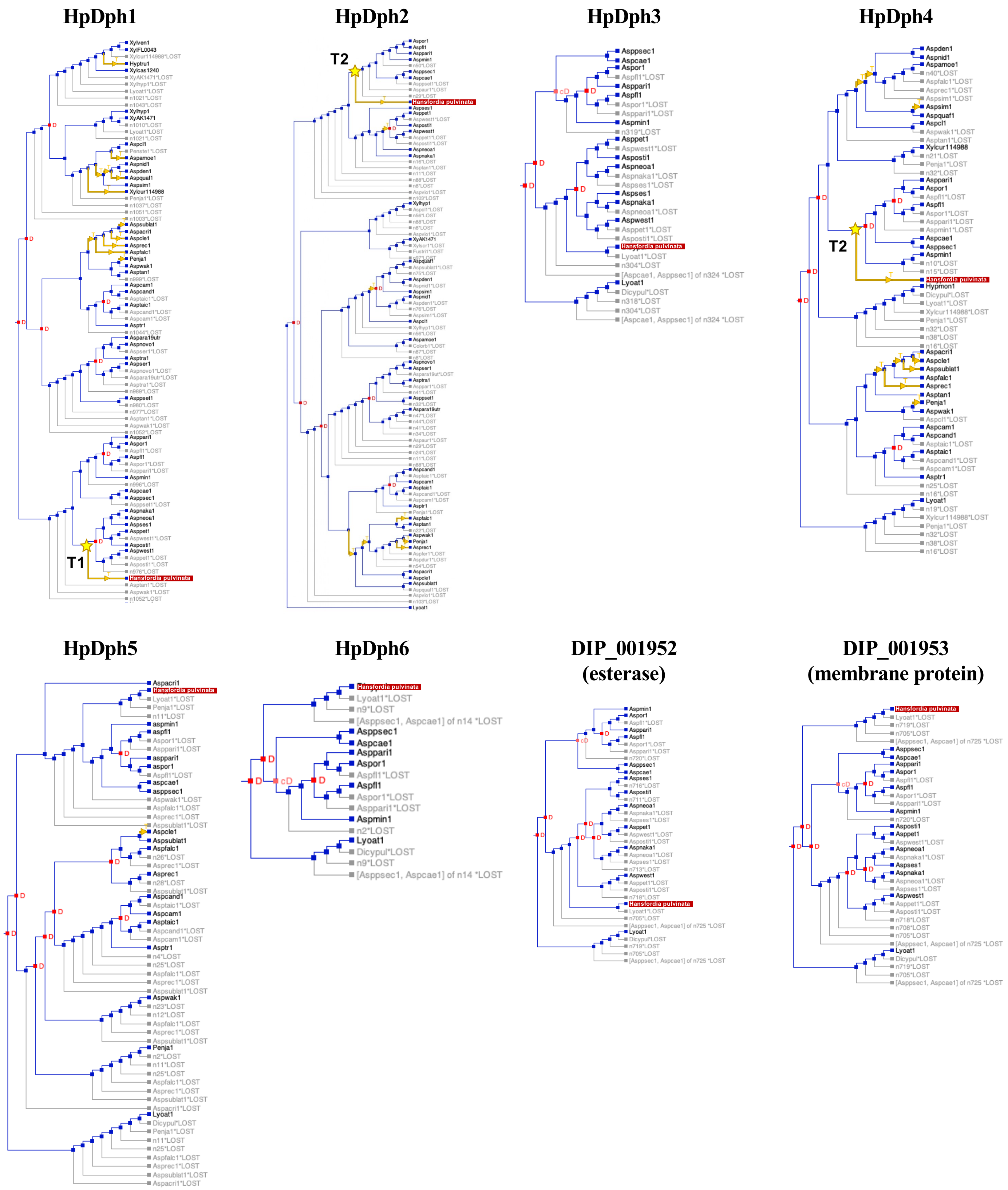


**Figure S14. Reconciliation of gene trees with species tree based on *DPH* nucleotide and amino acid sequences.** Phylogenetic protein trees were compared with the species tree using NOTUNG v.2.9 to infer duplications, losses, and horizontal transfer events for the HpDPH amino acid sequences. Duplication nodes are marked by red squares with a red D; losses are in grey; migrations are indicated with yellow arrows. Horizontal gene transfer events for the HpDph1 (T1) and HpDph2/HpDph4 (T2) proteins are indicated by yellow stars. *Hansfordia pulvinata* is highlighted by a red box. Event scores were calculated as total cost of duplications, transfers and losses. Costs/weights were set as duplications (D), 1.5; transfers (T), 8.0; losses (L), 1.0 (ratio D:T:L is 1:5.3:0.67).

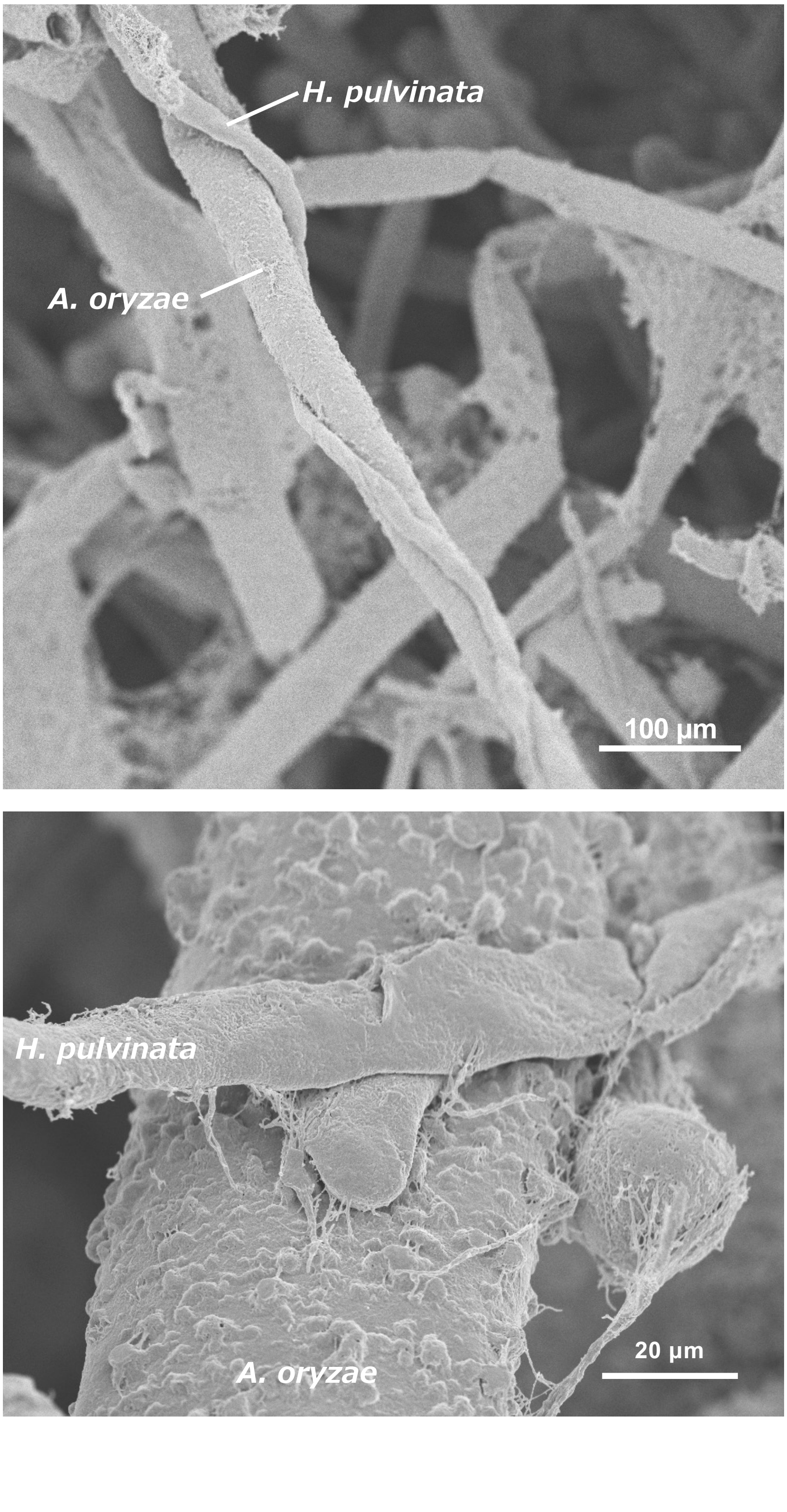


**Figure S15**. **Hyphal contact between the mycoparasite *Hansfordia pulvinata* 414-3 and *Aspergillus oryzae* RIB40.**

Coiling of hyphae of 414-3 around the thick, rough hyphae of RIB40 was rarely found.

**Supplemental table S1. Fungal genomic sequences used in this study**

| | Species name [a] | Abbreviation name in JGI | Class (kingdom) | Database | Nucleotide sequence |
|---|---|---|---|---|---|
| * | *Hansfordia pulvinata* 414-3 | - | Sordariomycetes (Ascomycota) | NCBI (ncbi.nlm.nih.gov/) | TCACTTGACCTTGTGGTACCGCGGCGTGGACTCGCTCCACCTCTCGTTGCCGCTCATCTG |
| * | *Aspergillus acristatulus* CBS 119.55 v1.0 | Aspacri1 | Eurotiomycetes (Ascomycota) | JGI (mycocosm.jgi.doe.gov) | CTAGCGCTGATTCGGCGAGTTGTAGCGCTGGGTCGTCAGACTCCATAACTCATTTCCAC |
| * | *Aspergillus amoeneus* CBS 111.32 v1.0 | Aspamoe1 | Eurotiomycetes (Ascomycota) | JGI (mycocosm.jgi.doe.gov) | CTACGCGCTGGCGGTTGCCTCCTTTTCAACATAACGCAGGGTCGTACGACTCCACAGCT |
| * | *Aspergillus arachidicola* v1.0 | Aspara19utr | Eurotiomycetes (Ascomycota) | JGI (mycocosm.jgi.doe.gov) | CTAGGCAGCACCAGGCGAATTGTATCGCAAAGTAGTCACACTCCACAGCTCATTACCA |
| * | *Aspergillus caelatus* CBS 763.97 v1.0 | Aspcae1 | Eurotiomycetes (Ascomycota) | JGI (mycocosm.jgi.doe.gov) | CTAAAAATGATAACGAGAAGTTGATTCACTCCAGCGCTCATTTCCGCTCATCTGAAACT |
| * | *Aspergillus campestris* IBT 28561 v1.0 | Aspcam1 | Eurotiomycetes (Ascomycota) | JGI (mycocosm.jgi.doe.gov) | GTTATATCGAAGAGTAGTCAAACTCCACAACTCATTCCCACTAATTTGTAATTCCAACC |
| * | *Aspergillus candidus* CBS 102.13 v1.0 | Aspcand1 | Eurotiomycetes (Ascomycota) | JGI (mycocosm.jgi.doe.gov) | CTAGTTAGGCGAGTTATAGCGAAGAGTAGTCAAACTCCACAACTCATTCCCACTAATCT |
| * | *Aspergillus clavatus* NRRL 1 from AspGD | Aspcl1 | Eurotiomycetes (Ascomycota) | JGI (mycocosm.jgi.doe.gov) | CTACGCACTAGCGGCTGCCTCTTTTTCAACATAGCGCAGGGTTGTCCGACTCCACAGCT |
| * | *Aspergillus cleistominutus* CBS 200.75 v1.0 | Aspcle1 | Eurotiomycetes (Ascomycota) | JGI (mycocosm.jgi.doe.gov) | CTAGCGCTGATTCGGCGAGTTGTAGCGCTGAGTCGTCAGACTCCATAACTCATTTCCAC |
| * | *Aspergillus creber* IBT 32277 v1.0 | Aspcreb1 | Eurotiomycetes (Ascomycota) | JGI (mycocosm.jgi.doe.gov) | TTAATATATCTTATACCGGCTTGTTGTGCCGGACCACTGCTCATTTCCACTTGCCATATAT |
| * | *Aspergillus dentatus CBS 114.63 v1.0* | Aspden1 | Eurotiomycetes (Ascomycota) | JGI (mycocosm.jgi.doe.gov) | CTACGCGCTAGCGGTTGCCTCCTTTTCAACATAGCGCAGGGTCGTACGACTCCACCGCT |
| * | *Aspergillus falconensis* CBS 271.91 v1.0 | Aspfalc1 | Eurotiomycetes (Ascomycota) | JGI (mycocosm.jgi.doe.gov) | GGAAGAAGCAGTAACAATCTATCACATTCCATATGAGCGCATTCTAGCCTTGATTCGGC |
| * | *Aspergillus flavus* NRRL3357 | Aspfl1 | Eurotiomycetes (Ascomycota) | JGI (mycocosm.jgi.doe.gov) | TATATCCTGCATGTGTTGGCATCGGAGGATAATCGTCTACCTGATGCCCCGAGGTCAGA |
| * | *Aspergillus karnatakaensis* CBS 102800 v1.0 | Aspkar1 | Eurotiomycetes (Ascomycota) | JGI (mycocosm.jgi.doe.gov) | TTCAACCCACCTAACTTCACGACACCCTGCTAAACCCAGCCCCCCTCCCGCTACATACCA |
| * | *Aspergillus minisclerotigenes* CBS 117635 v1.0 | Aspmin1 | Eurotiomycetes (Ascomycota) | JGI (mycocosm.jgi.doe.gov) | CTAAAAATGATAACGAGGAGTAGATTCACTCCAGCGCTCATTTCCACTCATCTGAAACT |
| * | *Aspergillus multicolor* v1.0 | Aspmul1 | Eurotiomycetes (Ascomycota) | JGI (mycocosm.jgi.doe.gov) | CTACTCAATTTGATACCGACTTGTCGTCCGACACCACTCCTCATTCCCACTAGCCAGATA |
| * | *Aspergillus nakazawae* v1.0 | Aspnaka1 | Eurotiomycetes (Ascomycota) | JGI (mycocosm.jgi.doe.gov) | TCAACGGTGGTAGCGAGGGGTCGATTCACTCCACCGCTCGTTGCCACTCATTTGGTCTTC |
| * | *Aspergillus neoauricomus* CBS112787 v1.0 | Aspneoa1 | Eurotiomycetes (Ascomycota) | JGI (mycocosm.jgi.doe.gov) | TCAACGGTGGTAGCGAGGGGTCGATTCACTCCACCGCTCGTTGCCACTCATTTGGTCTTC |
| * | *Aspergillus nidulans* | Aspnid1 | Eurotiomycetes (Ascomycota) | JGI (mycocosm.jgi.doe.gov) | CTACGCGCTAGCGGTTGCCTCCTTTTCAACATAGCGCAGGGTCGTACGACTCCACCGCT |
| * | *Aspergillus novoparasiticus* CBS 126849 v1.0 | Aspnovo | Eurotiomycetes (Ascomycota) | JGI (mycocosm.jgi.doe.gov) | CTAGGCAGCACCAGGCGAATTGTATCGCAAAGTAGTCACACTCCACAGCTCATTACCA |
| * | *Aspergillus oryzae* RIB40 | Aspor1 | Eurotiomycetes (Ascomycota) | JGI (mycocosm.jgi.doe.gov) | CTAAAAATGATAACGAGGAGTAGATTCACTCCAGCGCTCATTTCCACTCATCTGAAACT |
| * | *Aspergillus ostianus* v1.0 | Asposti1 | Eurotiomycetes (Ascomycota) | JGI (mycocosm.jgi.doe.gov) | TCAACGGTGGTAGCGAGGGGTCGATTCACTCCACCGCTCGTTGCCACTCATTTGGTCTTC |
| * | *Aspergillus parvisclerotigenus* CBS 121.62 v1.0 | Asppari1 | Eurotiomycetes (Ascomycota) | JGI (mycocosm.jgi.doe.gov) | CTAAAAATGATAACGAGGAGTAGATTCACTCCAGCGCTCATTTCCACTCATCTGAAACT |
| * | *Aspergillus petrakii* CBS 105.57 v1.0 | Asppet1 | Eurotiomycetes (Ascomycota) | JGI (mycocosm.jgi.doe.gov) | TATCTCGGCGATTCCTGAAATGGCAAGAGTTGACGCTGCCTAGACGGCCCCAAGGAAC |
| * | *Aspergillus pseudocaelatus* CBS 117616 v1.0 | Asppsec1 | Eurotiomycetes (Ascomycota) | JGI (mycocosm.jgi.doe.gov) | CTAAAAATGATAACGAGAAGTTGATTCACTCCAGCGCTCATTTCCGCTCATCTGAAACT |
| * | *Aspergillus pseudotamarii* CBS 117625 v1.0 | Asppset1 | Eurotiomycetes (Ascomycota) | JGI (mycocosm.jgi.doe.gov) | CTAGGTCGCACTAGGCGAATTGTACCGTAACGTAGTCACACTCCACAGCTCATTGCCGG |
| * | *Aspergillus quadrilineatus (floriformis)* CBS 937.73 v1.0 | Aspquaf1 | Eurotiomycetes (Ascomycota) | JGI (mycocosm.jgi.doe.gov) | CTACGCGCTAGCGGTTGCCTCCTTTTCAACATAGCGCAGGGTCGTACGACCCCACAGCT |
| * | *Aspergillus recurvatus* v1.0 | Asprec1 | Eurotiomycetes (Ascomycota) | JGI (mycocosm.jgi.doe.gov) | CTAGCGCTGATTCGGCGAGTTGTAGCGCTGAGTCGTCAGACTCCATAACTCATTTCCAC |
| * | *Aspergillus sergii* CBS 130017 v1.0 | Aspser1 | Eurotiomycetes (Ascomycota) | JGI (mycocosm.jgi.doe.gov) | CTAGGCAGCAGTAGGCGAATTGTATCGCAAAGTAGTCACACTCCACAGCTCATTACCA |
| * | *Aspergillus sesamicola* CBS 137324 v1.0 | Aspses1 | Eurotiomycetes (Ascomycota) | JGI (mycocosm.jgi.doe.gov) | TCAGGCTGCAATTGAGACATAGATGGACGCTACTATTGCTACTAGTACTCCGGAGTAGA |
| * | *Aspergillus similis* v1.0 | Aspsim1 | Eurotiomycetes (Ascomycota) | JGI (mycocosm.jgi.doe.gov) | GGGTTGACGTCGGGAGGAGAACAAGAAACACCTTTTGGTTTGCAACCACGTAGCTAGC |
| * | *Aspergillus sublatus* IBT 19356 v1.0 | Aspsublat1 | Eurotiomycetes (Ascomycota) | JGI (mycocosm.jgi.doe.gov) | CTAGCGCTGATTCGGCGAGTTGTAGCGCTGAGTCGTCAGACTCCATAACTCATTTCCAC |
| * | *Aspergillus sydowii* CBS 593.65 v1.0 | Aspsy1 | Eurotiomycetes (Ascomycota) | JGI (mycocosm.jgi.doe.gov) | ATACCGGCTCGTAGTCCGAGACCACTGCTCATTCCCACTCGCCATATATTTCAAGGCCTG |
| * | *Aspergillus taichungensis* IBT 19404 v1.0 | Asptaic1 | Eurotiomycetes (Ascomycota) | JGI (mycocosm.jgi.doe.gov) | TTAGTTAGGCGAGTTATAGCGAAGAGTAGTCAAACTCCACAACTCATTCCCACTAATCT |
| * | *Aspergillus tanneri* DTO 303-18 v1.0 | Asptan1 | Eurotiomycetes (Ascomycota) | JGI (mycocosm.jgi.doe.gov) | TTATGAGGTGACTCCTATTTGAGCTGGCTCTGCTCGGTACGAGATGAATTTGCATAGTA |
| * | *Aspergillus transmontanensis* CBS 130015 v1.0 | Asptra1 | Eurotiomycetes (Ascomycota) | JGI (mycocosm.jgi.doe.gov) | CTAGGCAGCACTAGGCGAATTGTATCGCAAAGTAGTCACACTCCACAGCTCATTACCA |
| * | *Aspergillus triticus* CBS266.81 v1.0 | Asptr1 | Eurotiomycetes (Ascomycota) | JGI (mycocosm.jgi.doe.gov) | CTAGTTAGGCGAGTTATAGCGAAGTGTAGTCAAGCTCCACAACTCATTCCCACTAATCT |
| * | *Aspergillus varians* CBS 505.65 v1.0 | Aspvar1 | Eurotiomycetes (Ascomycota) | JGI (mycocosm.jgi.doe.gov) | AGACTGCTAAGAGAGAGAATAGGATCTGTAGTCTAGGTATCCATGGATCTATATATTTA |
| * | *Aspergillus versicolor* v1.0 | Aspve1 | Eurotiomycetes (Ascomycota) | JGI (mycocosm.jgi.doe.gov) | CAACTTCCCCGACTCGCGCGAGTTCATCCCCGAGCGATGGCTAGACCCGGAAAAGCGT |
| * | *Aspergillus waksmanii* IBT 31900 v1.0 | Aspwak1 | Eurotiomycetes (Ascomycota) | JGI (mycocosm.jgi.doe.gov) | CCCATATGAACGCATCCTAGCGTTGATTCGGCGAGTTGTAGCGCTGAGTCGTCAGACTC |
| * | *Aspergillus westerdijkiae* CBS 112803 v1.0 | Aspwest1 | Eurotiomycetes (Ascomycota) | JGI (mycocosm.jgi.doe.gov) | TCAACGATGGTAGCGAGGGGTCGATTCACTCCACCGCTCGTTGCCACTCATTTGGTCTTC |
| * | *Penicillium antarcticum* DTO 356-E5 v1.0 | Penanta1 | Eurotiomycetes (Ascomycota) | JGI (mycocosm.jgi.doe.gov) | TTATTGCCGATATCTTGGAGTCCACTGACTCCATAGCTCATTTCCACGCATAAAACTTTC |
| * | *Penicillium atramentosum* RS17 v1.0 | Penatra1 | Eurotiomycetes (Ascomycota) | JGI (mycocosm.jgi.doe.gov) | TTATTGCCGATATCTTGGAGTCCACTGACTCCATAGCTCATTTCCACGCATAAAACTTTC |
| * | *Penicillium brasilianum* MG11 | Penbra1 | Eurotiomycetes (Ascomycota) | JGI (mycocosm.jgi.doe.gov) | TTACTTTTGGTAGCGAGGAGTCCACTGAGTCCATGTCTCCATTCCACTCATAAAGTATTC |
| * | *Penicillium coprophilum* IBT 31321 | Pencop1 | Eurotiomycetes (Ascomycota) | JGI (mycocosm.jgi.doe.gov) | GTCAGTCTCTTATTCGAGCAAAATGTTCATCGTGAGAGCGAAAGGTCAGGTTACTAATG |
| * | *Penicillium decumbens* IBT 11843 | Pendec1 | Eurotiomycetes (Ascomycota) | JGI (mycocosm.jgi.doe.gov) | CGGAAAACCAGGAGTCAGCTTCGTCGGATAGCTGAGTTTGTTAGATTGATGTGCATAGT |
| * | *Penicillium flavigenum* IBT 14082 | Penfla1 | Eurotiomycetes (Ascomycota) | JGI (mycocosm.jgi.doe.gov) | CTCCTGGTTGATATAGCGTCCATGCAGGTGACTACTCGTCGGCGACTCTTGTTGATATAA |
| * | *Penicillium janthinellum* ATCC 10455 v1.0 | Penja1 | Eurotiomycetes (Ascomycota) | JGI (mycocosm.jgi.doe.gov) | GTCCATTCCACCTCTTGGTATAGAGGCAGTTTGTGTATGCCTAGAGTGCAACTCAATCAT |
| * | *Penicillium nalgiovense* FM193 | Pennal1 | Eurotiomycetes (Ascomycota) | JGI (mycocosm.jgi.doe.gov) | GCATATAACTCATTCACTTCTATTGTAAAAAGTTCGTTGGACCAAAGAGGATTGCCTATA |
| * | *Penicillium steckii* IBT 24891 | Penste1 | Eurotiomycetes (Ascomycota) | JGI (mycocosm.jgi.doe.gov) | GGCCAGCATACTGTACATCGCCGTTGCGCAGCGCTATACCATCTTGGTGGAAACCAAAA |
| * | *Penicillium swiecickii* 182 6C1 v1.0 | Penswi1 | Eurotiomycetes (Ascomycota) | JGI (mycocosm.jgi.doe.gov) | GAGCTTTCAGCTCAAAAAGTGTTGGGACTAAAGTTTGATGACGGCTTGGTCAAAATCCT |
| * | *Hypoxylon monticulosum* FL0542 v1.0 | Hypmon1 | Sordariomycetes (Ascomycota) | JGI (mycocosm.jgi.doe.gov) | TCAACGACCTTCTGTAAAAGCAAGACCCCCACTCCTATTGTACCGTGGCGTTAGAAGAC |
| * | *Hypoxylon trugodes* CBS 135444 v1.0 | Hyptru1 | Sordariomycetes (Ascomycota) | JGI (mycocosm.jgi.doe.gov) | AATACCATACTTTCGGATTTCTACGCCATATTGTACTCTCTCGCATTGATAAACTTCTAGA |
| * | *Xylaria cf. castorea* CBS 124033 v1.0 | Xylcas1240 | Sordariomycetes (Ascomycota) | JGI (mycocosm.jgi.doe.gov) | CTAGGGGTAGGGGAGGTTAAATGCGGTGAAATCTTAATTTTCGTAGCGACCTCCGTAGA |
| * | *Xylaria curta* CBS 114988 v1.0 | Xylcur1149 | Sordariomycetes (Ascomycota) | JGI (mycocosm.jgi.doe.gov) | CTGTTATATGGTGTATATAAAAAGGCATGGTAATTATGCATGATCAGATCCTAAATACG |
| * | *Xylaria hypoxylon* OSC100004 v1.0 | Xylhyp1 | Sordariomycetes (Ascomycota) | JGI (mycocosm.jgi.doe.gov) | TCAGTTAACACTAGAGTAGCGAAGCGTGGTCTTGCTCCAAAGTTCATTACCACTCATCT |
| * | *Xylaria sp.* FL0043 v1.0 | XylFL0043 | Sordariomycetes (Ascomycota) | JGI (mycocosm.jgi.doe.gov) | CTAATAGCGAATAGAGCCCAGGCTATGCACATCTGCAGACGCCTCTTCCGGTGCCAGGT |
| * | *Xylariaceae sp.* AK1471 v1.0 | XyAK1471 | Sordariomycetes (Ascomycota) | JGI (mycocosm.jgi.doe.gov) | TCAGCTAACCTTAGAGTATCGAAGGGTGGTCTTGCTCCAGAGCTCATTGCCGCTCATCT |

| | | | | | |
|---|---|---|---|---|---|
| * | *Xylaria venustula* FL0490 v1.0 | Xylven1 | Sordariomycetes (Ascomycota) | JGI (mycocosm.jgi.doe.gov) | TTCCGGTGCCAAGTATCGCAGGGTAGTCCTGCTCCAGAGTTCATTTCCACACATTTGTAG |
| * | *Lyophyllum atratum* CBS 144462 v1.0 | Lyoat1 | Eubasidiomycetes (Basidiomycota) | JGI (mycocosm.jgi.doe.gov) | TCTCTATCTTTGCGTCAAATAGACATTGATCGCTAGCGTCACGGGCTACTACACAACCCG |
| | *Aspergillus albertensis* v1.0 | aspalbe | Eurotiomycetes (Ascomycota) | JGI (mycocosm.jgi.doe.gov) | GGACAACGAACTTAAGTCGACATGAGGTGGACGGGAGGCTGAAGAAGAAATGATCAG |
| | *Aspergillus alliaceus* CBS 536.65 v1.0 | Aspalli1 | Eurotiomycetes (Ascomycota) | JGI (mycocosm.jgi.doe.gov) | CTAGTCAACTACCACGCTATACCGCAACGTTGTCTGACTCCACAGCTCATTGCCACTCAT |
| | *Aspergillus amylovorus* CBS 600.67 v1.0 | Aspamy1 | Eurotiomycetes (Ascomycota) | JGI (mycocosm.jgi.doe.gov) | TCAGTCAGTCACCACGCTGTATCGCTGGGTTGTCCGACTCCAGAGCTCATTACCGCTCAT |
| | *Aspergillus assulatus* CBS 27911 v1.0 | Aspass1 | Eurotiomycetes (Ascomycota) | JGI (mycocosm.jgi.doe.gov) | TCATTTCCACTCATCTGGTATTCCAAGCCCTTGATATATAGCCTCAGATTCTCACTGCATG |
| | *Aspergillus aureoterreus* v1.0 | Aspaur1 | Eurotiomycetes (Ascomycota) | JGI (mycocosm.jgi.doe.gov) | TTCGTCGATCATCCAGTGCCACGAATGGATATACGCCATGGATGAATTCAGTTGGCTCGC |
| | *Aspergillus bertholletius* IBT 29228 v1.0 | Aspber1 | Eurotiomycetes (Ascomycota) | JGI (mycocosm.jgi.doe.gov) | CTAATCCACCAATAAATACCTCGGCGTTGTCTTACTCCACAGTTCGTTCCCGCTCATTTGC |
| | *Aspergillus biplanus* CBS 468.65 v1.0 | Aspbip1 | Eurotiomycetes (Ascomycota) | JGI (mycocosm.jgi.doe.gov) | TCATGCCTTGACAGCATAACGACCAGTAAGCTTGCTCCAAACCTCGTGACCGCTAGCGC |
| | *Aspergillus californicus* CBS 123895 v1.0 | Aspcalif1 | Eurotiomycetes (Ascomycota) | JGI (mycocosm.jgi.doe.gov) | TGTTTTGGTAAGTTAGAAAATGCATTCTTTGGTTCACGGTAGATGATATATCTAGAGTCC |
| | *Aspergillus crustosus* CBS 478.65 v1.0 | Aspcru1 | Eurotiomycetes (Ascomycota) | JGI (mycocosm.jgi.doe.gov) | ATGATCTCTTTCTCCCCTTTGCCAAGGAGGGGAGGAAGGGCGTCCGTGTTCTAATTCGAT |
| | *Aspergillus desertorum* CBS 653.73 v1.0 | Aspdese1 | Eurotiomycetes (Ascomycota) | JGI (mycocosm.jgi.doe.gov) | CTAGTCAACTACCACGCTATAACGCAACGTTGTCTGACTCCACAGCTCATTGCCACTCAT |
| | *Aspergillus dromiae* CBS 140633 v1.0 | Aspdro1 | Eurotiomycetes (Ascomycota) | JGI (mycocosm.jgi.doe.gov) | GTGCGATTCTTCCCTCTTCTACTGTTCAATTCAAGCAGAAACGAAAAGAGGAACTACAG |
| | *Aspergillus duricaulis* CBS 481.65 v1.0 | Aspdur1 | Eurotiomycetes (Ascomycota) | JGI (mycocosm.jgi.doe.gov) | TCAATCCGTACCCCGGCCATATCGCTCCGTGGTCCTGCTCCACAGCTCATTTCCGCTCATC |
| | *Aspergillus elegans* CBS 116.39 v1.0 | Aspele1 | Eurotiomycetes (Ascomycota) | JGI (mycocosm.jgi.doe.gov) | TCAACCGTGGTAGCGAGGGGTCGATTCGCTCCACCGCTCGTTGCCACTCATTTGATACTC |
| | *Aspergillus ferenczii* CBS 121594 v1.0 | Aspfer1 | Eurotiomycetes (Ascomycota) | JGI (mycocosm.jgi.doe.gov) | ACTGCTTACTGCTTACTGCTTCATAGTGGACAGGGGCAATCCCGCCACGTACATCCTGAA |
| | *Aspergillus flocculosus* v1.0 | Aspfloc1 | Eurotiomycetes (Ascomycota) | JGI (mycocosm.jgi.doe.gov) | TCAACAATGGTAGCGAGGAGTCGATTCACTCCACCGCTCATTGCCGCTCATGTGGTACTC |
| | *Aspergillus foveolata* CBS 279.81 v1.0 | Aspfove1 | Eurotiomycetes (Ascomycota) | JGI (mycocosm.jgi.doe.gov) | CTAGTCAACTACCGCGCTATAACGCAACGTTTTCCGACTCCACTGCTCATTGCCACTCATC |
| | *Aspergillus fructiculosus* v1.0 | Aspfru1 | Eurotiomycetes (Ascomycota) | JGI (mycocosm.jgi.doe.gov) | AAAAAAAAAAAGGAAATAGGAAGAAGCAGTAACAATCTATCACATTCCATATGAGCG |
| | *Aspergillus galapagensis* CBS 117522 v1.0 | Aspgal1 | Eurotiomycetes (Ascomycota) | JGI (mycocosm.jgi.doe.gov) | TGCCGCCCAGTGTGGTATTTGAAGTGTTAACACCCCTCACTGGAGGGATGCGTAATTCCC |
| | *Aspergillus granulosus* CBS 588.65 v1.0 | Aspgra1 | Eurotiomycetes (Ascomycota) | JGI (mycocosm.jgi.doe.gov) | CTAGACGGCAGCGCCAACCAAATTATTATAGCGCAGCGTCGTTCTACTCCACTCTTCATT |
| | *Aspergillus hiratsukae* CBS 294.93 v1.0 | Asphir1 | Eurotiomycetes (Ascomycota) | JGI (mycocosm.jgi.doe.gov) | TTAGAAACTCATTCTCTTGTATGGAAACACCTGCCGCTTGATTACTGCTTCATTATGCTAT |
| | *Aspergillus implicatus* CBS 484.95 v1.0 | Aspimp1 | Eurotiomycetes (Ascomycota) | JGI (mycocosm.jgi.doe.gov) | CTACAAGTCAAGATACAGCTCCGGATGATACCGGCGGCAGGCTGCCTGCCACAGCTCAT |
| | *Aspergillus indicus* v2.0 | Aspind2 | Eurotiomycetes (Ascomycota) | JGI (mycocosm.jgi.doe.gov) | CTAGTCAACTACCACGCTATACCGCAACGTTGTCTGACTCCACAGCTCATTGCCACTCAT |
| | *Aspergillus lucknowensis* CBS 449.75 v1.0 | Asppluc1 | Eurotiomycetes (Ascomycota) | JGI (mycocosm.jgi.doe.gov) | GGATCGCACATATATAGCCTGCACGCCGTGACAACCATCCCCCCGCCAAGGGAAGGCTC |
| | *Aspergillus muricatus* v1.0 | Aspmuri1 | Eurotiomycetes (Ascomycota) | JGI (mycocosm.jgi.doe.gov) | TTAAATATGGTACCGTGGAGTACTTTCGCTCCATCGCTCGTTTCCACTCATTTGAGCTTCC |
| | *Aspergillus neoechinulatus* CBS120.55 v1.0 | Aspne1 | Eurotiomycetes (Ascomycota) | JGI (mycocosm.jgi.doe.gov) | GAAGCAAGGGTGTTCGCGTTTTGATCCGGTAGGAGACCCGATCCCGAAGAGCAGACTTC |
| | *Aspergillus neoindicus* CBS 444.75 v1.0 | Aspneoi1 | Eurotiomycetes (Ascomycota) | JGI (mycocosm.jgi.doe.gov) | CTAGCTGGGACGATCAAAATTGTATCGCTGGGTGATCTTGCTCCAGAGCTCATTACCAGC |
| | *Aspergillus parasiticus* CBS 117618 v1.0 | Asppar1 | Eurotiomycetes (Ascomycota) | JGI (mycocosm.jgi.doe.gov) | CTAGGCAGCACTAGGCGAATTGTATCGTAAAGTAGTGACACTCCACAGCTCATTACCAG |
| | *Aspergillus pseudofelis* IBT 34107 v1.0 | Asppsf1 | Eurotiomycetes (Ascomycota) | JGI (mycocosm.jgi.doe.gov) | CTCAAAACAATACATGTGTCCGCCTATGTATTTCTTTAGTTATGGTTCGAGGCTACTTAAC |
| | *Aspergillus pseudoustus* CBS 123904 v1.0 | Asppust1 | Eurotiomycetes (Ascomycota) | JGI (mycocosm.jgi.doe.gov) | TTAGCCGACAACGCTATATCGCAACGTAGTCCGACTCCAGAGCTCATTCCCACTCATCTC |
| | *Aspergillus pulvericola* CBS 137327 v1.0 | Asppul1 | Eurotiomycetes (Ascomycota) | JGI (mycocosm.jgi.doe.gov) | GGTCAGGCTCTGACCGAAATTAGGTGCGAATATGACATCAGGGAGCTCAACAATGGTA |
| | *Aspergillus puulaauensis* IBT 32284 v1.0 | Asppuu1 | Eurotiomycetes (Ascomycota) | JGI (mycocosm.jgi.doe.gov) | TTAATAAATCTTATACCGGCTTGTTGTGCCGGACCACTGCTCATTTCCACTTGCCATATAT |
| | *Aspergillus roseoglobulosus* CBS112800 v1.0 | Aspros1 | Eurotiomycetes (Ascomycota) | JGI (mycocosm.jgi.doe.gov) | CTATTCTGACTGTAGCGATCGGCTATGATAACGAGGCGTAGATTCACTCCACCGCTCATT |
| | *Aspergillus siamensis* CBS 137452 v1.0 | Aspsia1 | Eurotiomycetes (Ascomycota) | JGI (mycocosm.jgi.doe.gov) | TTAAAGACTACTCTAAGTATGTCAATGGTTTAAAACTCCACAACATACCTCCCTGACTCC |
| | *Aspergillus silvaticus* CBS 128.55 v1.0 | Aspsil1 | Eurotiomycetes (Ascomycota) | JGI (mycocosm.jgi.doe.gov) | GGTCCAGTAGCTTAGGTAGGGAATCATGTTCAGGTACGAATTTGATATCTCGTTGTCGTC |
| | *Aspergillus spectabilis* CBS 429.77A v1.0 | Aspspe1 | Eurotiomycetes (Ascomycota) | JGI (mycocosm.jgi.doe.gov) | CTACTCAACCCGGTACCGGCTCGTTGTCCTTGACCACTCCTCATTTCCACTCGCCAGATAC |
| | *Aspergillus steynii* IBT 23096 v1.0 | Aspste1 | Eurotiomycetes (Ascomycota) | JGI (mycocosm.jgi.doe.gov) | CTACTCCAAATGATGATACCTCAAGGTAGTCTGGCTCCATCGCTCATTCCCACTCATCTGG |
| | *Aspergillus stercoraria* CBS 428.93 v1.0 | Aspstec1 | Eurotiomycetes (Ascomycota) | JGI (mycocosm.jgi.doe.gov) | CTAGTCAACTACCACGCTATAACGCAACGTTGTCTGACTCCACAGCTCATTGCCACTCAT |
| | *Aspergillus stella-maris* CBS 113639 v1.0 | Aspstel1 | Eurotiomycetes (Ascomycota) | JGI (mycocosm.jgi.doe.gov) | GTGGTAGCGAGGGGTCGATTCGCTCCACCGCTCGTTGCCACTCATTTGATACTCAAGTCC |
| | *Aspergillus subauricomus* CBS 638.78 v1.0 | Aspsub1 | Eurotiomycetes (Ascomycota) | JGI (mycocosm.jgi.doe.gov) | TCAGACATGATACCGCGGAGTGGATTCGCTCCAGAGTTCGTTCCCACTCATTTGATATTC |
| | *Aspergillus tennesseensis* IBT 32283 v1.0 | Aspten1 | Eurotiomycetes (Ascomycota) | JGI (mycocosm.jgi.doe.gov) | CCACATACTTAATATATCTTATACCGGCTTGTTGTGCCGGACCACTGCTCATTTCCACTTG |
| | *Aspergillus tetrazonus* CBS 591.65A v1.0 | Asptetr1 | Eurotiomycetes (Ascomycota) | JGI (mycocosm.jgi.doe.gov) | CTACTCAACCCGGTACCGGCTCGTTGTCCGACTCCACTCTTCATTCCCACTGGCCAGGTAC |
| | *Aspergillus thermomutatus* v1.0 | Aspthe1 | Eurotiomycetes (Ascomycota) | JGI (mycocosm.jgi.doe.gov) | TGTCGTGGTCGACTGTAGCATCTATCGGGAAGTCTTCGTATTCTGCCAGCAAGTCAGGAA |
| | *Aspergillus venezuelensis* CBS 868.97 v1.0 | Aspven1 | Eurotiomycetes (Ascomycota) | JGI (mycocosm.jgi.doe.gov) | CAACTTCCCCGACTCGCGCGAGTTCATCCCCGAGCGATGGCTAGACCCGGAAAAGCGTA |
| | *Aspergillus violaceofuscus* CBS 115571 v1.0 | Aspvio1 | Eurotiomycetes (Ascomycota) | JGI (mycocosm.jgi.doe.gov) | CTACAAGCTAGAGTCAGGCTCTTGATCGACATAGCGCAGGGTTGTCCGACTCCACAGCT |
| | *Aspergillus viridinutans* CBS 127.56 v1.0 | Aspvir1 | Eurotiomycetes (Ascomycota) | JGI (mycocosm.jgi.doe.gov) | TCAGCTCAGAACGAACTTTTGGAATTTAACCATACCTAAGGATAAAGCACCATACACGC |
| | *Aspergillus westlandensis* CBS 123905 v1.0 | Aspwes1 | Eurotiomycetes (Ascomycota) | JGI (mycocosm.jgi.doe.gov) | TTACACATGATAGCGCGGGGTGGATTCGCTCCACAGCTCGTTCCCAATCATTTGACATTC |
| | *Cladophialophora carrionii* CBS 160.54 | Claca1 | Eurotiomycetes (Ascomycota) | JGI (mycocosm.jgi.doe.gov) | CTAACCCACAGTCACGTCGTTATATCGCGGCGTAATGAGACTCCAGACTTCATTCCCGGC |
| | *Cladophialophora yegresii* CBS 114405 | Claye1 | Eurotiomycetes (Ascomycota) | JGI (mycocosm.jgi.doe.gov) | CTAGCCAACAGTCGCGTCATTATATCGTGGCGTGATGAGACTCCAGACCTCATTCCCGGC |
| | *Penicillium canescens* ATCC 10419 v1.0 | Penca1 | Eurotiomycetes (Ascomycota) | JGI (mycocosm.jgi.doe.gov) | GATATTCGGTGAGAAATATAGCGCAGTGAGATGTTTCTTTCCATAACAATTACATGGTTC |
| | *Penicillium chrysogenum* v1.0 | Pench1 | Eurotiomycetes (Ascomycota) | JGI (mycocosm.jgi.doe.gov) | TTAAAGGACGGGCTCATCTGAGACAGAATGGTATCTGAGTGTCGTCTGGCTCCACCTCTC |
| | *Penicillium digitatum* Pd1 | Pendig1 | Eurotiomycetes (Ascomycota) | JGI (mycocosm.jgi.doe.gov) | CTAGACCAGATTATTATATCGCAACGTGGTCTTACTCCACTCCTCGTTGCCACTCATCTGC |
| | *Penicillium expansum* d1 | Penexp1 | Eurotiomycetes (Ascomycota) | JGI (mycocosm.jgi.doe.gov) | CTAGACCACATTGTTATATCGTAAAGTGGTCTTACTCCACTCCTCGTTACCACTCATCTGC |
| | *Penicillium fellutanum* ATCC 48694 v1.0 | Penfe1 | Eurotiomycetes (Ascomycota) | JGI (mycocosm.jgi.doe.gov) | GTTGGTGAGGTCTATATCATGAGGAGTGACAAGCCATATATCGTATGAGGCAGGCTATC |
| | *Penicillium glabrum* DAOM 239074 v1.0 | Pengl1 | Eurotiomycetes (Ascomycota) | JGI (mycocosm.jgi.doe.gov) | TCATCGATAGCGTGGCGTCCATTGACTCCATAGCTCTATCCCGCTCATGAGATACTCAAC |
| | *Penicillium italicum* PHI-1 | Penita1 | Eurotiomycetes (Ascomycota) | JGI (mycocosm.jgi.doe.gov) | TCATTGTCGATACCGCGGAGTCCACTGACTCCATAACTCGATGCCGCACATAAAGTATTC |
| | *Penicillium polonicum* IBT 4502 | Penpol1 | Eurotiomycetes (Ascomycota) | JGI (mycocosm.jgi.doe.gov) | TCATTGTCGATATCGTGGAGTCCACTGACTCCATAACTCGATGCCGCACATAAAGTATTC |
| | *Penicillium raistrickii* ATCC 10490 v1.0 | Penra1 | Eurotiomycetes (Ascomycota) | JGI (mycocosm.jgi.doe.gov) | GTCATTCAAGCAGATGTATTGAAGAAAAACTCTTATCAAAGGCCTTCATCACGGATTCCC |
| | *Penicillium solitum* IBT 29525 | Pensol1 | Eurotiomycetes (Ascomycota) | JGI (mycocosm.jgi.doe.gov) | TTAGTTGTATCGACGCGTGGCCTTACTCCACTGCTCATTGCCGCTCATCTGGTGCTCTAGC |
| | *Penicillium sp.* | Pensp1 | Eurotiomycetes (Ascomycota) | JGI (mycocosm.jgi.doe.gov) | CTCGTAAATAGATGGTATTTCTATGACTCTCTGTGTGCTTCTTGGATATCCTTCTCGGTAC |
| | *Alternaria alternata* SRC1lrK2f v1.0 | Altal1 | Dothideomycetes (Ascomycota) | JGI (mycocosm.jgi.doe.gov) | CTACATCTGTACAGGATGGTACCTCGAAGTCGTCGAGCTCCACACTTCATTCCCACTCAT |
| | *Alternaria sp.* UNIPAMPA012 v1.0 | Altsp012 | Dothideomycetes (Ascomycota) | JGI (mycocosm.jgi.doe.gov) | CTACATCTGTACAGGATGGTACCTCGAAGTCGTCGAGCTCCACACTTCATTCCCACTCAT |

| | | | | |
|---|---|---|---|---|
| *Cercospora berteroae* CBS 538.71 | Cerbe1 | Dothideomycetes (Ascomycota) | JGI (mycocosm.jgi.doe.gov) | TTATGATCCAACTGCACCACCCAGCAGCTCTGGATGACCAGAATTGAACTCCTCCGCAT |
| *Cercospora zeae-maydis* v1.0 | Cerzm1 | Dothideomycetes (Ascomycota) | JGI (mycocosm.jgi.doe.gov) | TTATGATCCAACATTACCACTCAGCAGCCGCGGATTCGCGTAATTAAACTCCTCCTCATC |
| *Phyllosticta capitalensis* CBS 173.77 v2.0 | Phycapi2 | Dothideomycetes (Ascomycota) | JGI (mycocosm.jgi.doe.gov) | CTACCGGTCGCTTGACTCCGAAAATCCATAGCGGGGAGTGGTGATGCTCCAATGCAAA |
| *Phyllosticta citriasiana* CBS 120486 v1.0 | Phycit1 | Dothideomycetes (Ascomycota) | JGI (mycocosm.jgi.doe.gov) | CACCTTTCCCTGTCCCGCGCCCCGTCCACCACTTTTATCAAGCATCGCTTGCCCGTGCTGT |
| *Phyllosticta citricarpa* CBS 141352 v1.0 | Pcit141352 | Dothideomycetes (Ascomycota) | JGI (mycocosm.jgi.doe.gov) | TTACCCCCTCGACTCTGAAAATCCGTAGCGCGGAGTCGTGATGCTCCAATGCAAGTGGC |
| *Phyllosticta citricarpa* CBS 127454 v1.0 | Phycitr1 | Dothideomycetes (Ascomycota) | JGI (mycocosm.jgi.doe.gov) | TTACATGCCGGCAAAGGACGAACTGCTGCTCGACTCCGAAAATCCGTACCGCGGAGTT |
| *Phyllosticta sp.* CPC 27913 v1.0 | Phycpc1 | Dothideomycetes (Ascomycota) | JGI (mycocosm.jgi.doe.gov) | TGAAAATCCGTAGCGCGGAGTCGTGATGCTCCAATGCAAGTGGCCGCTCATGAGATCC |
| *Colletotrichum cereale* CBS 129662 v1.0 | Colce1 | Sordariomycetes (Ascomycota) | JGI (mycocosm.jgi.doe.gov) | CTAAGCCGTCGAAGGTGCGTTGTAGCGCAGTGTCGTCTTGCTCCAAAGCTCGTTCCCGC |
| *Colletotrichum eremochloae* CBS129661 v1.0 | Coler1 | Sordariomycetes (Ascomycota) | JGI (mycocosm.jgi.doe.gov) | TTTGAAGGCAGACTCAAATTCAAAATAAAAATAAAAACATCGTGAAGTGACCAGTCTT |
| *Colletotrichum godetiae* CBS 193.32 v1.0 | Colgo1 | Sordariomycetes (Ascomycota) | JGI (mycocosm.jgi.doe.gov) | CTAGACAGAGTGATATCTTGGAGTAATGCTGCTCCATGCCTCATTCCCACTCATTTGATA |
| *Colletotrichum orbiculare* 104-T | Colorb1 | Sordariomycetes (Ascomycota) | JGI (mycocosm.jgi.doe.gov) | CTAAGCGGTGGGAGGAGCATTGTACCGCAACGTGGTCTTGCTCCAAAGCTCGTTGCCGC |
| *Colletotrichum sublineola* CBS 131301 v1.0 | Colsu1 | Sordariomycetes (Ascomycota) | JGI (mycocosm.jgi.doe.gov) | ATACCGCGGAGTTGTCCGGCTCCAGAGCTCGTTTCCACTCATCTGGTACTCCAAGCCTTT |
| *Colletotrichum zoysiae* MAFF235873 v1.0 | Colzo1 | Sordariomycetes (Ascomycota) | JGI (mycocosm.jgi.doe.gov) | GTTCAGCACATTACATACGCTTCCGGTAGTCTTGATCCGTATAACTTAATTGCTTTCAAT |
| *Fusarium redolens* MPI-CAGE-AT-0023 v1.0 | Fusre1 | Sordariomycetes (Ascomycota) | JGI (mycocosm.jgi.doe.gov) | CTACTCAGAATAGCGACGTGTAGTCTTGCTCCAGAGCTCATTGCCACTCATCTGATACTC |
| *Fusarium avenaceum* MPI-SDFR-AT-0044 v1.0 | Fustri1 | Sordariomycetes (Ascomycota) | JGI (mycocosm.jgi.doe.gov) | TTACACGTTGTTATACCGAGGTGTCAACCTGCTCCATGCTTCATTGCCGCTCATTTGATAC |
| *Hypoxylon sp.* FL1284 v2.0 | HyFL1284 | Sordariomycetes (Ascomycota) | JGI (mycocosm.jgi.doe.gov) | CTGCATAATAAGCCACTTGCATGTAGGTACCTCTCAATGTTTGCATTTATTGGTAAGCCT |
| *Xylaria scruposa* CBS 123580 v1.0 | Xylscr1 | Sordariomycetes (Ascomycota) | JGI (mycocosm.jgi.doe.gov) | CGATGATACATGTACGATGCCCCTCATTACTCCAAAGGTGCTAGGTACCGCAATGTGGT |
| *Xylariaceae sp.* FL0662B v1.0 | XylFL0662B | Sordariomycetes (Ascomycota) | JGI (mycocosm.jgi.doe.gov) | GTGTAGAGTAAAATCACATATGAAGTAAGTTAAATACTAAACTATGAATGACCAGACA |

[a] Asterisks indicate sequences of 60 fungal species used in the phylogenetic analysis shown in supplemental figure S6.

**Supplemental table S2. Amino acid sequences of deoxyphomenone biosynthesis (*DPH*) genes from *Hansfordia pulvinata* and *Aspergillus* species**

| Organism | Gene function | DDBJ accession number or JGI protein ID [a] | Protein sequence [b] | Database |
|---|---|---|---|---|
| *Hansfordia pulvinata* 414-3 | Esterase (HP001952) | LC779504 | MDASTALGLIPTISRSFYGVAAASA | DDBJ/NCBI/EMBL |
| *Hansfordia pulvinata* 414-3 | Integral membrane ptorein (HP001953) | LC779505 | MAGEENKGPGFTAACIIVTVAAVI | DDBJ/NCBI/EMBL |
| *Hansfordia pulvinata* 414-3 | Sesquiterpene synthase (DPH1, HP001954) | LC779506 | MLSTLRSFELHKLLGLTGSSSNDST | DDBJ/NCBI/EMBL |
| *Hansfordia pulvinata* 414-3 | Short chain dehydrogenase (DPH2, HP001955) | LC779507 | MTTLRLSQAAIPDCTGKTVVITGGS | DDBJ/NCBI/EMBL |
| *Hansfordia pulvinata* 414-3 | CytochromeP450 (DPH3, HP001956) | LC779508 | MALITLALLCVAAWVLRRLGLAVY | DDBJ/NCBI/EMBL |
| *Hansfordia pulvinata* 414-3 | CytochromeP450 (DPH4, HP001957) | LC779509 | MAPRPTSGFWVLVFAVSWLGQVV | DDBJ/NCBI/EMBL |
| *Hansfordia pulvinata* 414-3 | CytochromeP450 (DPH5, HP001958) | LC779510 | MSLTVDLEALRWKLVPVLLAFLSW | DDBJ/NCBI/EMBL |
| *Hansfordia pulvinata* 414-3 | MFS transporter (DPH6, HP001959) | LC779511 | MATEKHPASLPGSQDAPPKTEAPP | DDBJ/NCBI/EMBL |
| *Aspergillus oryzae* RIB40 | Esterase | 10092* | MAGLWLGLGFTAGKSIAITLWSAIT | JGI (mycocosm.jgi.doe.gov) |
| *Aspergillus oryzae* RIB40 | Integral membrane ptorein | 10091* | MGAKENKGPGMTAASIVLTVIAVV | JGI (mycocosm.jgi.doe.gov) |
| *Aspergillus oryzae* RIB40 | Sesquiterpene synthase (AoDPH1) | 10090 | MLQRLWALSTSAIKLPFQPFSFGAP | JGI (mycocosm.jgi.doe.gov) |
| *Aspergillus oryzae* RIB40 | Short chain dehydrogenase (AoDPH2) | 10089* | MASMQLSDADIPSCAGKTVVITGG | JGI (mycocosm.jgi.doe.gov) |
| *Aspergillus oryzae* RIB40 | CytochromeP450 (AoDPH3) | 10088* | MTLISLSLLALSLWIIIRVLVIIYRLA | JGI (mycocosm.jgi.doe.gov) |
| *Aspergillus oryzae* RIB40 | CytochromeP450 (AoDPH5) | 10086 | MRSSTQLTALYWVHLVIYNVFFHP | JGI (mycocosm.jgi.doe.gov) |
| *Aspergillus oryzae* RIB40 | CytochromeP450 (AoDPH4) | 10087* | MDLFPRDYLFAGLAVFIFWWIIDHS | JGI (mycocosm.jgi.doe.gov) |
| *Aspergillus oryzae* RIB40 | MFS transporter (AoDPH6) | 10084 | MSTKTRSSQTGETAVSSRISTPATLI | JGI (mycocosm.jgi.doe.gov) |
| *Aspergillus flavus* NRRL3357 v1.0 | Esterase | 2229159 | MAGLWLGLGFTAGKSIAITLWSAIT | JGI (mycocosm.jgi.doe.gov) |
| *Aspergillus flavus* NRRL3357 v1.0 | Integral membrane ptorein | 2229158 | MGAKENKGPGMTAASIVLTVIAVV | JGI (mycocosm.jgi.doe.gov) |
| *Aspergillus flavus* NRRL3357 v1.0 | Sesquiterpene synthase | 1834709 | MLQRLWALSTSAIKLPFQPFSFGAP | JGI (mycocosm.jgi.doe.gov) |
| *Aspergillus flavus* NRRL3357 v1.0 | Short chain dehydrogenase | 2200580* | MASMQLSDADIPSCAGKTVVITGG | JGI (mycocosm.jgi.doe.gov) |
| *Aspergillus flavus* NRRL3357 v1.0 | CytochromeP450 | 2200579 | MTLISLSLLALSLWIIIRVLVIIYRLA | JGI (mycocosm.jgi.doe.gov) |
| *Aspergillus flavus* NRRL3357 v1.0 | CytochromeP450 | 2063894 | MRSSTQLTALYWIHLAIYNVFFHPI | JGI (mycocosm.jgi.doe.gov) |
| *Aspergillus flavus* NRRL3357 v1.0 | CytochromeP450 | 2062078 | MDLFPRDYLFAGLAVFIFWWIIDHS | JGI (mycocosm.jgi.doe.gov) |
| *Aspergillus flavus* NRRL3357 v1.0 | MFS transporter | 2210056 | MSTKTRSSQTGETAVSSRISTPATLI | JGI (mycocosm.jgi.doe.gov) |
| *Aspergillus parvisclerotigenus* CBS 121.62 v1.0 | Esterase | 166540 | MAGLWLGLGFTAGKSIAITLWSAIT | JGI (mycocosm.jgi.doe.gov) |
| *Aspergillus parvisclerotigenus* CBS 121.62 v1.0 | Integral membrane ptorein | 166547 | MGAKENKGPGMTAASIVLTVIAVV | JGI (mycocosm.jgi.doe.gov) |
| *Aspergillus parvisclerotigenus* CBS 121.62 v1.0 | Sesquiterpene synthase | 403455* | MLQRLWALSTSAIKLPFQPFSFGAP | JGI (mycocosm.jgi.doe.gov) |
| *Aspergillus parvisclerotigenus* CBS 121.62 v1.0 | Short chain dehydrogenase | | MASMQLSDADIPSCAGKTVVITGG | JGI (mycocosm.jgi.doe.gov) |
| *Aspergillus parvisclerotigenus* CBS 121.62 v1.0 | CytochromeP450 | 403457 | MTLISLSLLALSLWIIIRVLVIIYRLA | JGI (mycocosm.jgi.doe.gov) |
| *Aspergillus parvisclerotigenus* CBS 121.62 v1.0 | CytochromeP450 | 375701 | MRSSTQLTALYWVHLVIYNVFFHP | JGI (mycocosm.jgi.doe.gov) |
| *Aspergillus parvisclerotigenus* CBS 121.62 v1.0 | CytochromeP450 | 334960* | MDLFPRDYLFAGLAVFIFWWIIDHS | JGI (mycocosm.jgi.doe.gov) |
| *Aspergillus parvisclerotigenus* CBS 121.62 v1.0 | MFS transporter | 334970 | MSTKTRSSQTGETAVSSRISTPATLI | JGI (mycocosm.jgi.doe.gov) |
| *Aspergillus minisclerotigenes* CBS 117635 v1.0 | Esterase | 229809* | MAGLWLGLGFTAGKSIAITLWSAIT | JGI (mycocosm.jgi.doe.gov) |
| *Aspergillus minisclerotigenes* CBS 117635 v1.0 | Integral membrane ptorein | 242403 | MGAKENKGPGMTAASIVLTVIAVV | JGI (mycocosm.jgi.doe.gov) |
| *Aspergillus minisclerotigenes* CBS 117635 v1.0 | Sesquiterpene synthase | 229813* | MLQHLWAIATSAIKLPFQPFSFGAP | JGI (mycocosm.jgi.doe.gov) |
| *Aspergillus minisclerotigenes* CBS 117635 v1.0 | Short chain dehydrogenase | | MASMQLSDADIPSCAGKTVVITGG | JGI (mycocosm.jgi.doe.gov) |
| *Aspergillus minisclerotigenes* CBS 117635 v1.0 | CytochromeP450 | 251374 | MTLISLSLLALSLWIIIRVLVIIYRLA | JGI (mycocosm.jgi.doe.gov) |
| *Aspergillus minisclerotigenes* CBS 117635 v1.0 | CytochromeP450 | 229816 | MDRVVWLSLAVALTALYWVHLVI | JGI (mycocosm.jgi.doe.gov) |

| | | | | |
|---|---|---|---|---|
| *Aspergillus minisclerotigenes* CBS 117635 v1.0 | CytochromeP450 | 229815* | MDLLPRDYLFAGLAVFIFWWIIDH | JGI (mycocosm.jgi.doe.gov) |
| *Aspergillus minisclerotigenes* CBS 117635 v1.0 | MFS transporter | 216838 | MSTKTRSSHTGETAVSSRISTPATP | JGI (mycocosm.jgi.doe.gov) |
| *Aspergillus caelatus* CBS 763.97 v1.0 | Esterase | 160431 | MAGLWLGLGLTASKSVAVALWSA | JGI (mycocosm.jgi.doe.gov) |
| *Aspergillus caelatus* CBS 763.97 v1.0 | Integral membrane ptorein | 25776 | MGAKENKGPGMTAASIVLTVIAV | JGI (mycocosm.jgi.doe.gov) |
| *Aspergillus caelatus* CBS 763.97 v1.0 | Sesquiterpene synthase | 147588* | MLQRLWSLSTSAIKLPFQPVSFGA | JGI (mycocosm.jgi.doe.gov) |
| *Aspergillus caelatus* CBS 763.97 v1.0 | Short chain dehydrogenase | | MAAIQLSDADIPSCAGKVVVITGG | JGI (mycocosm.jgi.doe.gov) |
| *Aspergillus caelatus* CBS 763.97 v1.0 | CytochromeP450 | 132601 | MTLIYLSLLVLCLWIISRVLVIIYRL | JGI (mycocosm.jgi.doe.gov) |
| *Aspergillus caelatus* CBS 763.97 v1.0 | CytochromeP450 | 147591* | MVDRVVWLSLAVALIALYWANLV | JGI (mycocosm.jgi.doe.gov) |
| *Aspergillus caelatus* CBS 763.97 v1.0 | CytochromeP450 | 132604 | MDLFTRGYLLGGVVVCMFWWIVA | JGI (mycocosm.jgi.doe.gov) |
| *Aspergillus caelatus* CBS 763.97 v1.0 | MFS transporter | 160439 | MSTKTRSSQTGETAVSSGTSTPATL | JGI (mycocosm.jgi.doe.gov) |
| *Aspergillus pseudocaelatus* CBS 117616 v1.0 | Esterase | 290130 | MAGLWLGLGLTASKSVAVALWSA | JGI (mycocosm.jgi.doe.gov) |
| *Aspergillus pseudocaelatus* CBS 117616 v1.0 | Integral membrane ptorein | 91998 | MGAKENKGPGMTAASIVLTVIAV | JGI (mycocosm.jgi.doe.gov) |
| *Aspergillus pseudocaelatus* CBS 117616 v1.0 | Sesquiterpene synthase | 290130* | MLQRLWSLSTSAIKLPFQPVSFGA | JGI (mycocosm.jgi.doe.gov) |
| *Aspergillus pseudocaelatus* CBS 117616 v1.0 | Short chain dehydrogenase | | MAAIQLSDADIPSCAGKVVVITGG | JGI (mycocosm.jgi.doe.gov) |
| *Aspergillus pseudocaelatus* CBS 117616 v1.0 | CytochromeP450 | 305986 | MTLIYLSLLVLCLWIISRVLVIIYRL | JGI (mycocosm.jgi.doe.gov) |
| *Aspergillus pseudocaelatus* CBS 117616 v1.0 | CytochromeP450 | 278998* | MVDRVVWLSLAVALIALYWANLV | JGI (mycocosm.jgi.doe.gov) |
| *Aspergillus pseudocaelatus* CBS 117616 v1.0 | CytochromeP450 | 278999 | MDLFTREYLLGGVVVCMFWWIVA | JGI (mycocosm.jgi.doe.gov) |
| *Aspergillus pseudocaelatus* CBS 117616 v1.0 | MFS transporter | 290122 | MSTKTRSSQTGETAVSSGTSTHATI | JGI (mycocosm.jgi.doe.gov) |
| *Aspergillus nakazawae* v1.0 | Esterase | 234342 | MDQIRIGATVGKAVAAGVWGGLA | JGI (mycocosm.jgi.doe.gov) |
| *Aspergillus nakazawae* v1.0 | Integral membrane ptorein | 234343 | MENKGPGMIAASIVLTTVAFLLFCI | JGI (mycocosm.jgi.doe.gov) |
| *Aspergillus nakazawae* v1.0 | Sesquiterpene synthase | 234344 | MLATIWTALSKASATPKSAELSPIE | JGI (mycocosm.jgi.doe.gov) |
| *Aspergillus nakazawae* v1.0 | Short chain dehydrogenase | 234345 | MSSLTITDADIPDCSGKTVVITGGS | JGI (mycocosm.jgi.doe.gov) |
| *Aspergillus nakazawae* v1.0 | CytochromeP450 | 224099 | MAWFIVPVALLLLPAVIAVYRLVL | JGI (mycocosm.jgi.doe.gov) |
| *Aspergillus sesamicola* CBS 137324 v1.0 | Esterase | 344839* | MEQIRIGATVGKAVAAGVWGGLA | JGI (mycocosm.jgi.doe.gov) |
| *Aspergillus sesamicola* CBS 137324 v1.0 | Integral membrane ptorein | 301945 | MENNGPGMIAASIVLTTVAFLLFCI | JGI (mycocosm.jgi.doe.gov) |
| *Aspergillus sesamicola* CBS 137324 v1.0 | Sesquiterpene synthase | 301946 | MLATIWAALSKASATPKSAEFSPIE | JGI (mycocosm.jgi.doe.gov) |
| *Aspergillus sesamicola* CBS 137324 v1.0 | Short chain dehydrogenase | 344844 | MSTLTITDADIPDCSGKTFVITGGS | JGI (mycocosm.jgi.doe.gov) |
| *Aspergillus sesamicola* CBS 137324 v1.0 | CytochromeP450 | 344846 | MAWFIVPVALLLLPAVIAVYRLVL | JGI (mycocosm.jgi.doe.gov) |
| *Aspergillus neoauricomus* CBS112787 v1.0 | Esterase | 169847 | MEQIRIGATVGKAVAAGVWGGLA | JGI (mycocosm.jgi.doe.gov) |
| *Aspergillus neoauricomus* CBS112787 v1.0 | Integral membrane ptorein | 141434 | MENKGPGMIAASIVLTTVAFLLFCI | JGI (mycocosm.jgi.doe.gov) |
| *Aspergillus neoauricomus* CBS112787 v1.0 | Sesquiterpene synthase | 141433 | MLATIWTALSKASATPKSAELSPIE | JGI (mycocosm.jgi.doe.gov) |
| *Aspergillus neoauricomus* CBS112787 v1.0 | Short chain dehydrogenase | 141432 | MSSLTITDADIPDCSGKTVVITGGS | JGI (mycocosm.jgi.doe.gov) |
| *Aspergillus neoauricomus* CBS112787 v1.0 | CytochromeP450 | 141431 | MKMAWFIVPVALLLLPAVIAVYRL | JGI (mycocosm.jgi.doe.gov) |
| *Aspergillus ostianus* v1.0 | Esterase | 185058 | MDQIRIGATVGKAVAIGVWGGLA | JGI (mycocosm.jgi.doe.gov) |
| *Aspergillus ostianus* v1.0 | Integral membrane ptorein | 185057 | MENKGPGMIAASIVLTTVAFLLFCI | JGI (mycocosm.jgi.doe.gov) |
| *Aspergillus ostianus* v1.0 | Sesquiterpene synthase | 185056 | MLATIWTALSKASVTPKSADLSPID | JGI (mycocosm.jgi.doe.gov) |
| *Aspergillus ostianus* v1.0 | Short chain dehydrogenase | 185055 | MASLTITDADIPDCSGKTVVITGGS | JGI (mycocosm.jgi.doe.gov) |
| *Aspergillus ostianus* v1.0 | CytochromeP450 | 185054 | MQAGLGDGPKWSGPSIRLSARAIT | JGI (mycocosm.jgi.doe.gov) |
| *Aspergillus petrakii* CBS 105.57 v1.0 | Esterase | 285361 | MDQIRIGATVGKAVAAGVWGGLA | JGI (mycocosm.jgi.doe.gov) |
| *Aspergillus petrakii* CBS 105.57 v1.0 | Integral membrane ptorein | 285360 | MENKGPGMIAASIVLTTVAFLLFCI | JGI (mycocosm.jgi.doe.gov) |
| *Aspergillus petrakii* CBS 105.57 v1.0 | Sesquiterpene synthase | 285359 | MLATIWTALSKASATPKSADVSPIE | JGI (mycocosm.jgi.doe.gov) |
| *Aspergillus petrakii* CBS 105.57 v1.0 | Short chain dehydrogenase | 285358 | MASLTITDADIPDCSGKTVVITGGS | JGI (mycocosm.jgi.doe.gov) |
| *Aspergillus petrakii* CBS 105.57 v1.0 | CytochromeP450 | 296418 | MTMAWFIVPVALLLLPVAIAVYRL | JGI (mycocosm.jgi.doe.gov) |

| | | | | |
|---|---|---|---|---|
| *Aspergillus westerdijkiae* CBS 112803 v1.0 | Esterase | 289693 | MDQIRIGATVGKAVAAGVWGGLA | JGI (mycocosm.jgi.doe.gov) |
| *Aspergillus westerdijkiae* CBS 112803 v1.0 | Integral membrane ptorein | 265657 | MENKGPGMIAASIVLTTVAFILFCL | JGI (mycocosm.jgi.doe.gov) |
| *Aspergillus westerdijkiae* CBS 112803 v1.0 | Sesquiterpene synthase | 265656 | MLATIWTALSKASPTPKSAELSPIEI | JGI (mycocosm.jgi.doe.gov) |
| *Aspergillus westerdijkiae* CBS 112803 v1.0 | Short chain dehydrogenase | 265655* | MASLTITDADIPDCSGKTVVITGGS | JGI (mycocosm.jgi.doe.gov) |
| *Aspergillus westerdijkiae* CBS 112803 v1.0 | CytochromeP450 | 289683 | MAWLIVSVALLLLPVVIAVYRLVLI | JGI (mycocosm.jgi.doe.gov) |
| *Aspergillus amoeneus* CBS 111.32 v1.0 | Sesquiterpene synthase | 227190 | MPSAIDTDAISTQLLLNGVAKRHS( | JGI (mycocosm.jgi.doe.gov) |
| *Aspergillus amoeneus* CBS 111.32 v1.0 | Short chain dehydrogenase | 227192 | MTSLDLSADDIPRLEGRTAIITGGCS | JGI (mycocosm.jgi.doe.gov) |
| *Aspergillus amoeneus* CBS 111.32 v1.0 | CytochromeP450 | 227194 | MSASWVEQPNSLLDISKVVFVLTL | JGI (mycocosm.jgi.doe.gov) |
| *Aspergillus dentatus* CBS 114.63 v1.0 | Sesquiterpene synthase | 315840 | MPSAIDTDAISTQLLLNGVGKSHSI | JGI (mycocosm.jgi.doe.gov) |
| *Aspergillus dentatus* CBS 114.63 v1.0 | Short chain dehydrogenase | 315842 | MTSLNLSVDDIPRLDGKTAIITGGC | JGI (mycocosm.jgi.doe.gov) |
| *Aspergillus dentatus* CBS 114.63 v1.0 | CytochromeP450 | 315844 | MSASWVDEPSSLLGIFKVVFVLKK | JGI (mycocosm.jgi.doe.gov) |
| *Aspergillus nidulans* | Sesquiterpene synthase | 8825 | MPSAIDTDAISTQLLLNGVAKSHSI | JGI (mycocosm.jgi.doe.gov) |
| *Aspergillus nidulans* | Short chain dehydrogenase | 8823 | MYICPFRDTTNHRFLGGCSGIGWE | JGI (mycocosm.jgi.doe.gov) |
| *Aspergillus nidulans* | CytochromeP450 | 8821 | MSASWVDEPSSLLGIFEVVFVLKRI | JGI (mycocosm.jgi.doe.gov) |
| *Aspergillus similis* v1.0 | Sesquiterpene synthase | 210129 | MPSAIDTDAISTQLLLNGVAKSHSI | JGI (mycocosm.jgi.doe.gov) |
| *Aspergillus similis* v1.0 | Short chain dehydrogenase | 175589 | MTSLNLSVDDIPRLDGKTAIITGGC | JGI (mycocosm.jgi.doe.gov) |
| *Aspergillus similis* v1.0 | CytochromeP450 | 210125 | MSASWVDEPSSLLGIFKVVFVLKK | JGI (mycocosm.jgi.doe.gov) |
| *Aspergillus quadrilineatus* (*floriformis*) CBS 937.7. | Sesquiterpene synthase | 144829 | MPSAIDTDAISTQLLLNGVAKSHSI | JGI (mycocosm.jgi.doe.gov) |
| *Aspergillus quadrilineatus* (*floriformis*) CBS 937.7. | Short chain dehydrogenase | 144831 | MTSLNLSVDDIPRLDGKTAIITGGC | JGI (mycocosm.jgi.doe.gov) |
| *Aspergillus quadrilineatus* (*floriformis*) CBS 937.7. | CytochromeP450 | 145836 | MSASWVDEPSSLLGIFKVVFVLKK | JGI (mycocosm.jgi.doe.gov) |
| *Aspergillus clavatus* NRRL 1 from AspGD | Sesquiterpene synthase | 3242 | MPSAIDTGAISTQLLLNGAIKSHSG | JGI (mycocosm.jgi.doe.gov) |
| *Aspergillus clavatus* NRRL 1 from AspGD | Short chain dehydrogenase | 3244 | MTSLNLSVDDIPRLEGKTAIITGIFY | JGI (mycocosm.jgi.doe.gov) |
| *Aspergillus clavatus* NRRL 1 from AspGD | CytochromeP450 | 3246 | MSASWGYEPTSWLDIFTVVFVLVT | JGI (mycocosm.jgi.doe.gov) |
| *Lyophyllum atratum* CBS 144462 v1.0 | Sesquiterpene synthase | 1784547 | MPEPPSSFIAHHPSTPQFNPLLDPPI | JGI (mycocosm.jgi.doe.gov) |

[a] Asterisks indicate that the sequence information was reanalyzed based on the sequence of *A. oryzae* and re-annotated as protein seqneces. 403455, 229813, 147588, and 290130 were registered as one amino acid sequece consisting of *DPH1* and *DPH2* homologous genes. *DPH5* homologous sequences of *Aspergillus* species registered in JGI were reannotated based on the DIP_001958 sequence.

**Supplemental table S3. Primer sequences used in this study**

| Primer name | Primer sequence (5'-3') | Purpose |
|---|---|---|
| Dpprx2LF3 | ACCATGATTACGCCAGGAAGGTGGTCGTTACGAGT | Amplification of the upstream of *DPH1* gene for pGW43_dpprx2ko |
| Dpprx2LR3 | TAGGGAACTGCCAGGACTGTGCCTATGCCTGTGTG | |
| Dpprx2UF3 | AGCTGTTTCCTGGCAGGATGTAAAGAGGGCAGTCG | Amplification of the downstream of *DPH1* gene for pGW43_dpprx2ko |
| Dpprx2UR3 | GGCCAGTGAATTATCAAAGGCAATGCAATGCGAAG | |
| pRM254 attL1_F | CCTGGCAGTTCCCTACTCTC | Amplification of *gen* cassettes for pGW43_dpprx2ko |
| pRM254 attL2_R | TGCCAGGAAACAGCTATGAC | |
| pPM43GW_RB_F | TTGATAATTCACTGGCCGTC | Linearization of pPM43GW |
| pPM43GW_LB_R | TGGCGTAATCATGGTCATAG | |
| DPKOD-F | ACTACCACCACCACGACAGC | Screening for *DPH1* knocked-out mutants |
| DPKOD-R | AATTACACCCTTTGCGCCCT | |
| DPKOU-F | ATTGAATCCTGTTGCCGGTC | |
| DPKOU-R | ACGACATCGCCATCAACACG | |
| pPTREXeGFP_F | ATTCTAGAAGTCCTGAATAGTAGTTTGTGG | Amplification of *EGFP* cassettes |
| pPTREXeGFP_R | CCATTGGTAACGAAATGTAAAAGCTAGG | |
| AoDPH1_LB_F | CGAGCAGCTGAAGCTATCTCGGCGACACTCTCATTTGA | Amplification of *AoDPH1* gene |
| AoDPH1_LB_R | CAGGACTTCTAGAATTGTGAGCCTTGACAGCTGGATATC | |
| AoDPH1_RB_F | TTTCGTTACCAATGGCTCACACGCACCTCGTGCTGC | |
| AoDPH1_RB_R | GGAGACCGGCAGATCATTTAAATGATACGTGATATGCTCGTCGATCG | |
| AspDPH1_F | AACCTCAGCTATCAAGCTACCG | Detection of *AoDPH1/AfDPH1* gene |
| AspDPH1_R | GTCACGAGCTGTTTATGTACCG | |
| EGFP_F | TGACCCTGAAGTTCATCTGCAC | Detection of insertion sequence |
| EGFP_R | GATGTTGTGGCGGATCTTGAAG | |
| DP1953_F | AGAAGGGCTGAAACGGTAGCTT | Detection of DIP_001953 |
| DP1953_R | GAATCCTGGAGATGCTGACGAT | |
| DPDPH1-F | AGCTGTAGATGTCGTTGACC | Detection of *DPH1* gene |
| DPDPH1-R | GCCTTCTCACGCTGCTGTTTCT | |
| DpACTrt_F1n | AGGACTCTTACGTCGGTGAC | qPCR detection of *H. pulvinata* actin gene |
| DpACTrt_R1n | TCCATGTCATCCCAGTTGGT | |
| Dp1952rt_F1i | GACAGCCCTGGGTCTTATCC | qPCR detection of DIP_001952 |
| Dp1952rt_R1i | CCGGTGGGCTTTACCCATTA | |
| Dp1953rt_F1i | CTCGTGAACCAGCGCAAATA | qPCR detection of DIP_001953 |
| Dp1953rt_R1i | GCGCACTTGATGAGGAACAT | |
| Dp1954rt_F1i | TTTGAGTACCGCGAGAAGGA | qPCR detection of DPH1 |
| Dp1954rt_R1i | TCGTTGACCACCGAGATGT | |
| Dp1955rt_F1i | ATGACTACCCTTCGGCTGTC | qPCR detection of DPH2 |
| Dp1955rt_R1i | CACGTCGAAGAGGAACACG | |
| Dp1956rt_F1i | GGAGGGCCAGATCATCATAGG | qPCR detection of DPH3 |
| Dp1956rt_R1i | TGTCGAGGATGTGGAAGACG | |
| Dp1957rt_F1i | GATGAGCCTCGAGGAGCTG | qPCR detection of DPH4 |
| Dp1957rt_R1i | GAGAGCAGGTGGTACGTCA | |
| Dp1958rt_F1i | CAGGGAGAACATCCAGCAGA | qPCR detection of DPH5 |
| Dp1958rt_R1i | GGGACGTAGAACAGACTTCCA | |
| Dp1959rt_F1i | CGTTCTCGTCATCACTGTCG | qPCR detection of DPH6 |
| Dp1959rt_R1i | TCGGTGATTCTAGGGATGGC | |

[a] An overhang sequence in each primer for overlap with sequences at the ends of the linearized plasmid, *gen/gfp* cassettes, or genes is underlined.
